# A Small-Molecule Approach to Bypass In Vitro Selection of New Aptamers: Designer Pre-Ligands Turn Baby Spinach into Sensors for Reactive Inorganic Targets

**DOI:** 10.1101/2023.07.29.551132

**Authors:** Tushar Aggarwal, Liming Wang, Bryan Gutierrez, Hakan Guven, Huseyin Erguven, Enver Cagri Izgu

**Author notes:** Corresponding authors: Izgu, E. C.

## Abstract

Fluorescent light-up aptamer (FLAP) systems are promising biosensing platforms that can be genetically encoded. Here, we describe how a single FLAP that works with specific organic ligands can detect multiple, structurally unique, non-fluorogenic, and reactive inorganic targets. We developed 4-*O*-functionalized benzylidene imidazolinones as pre-ligands with suppressed fluorescent binding interactions with the RNA aptamer Baby Spinach. Inorganic targets, hydrogen sulfide (H_2_S) or hydrogen peroxide (H_2_O_2_), can specifically convert these pre-ligands into the native benzylidene imidazolinones, and thus be detected with Baby Spinach. Adaptation of this approach to live cells opened a new opportunity for top-down construction of whole-cell sensors: *Escherichia coli* transformed with a Baby Spinach-encoding plasmid and incubated with pre-ligands generated fluorescence in response to exogenous H_2_S or H_2_O_2_. Our approach eliminates the requirement of in vitro selection of a new aptamer sequence for molecular target detection, allows for the detection of short-lived targets, thereby advancing FLAP systems beyond their current capabilities. Leveraging the functional group reactivity of small molecules can lead to cell-based sensors for inorganic molecular targets, exploiting a new synergism between synthetic organic chemistry and synthetic biology.

## INTRODUCTION

Aptamers are structured nucleic acid sequences that selectively bind a molecular target with high affinity. Aptamer sequences can be found in nature as gene regulation elements located in the untranslated regions of mRNAs,^1^ or discovered in the laboratory by in vitro selection (or systematic evolution of ligands by exponential enrichment, called SELEX) from a random pool of nucleic acids.^2,3^ Some in vitro-selected aptamers can bind and increase the fluorescence quantum yield (<λ_F_) of fluorogenic small-molecule ligands that are otherwise poorly fluorescent in their unbound state.^4^ These aptamers, often referred to as fluorescent light-up aptamers (FLAPs), can serve as genetically encodable and non-invasive (bio)sensing platforms and have attracted significant attention across a number of areas, including environmental monitoring,^5^ forensics,^6^ bioanalyte detection and clinical diagnostics,^7–11^ live cell imaging,^12–16^ and synthetic biology.^17,18^

Currently, there exist fundamental challenges that limit the broader potential of FLAP-based sensing platforms. A FLAP consensus sequence that binds a particular organic molecule has no direct utility for detecting an inorganic target. Further, the aptamer ligand must have inherent fluorogenicity to generate signal in a FLAP system, which renders non-fluorescent chemical cues or biomarkers undetectable. Recent aptamer-based approaches for sensing non-fluorescent targets have employed a secondary aptamer domain that binds the target to induce folding or conformational stabilization of a tethered FLAP.^19,12,20^ Though elegant in design, the target-binding aptamer domain must be in vitro selected or engineered for each new target of interest. Additionally, programming complex RNA networks to correctly and conditionally interact can be difficult due, in part, to the highly dynamic interactions among long RNA transcripts^21,22^ and inherent limitations in RNA stability or folding under the suboptimal physiological conditions.^23^ Finally, the aptamer-ligand binding is an equilibrium process that relies on the availability of free ligand.^24^ As a result, it would be challenging to realize an aptamer system that can bind targets with limited biological lifetime.

Here we describe a structure- and reactivity-guided small-molecule approach that can help address these fundamental challenges. This approach allows for a single aptamer, known to bind a specific class of organic fluorogens, to detect multiple, structurally unique, non-fluorogenic, and reactive inorganic targets (Figure 1). The RNA aptamer Spinach binds 4-hydroxybenzylidene imidazolinone (HBI) and its halogenated analogs with a *K*_d_ in nanomolar-to-micromolar ranges.^25–28^ Guided by molecular docking simulations and functional group reactivity principles in synthetic organic chemistry, we developed 4-*O*-functionalized benzylidene imidazolinone architectures that exhibit weak or inhibited fluorescent binding toward Baby Spinach. This aptamer (51-nt) is a truncated version of Spinach2 (95-nt),^29^ which is the structurally more stable version of Spinach (97-nt).^27^ These HBI-derived small molecules, here referred to as “pre-ligands”, can be converted into native (4-*O*-free) HBIs by an inorganic target. In this study, we focused on redox-active inorganics with major biological roles: hydrogen sulfide (H_2_S) and hydrogen peroxide (H_2_O_2_). H_2_S is a reducing agent that has cytoprotective properties against oxidative stress^30^ and is involved in signal transduction pathways pertaining to neurological^31–34^ and cardiovascular^35,36^ processes. H_2_S can diffuse through lipid membranes without specialized channels.^37^ H_2_O_2_ is an oxidant^38^ involved in a wide array of biological processes, including signal transduction,^39,40^ cell differentiation and proliferation,^41^ immune response,^42^ mitochondrial dysfunction,^40^ and oncogene activity.^41,43^ H_2_O_2_ transport across membranes occurs through simple diffusion or is facilitated by aquaporins in aerobic organisms.^44,45^

**Figure 1.**
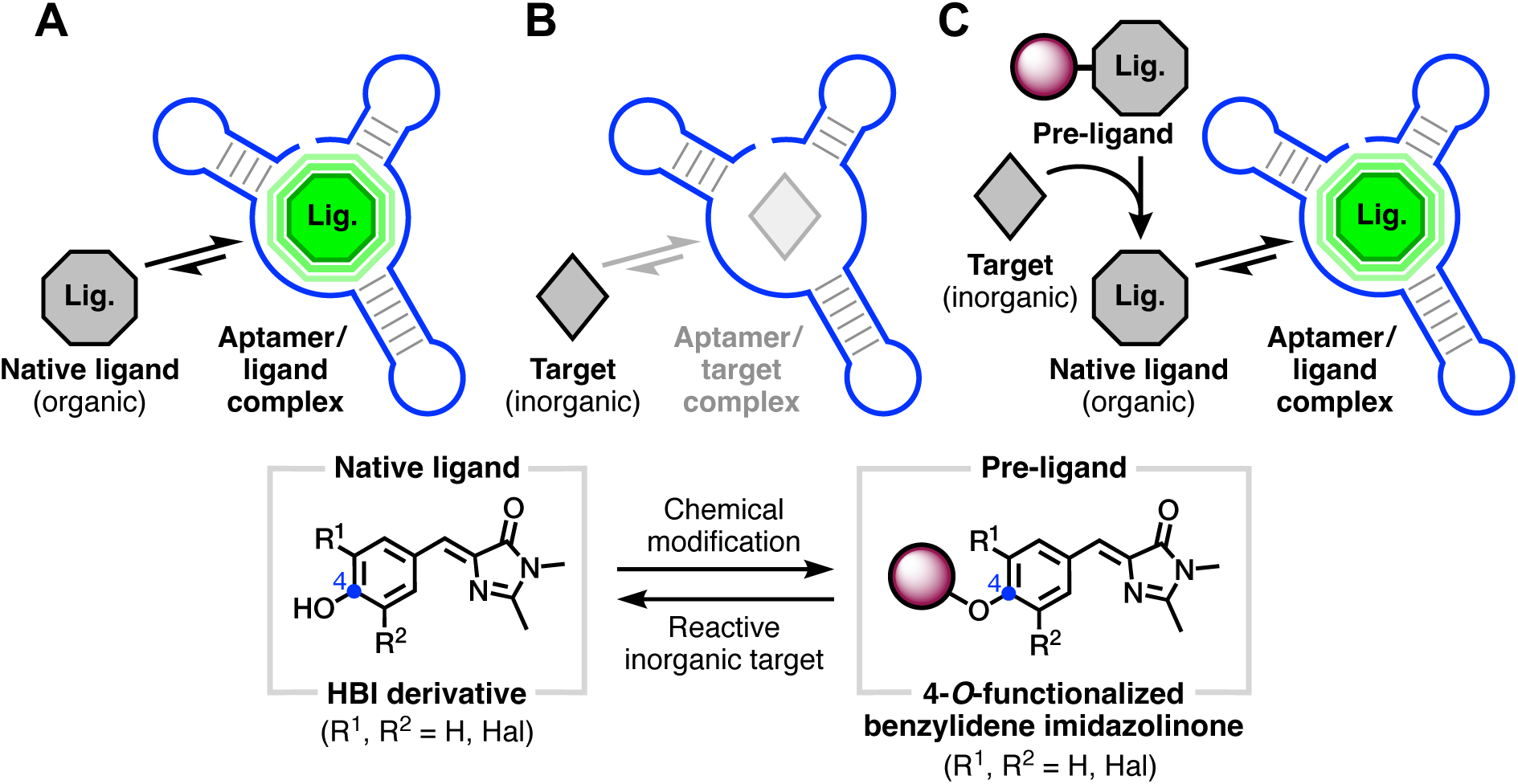
Conceptual illustration of how a FLAP system can be repurposed to detect an inorganic molecular target without aptamer engineering. **(A)** Strong fluorescence induced by non-covalent binding interactions between an aptamer and its native, inherently fluorogenic ligand (e.g., HBI derivative). **(B)** Inorganic targets, like majority of biologically relevant molecules, may lack inherent fluorogenicity, affinity for a structured nucleic acid sequence, or both. **(C)** If exists, the chemical reactivity of the inorganic target can be utilized to conditionally form the native ligand in situ from a pre-ligand design with inhibited fluorogenic aptamer affinity (e.g., 4-*O*-functionalized benzylidene imidazolinone). This would allow for the detection of inorganic targets by a known aptamer, thereby expanding the application space of structured nucleic acids without in vitro selection or additional oligonucleotide scaffold engineering.

The FLAP-mediated activity-based sensing method using Baby Spinach and pre-ligands can detect H_2_S or H_2_O_2_ with good selectivity against other redox species. Adaptation of this method to live cells through plasmid transfection opened a new opportunity for building whole-cell biosensors: *Escherichia coli* competent cells (BL21-DE3) transformed with a Baby Spinach-encoding plasmid and incubated with the select pre-ligand generated fluorescence signal in response to H_2_S or H_2_O_2_. Whether in cell-free conditions or with *E*. *coli*, utilizing the FLAP and pre-ligand makes it possible to detect chemical species that are traditionally not considered as aptamer ligands due to their inherent reactivity and short life. Further, it eliminates the need of in vitro selection (or SELEX) of a new aptamer consensus sequence, thereby expanding the current capabilities of nucleic acid-based detection. Our method leverages functional group reactivity of small molecules to build in situ cell-based sensors for chemically distinct and reactive molecular targets, which exploits a previously overlooked synergism between synthetic organic chemistry and synthetic biology.

## RESULTS AND DISCUSSION

### Docking calculations aid in experimental interpretation and guide pre-ligand design

To structurally rationalize the effect of ligand derivatization on Spinach binding, we performed molecular docking (Figures 2 and S1) using Auto-dock Vina 1.2.0,^46,47^ which presents a success rate of >90% in detecting receptor-ligand bindings with an average RMSD of <2.0 Å.^48^ Recognizing the limitation of Auto-dock Vina for heteroatoms, we used methoxycarbonyl, phenyl, benzyl, and benzoyl substituents to simply mimic potential 4-*O* caging groups. Our initial simulations used DFHBI (Figures 2) due to the availability of Spinach/DFHBI co-crystal structure.^27^ Docking results with the other benzylidene architectures are presented in Figure S1. Based on the crystal structure,^27^ DFHBI forms key hydrogen bond interactions with G26/G31/A64 nucleobases of Spinach (Figure 2A). These interactions were largely retained with similar bond distances in our docking simulation with DFHBI (Figure 2B, docking score of −8.9 kcal mol^−1^), validating that the native ligand-aptamer binding interactions can be attained *in silico* using AutoDock Vina. Strikingly, when the 4-*O* atom of DFHBI was connected to methoxycarbonyl, phenyl, benzyl, or benzoyl group, the resulting BI derivative bound G26/G31/A64 pocket with a near 180° rotation, positioning either the 4-*O*-benzylidene motif opposite to G26 (for methylcarbonyl-DFHBI, benzoyl-DFHBI, and benzyl DFHBI) or the imidazolinone carbonyl opposite to G31 (for phenyl-DFHBI). For all these benzylidene imidazolinones, the native H-bonding interactions within the G26/G31/A64 pocket were abolished and the docking scores (methylcarbonyl-DFHBI: −5.6, phenyl-DFHBI: 2.7, benzoyl-DFHBI: −5.6, and Benzyl-DFHBI: −6.6 kcal mol^−1^) indicated significantly weaker binding affinity. Taken together, the docking results provided support for a scenario in which the positioning of the benzylidene imidazolinone motif within the G26/G31/A64 binding pocket of Spinach can be altered by modifying the 4-*O*-benzylidene atom with a range of functional groups.

**Figure 2.**
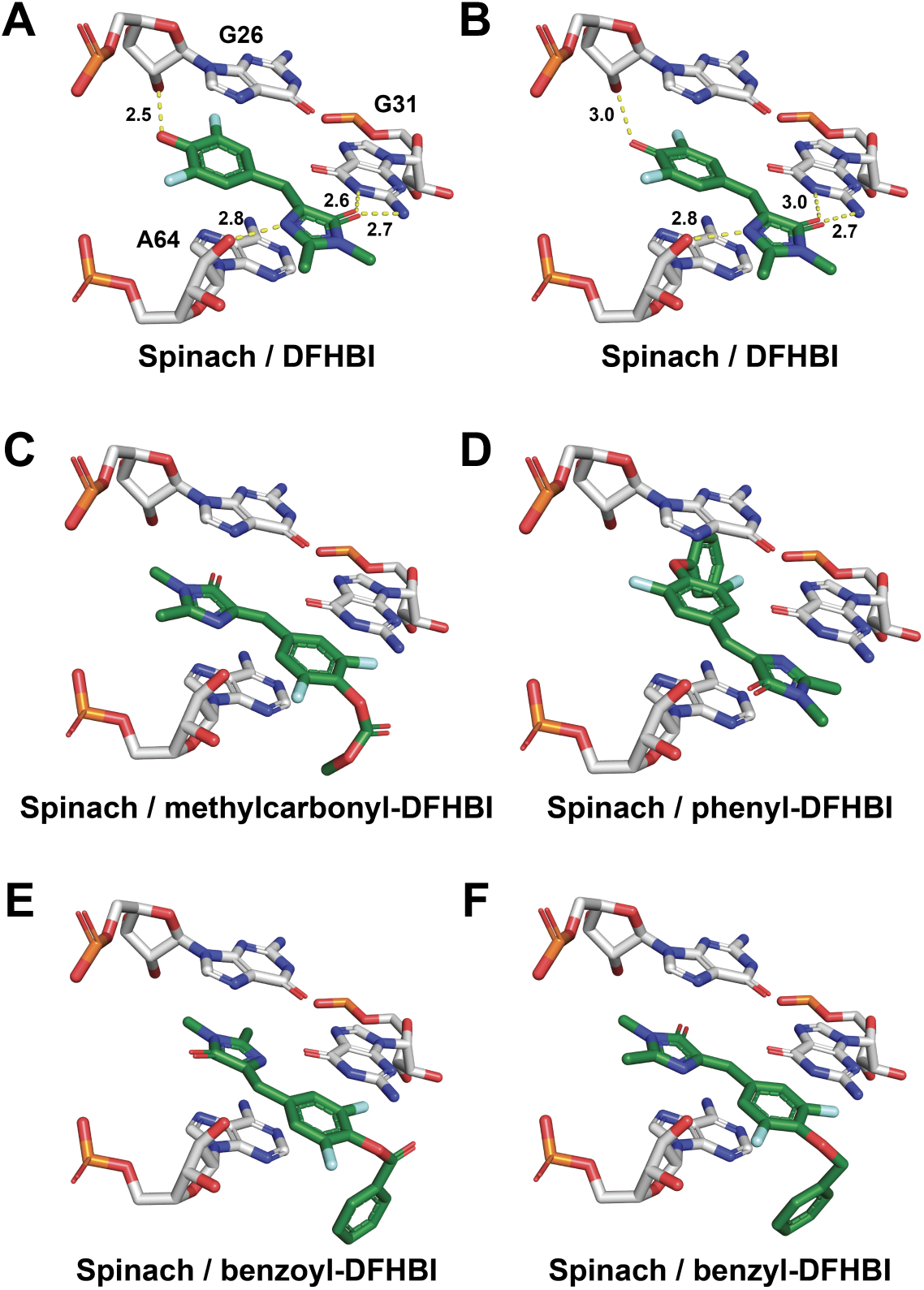
Docked poses: binding mode of the selected 4-*O*-aryl-modified small molecule derivatives of DFHBI. **(A)** Model of key hydrogen-bonding interactions within the crystal structure of Spinach-DFHBI (PDB:4TS2) near the binding site. **(B)** binding mode of the DFHBI ligand prepared in silico and docked using AutoDock Vina. **(C-F)** docked poses of the 4-*O*-aryl-modified small molecule derivatives of DFHBI.

### Building the Baby Spinach pre-ligand library

Redox-mediated chemical transformations of functional groups have been successfully implemented to release phenols.^49–52^ Inspired by these transformations and guided by our molecular docking simulations, we developed a library of small molecules to serve as Baby Spinach pre-ligands (Figure 3). This library included compounds synthesized from various HBI fluorogens (see supplementary information for synthesis details) and designed to react with either H_2_S (Figure 3A, left panel) or H_2_O_2_ (Figure 3A, right panel). The H_2_S-targeting designs included **<u>a</u>**zido**<u>e</u>**thyl**<u>c</u>**arbonyl-modified MFHBI (AEC-MFHBI) and DFHBI (AEC-DFHBI); **<u>d</u>**<u>i**n**</u>itro**<u>p</u>**henyl-modified MFHBI (DNP-MFHBI); **<u>p</u>**yridyl**<u>d</u>**<u>i**s**</u>ulfunyl**<u>b</u>**<u>en**z**oyl</u>-modified MFHBI (PyDSBz-MFHBI) and DBrHBI (PyDSBz-DBrHBI). The H_2_O_2_-targeting designs included 4-(**<u>m</u>**ethyl)**<u>b</u>**enzene**<u>b</u>**oronic **<u>a</u>**cid **<u>p</u>**inacol **<u>e</u>**ster-modified MFHBI (MBBAPE-MFHBI) and DFHBI (MBBAPE-DFHBI); and 4-(**<u>m</u>**ethyl)**<u>b</u>**enzene**<u>b</u>**oronic **<u>a</u>**cid-modified MFHBI (MBBA-MFHBI).

**Figure 3.**
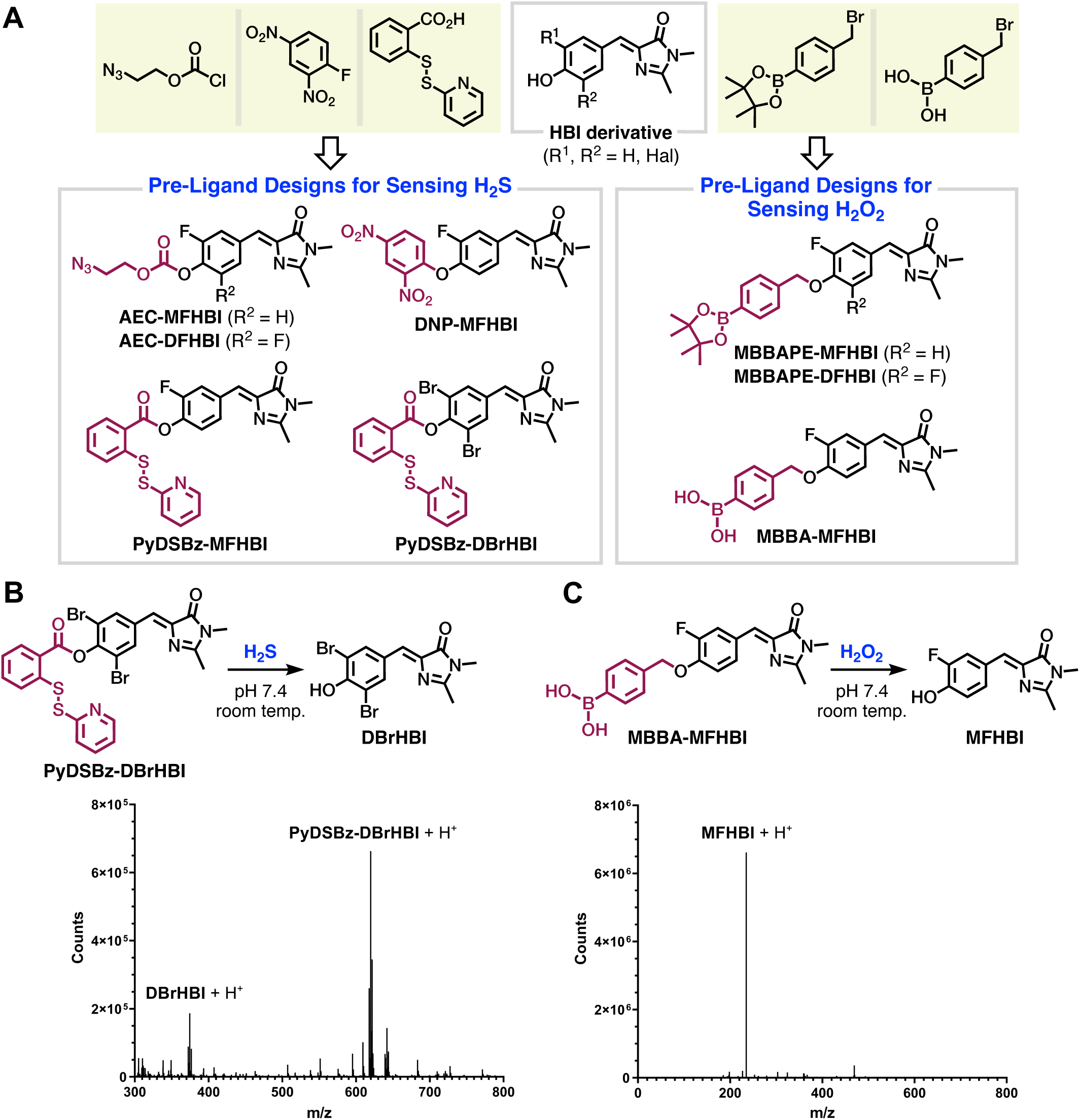
Baby Spinach pre-ligand designs and native ligand formation. **(A)** Pre-ligands synthesized from respective building blocks (yellow panels) for detecting (left) H_2_S or (right) H_2_O_2_. **(B**, top) H_2_S-induced generation of DBrHBI from PyDSBz-DBrHBI. (Bottom) Overlay of extracted mass spectrum acquired following liquid chromatography. For PyDSBz-DBrHBI, [M+H]^+^ requires 619.9131; found 619.9116; for DBrHBI, [M+H]^+^ requires 374.9162; found 374.9198. **(C**, top) H_2_O_2_-induced generation of MFHBI from MBBA-MFHBI. (Bottom) Spectrum acquired via direct injection. For MFHBI, [M+H]^+^ requires 235.0878; found 235.0922. No significant peak related to MBBA-MFHBI was observed. **(B** and **C**, bottom) Crude reaction aliquot was taken after 1 hour of mixing pre-ligand with the respective redox agent and analyzed in positive ionization mode. Peak heights are not an accurate representation of relative concentrations.

Regarding the pre-ligands designed to react with H_2_S, both AEC-MFHBI and AEC-DFHBI displayed poor selectivity against biologically relevant reactive sulfur species (RSS) other than H_2_S, such as cysteine (Cys) and glutathione (GSH). We hypothesized that the conversion of these pre-ligands to MFHBI or DFHBI by Cys and GSH is through nucleophilic attack of thiol sulfur to the benzylidene carbonate. To test this hypothesis, we synthesized **<u>m</u>**ethyl**<u>c</u>**arbonyl-MFHBI (MC-MFHBI) and subjected the mixtures of MC-MFHBI and Baby Spinach to Cys, GSH, and H_2_S separately (Figure S2). Signal-to-background ratios (*F/F_0_*s) were then calculated, where *F* and *F_0_* are the fluorescence intensities for samples with and without the redox species, respectively. Addition of Cys and GSH led to *F/F_0_* values of 18.5 and 3.5, which are comparable to that of AEC-MFHBI (10.9 and 6.3), while H_2_S treatment displayed less than 30% fluorescence increase after 2 hours of incubation. These results indicate that 1) H_2_S has weaker nucleophilic reactivity toward the benzylidene 4-*O*-carbonate than Cys and GSH, supporting previous reports,^53^ and 2) the conjugate bases of MFHBI and DFHBI can serve as good leaving groups. Therefore, we reasoned that limiting the steric access to the benzylidene 4-*O*-carbonyl site would largely evade nucleophilic thiol attack, thereby preventing non-specific signal generation while favoring elimination. Our reasoning is mechanistically supported by the previous observations that sterically hindered esters show increased stability against hydrolysis while favoring self-immolation.^54^ Further, in the presence of Baby Spinach, both DNP-MFHBI and PyDSBz-MFHBI provided similar *F*/*F_0_* values for Cys and GSH. We presumed that the dinitrobenzyl moiety in DNP-MFHBI and the ester in PyDSBz-MFHBI may readily react with thiols; thus, incorporation of the second fluorine atom on the benzylidene ring would render these functional groups even more reactive. Therefore, we excluded the synthesis of their DFHBI analogs. Among the pre-ligands we have explored, PyDSBz-DBrHBI possesses a sterically hindered benzylidene 4-*O*-carbonyl, which in turn would increase its selectivity toward H_2_S. For the pre-ligands designed to react with H_2_O_2_, both MBBAPE-MFHBI and MBBAPE-DFHBI displayed poor stability in near-neutral buffer and in the absence of H_2_O_2_. We observed their conversion to MFHBI and DFHBI by high-resolution mass spectrometry (HRMS), respectively, and their mixtures with Baby Spinach exhibited spontaneously increasing levels of fluorescence, suggesting that these DFHBI-derived compounds are too reactive to serve as pre-ligands. In contrast, the monofluoro analog, MBBA-MFHBI, is sufficiently stable in buffer. MBBA-DFHBI was not included into the library due to unresolved difficulties in its synthesis and isolation. In addition to these observations on selectivity and stability, our preliminary findings with the HBI derivatives indicated that the aqueous solubility of boronic acid-caged pre-ligand is superior to its boronic acid pinacol ester counterpart. All the pre-ligand designs we studied here, except for the MBBAPE analogs, are soluble in aqueous buffer (1% v/v DMSO, pH 7.4) with concentrations reaching at least 100 µM. Based on all these evaluations, we therefore elected PyDSBz-DBrHBI and MBBA-MFHBI as pre-ligands to react with H_2_S and H_2_O_2_, respectively.

Conversions of PyDSBz-DBrHBI and MBBA-MFHBI to their respective redox products DBrHBI and MFHBI were evaluated through HRMS (Figures 3B and 3C). For each pre-ligand design, the crude reaction aliquot taken after 1 hour of mixing with the respective redox agent in physiologically relevant conditions. The spectral data indicated the presence of native ligand, DBrHBI or MFHBI, which validated the reactivity of benzylidene 4-*O*-modifications. Furthermore, we determined the benzylidene 4-O*H* p*K*_a_ of DBrHBI as 5.2 (Figure S3) and of MFHBI as 6.8 (Figure S4), indicating that the native ligands to be generated from PyDSBz-DBrHBI and MBBA-MFHBI should exist primarily in their fluorescently enhanced anionic forms at physiological pH.

### Binding affinities of pre-ligands characterized by fluorescence quenching

With the chemically feasible Baby Spinach pre-ligands identified, we conducted a fluorescence quenching assay to gain an understanding of the relative fluorescent binding affinities of these pre-ligands compared to their native counterparts (Figure 4A). To this end, we incubated mixtures of Baby Spinach (1 µM) and its native ligand (2 µM) with varied concentrations of the pre-ligand each and recorded their fluorescence intensities after 1 hour of incubation. The sample containing 50 µM of PyDSBz-DBrHBI exhibited a fluorescence intensity similar to that of 0 µM PyDSBz-DBrHBI, representing minimal competition between PyDSBz-DBrHBI and DBrHBI in binding Baby Spinach (Figure 4B). Incubation of the mixture of Baby Spinach and MFHBI with 50 µM of MBBA-MFHBI, which is 25 equivalents of MFHBI, retained fluorescence intensity at ∼70% (Figure 4C). This observation indicates that the binding of MBBA-MFHBI to Baby Spinach may occur but is substantially weaker than the binding between MFHBI and Baby Spinach. These results correlate well with the predicted disfavored binding of pre-ligands to Spinach from our docking calculations (*cf*. Figures 2E-F and S1).

**Figure 4.**
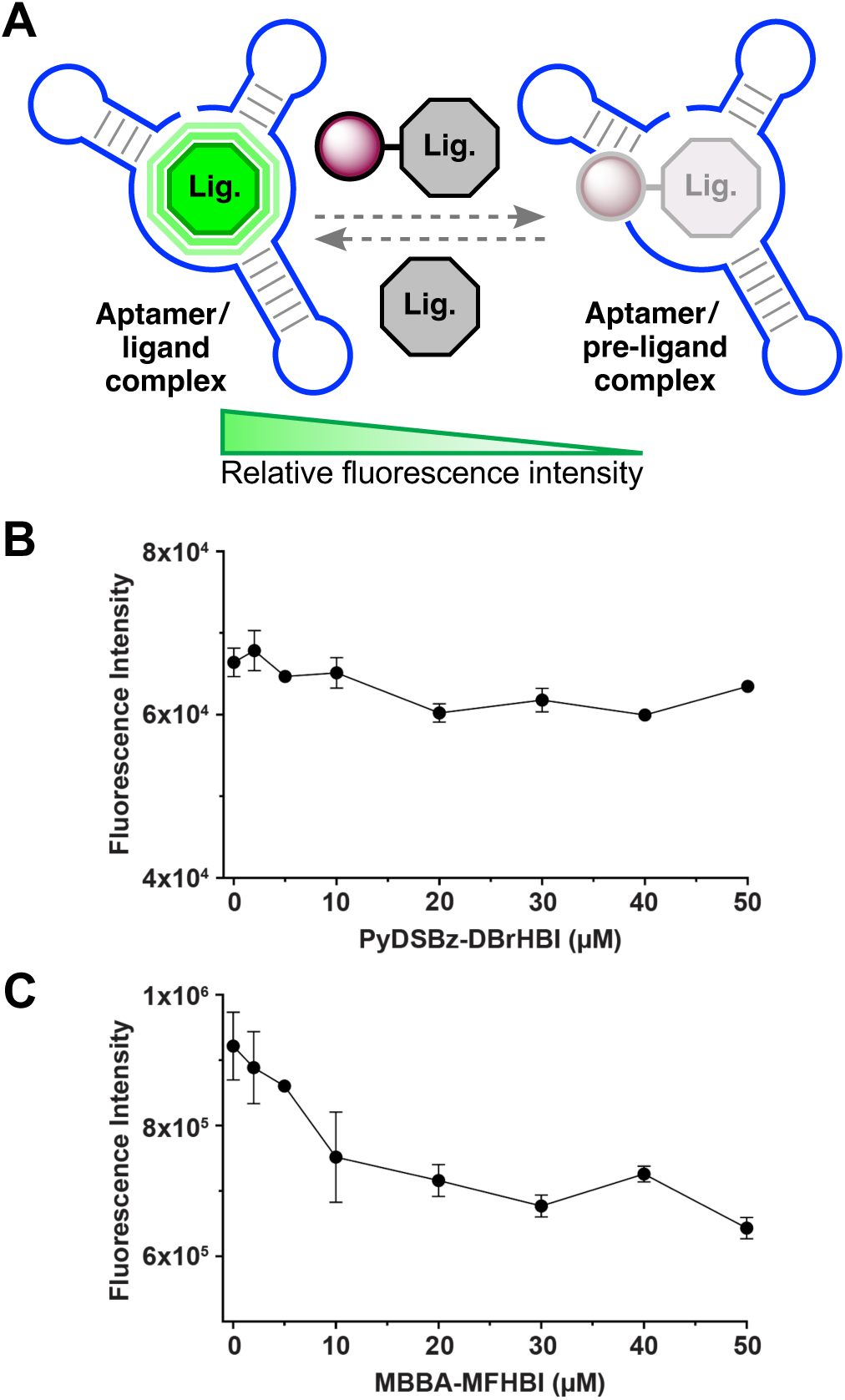
Fluorescence quenching with pre-ligands. **(A)** Aptamer affinities of pre-ligands were assessed via competitive ligand binding assay. Fluorescence intensities of **(B)** Baby Spinach/DBrHBI complex vs PyDSBz-DBrHBI concentration, and **(C)** Baby Spinach/MFHBI complex vs MBBA-MFHBI concentration. Error bars represent standard deviation, *n* = 3.

### Cell-free redox sensing with Baby Spinach and pre-ligands

The fluorescent signal generation rates, magnitudes, and selectivities of PyDSBz-DBrHBI and MBBA-MFHBI were evaluated via microplate reader measurements (Figure 5). The *F/F_0_* values for 60-minute incubation of the selected pre-ligand (5 µM) with Baby Spinach (1 µM) were calculated. Mammalian cells have been shown to respond to 10–600 µM H_2_S as an exogenous source,^55^ therefore, we reasoned that addition of Na_2_S at 500 µM into a buffer at pH 7.4 can serve as a proper mimic for biological H_2_S environment. At neutral pH, around 20% of its sulfur containing forms should exist as H_2_S (∼100 µM), based on the p*K*_a_ (∼6.9) of NaSH.^56^ Results indicated that PyDSBz-DBrHBI achieved an *F/F_0_* of 37.3 with 500 µM Na_2_S as the source of H_2_S, indicating an increase in fluorescence ∼3600%. We also recorded the fluorescence intensities every 10 minutes over a 60-minute incubation period and compared them with those of Baby Spinach with DBrHBI (Figure 5A). Fluorescence intensities of samples containing PyDSBz-DBrHBI plateaued at ∼50 minutes and reached up to ∼30% of that with DBrHBI. To assess the performance of the H_2_O_2_-responsive pre-ligand, MBBA-MFHBI, we monitored the relative fluorescence intensity change of Baby Spinach solution containing either MFHBI or MBBA-MFHBI in presence of H_2_O_2_ over 60 minutes (Figure 5D). After the addition of H_2_O_2_ (estimated concentration of 100 µM upon mixing) to the solution containing MBBA-MFHBI, we observed ∼2500% increase in fluorescence intensity (*F/F_0_* = 25.2) compared to only 28% increase without H_2_O_2_ addition, suggesting that phenyl boronic acid group of MBBA-MFHBI is stable at physiological pH and can generate free MFHBI upon reacting with H_2_O_2_. Fluorescence intensities of MBBA-MFHBI in the presence of H_2_O_2_ reached up to ∼75% that of MFHBI within 1 hour.

**Figure 5.**
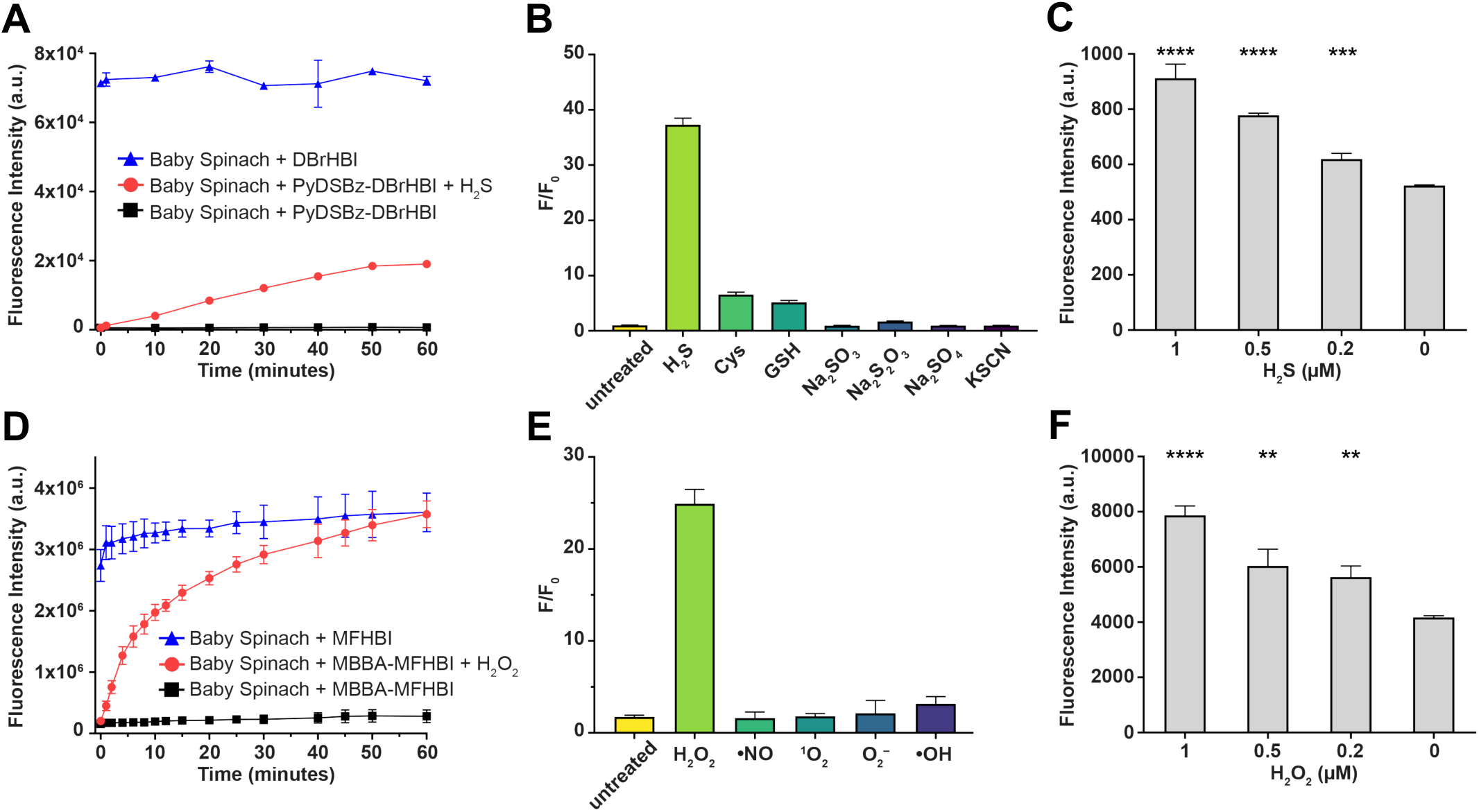
Performances of PyDSBz-DBrHBI and MBBA-MFHBI with Baby Spinach in cell-free conditions. **(A and D)** Time-dependent fluorescence increase of pre-ligands upon addition of their respective redox agents (Na_2_S at 500 µM, H_2_O_2_ at 100 µM). **(B and E)** The fluorescence fold increase (*F*/*F_0_*) with the addition of RSS (500 µM) and ROS (100 µM), respectively. **(C and F)** LoD measurements for PyDSBz-DBrHBI and MBBA-MFHBI, respectively. Error bars represent standard deviation based on triplicates. Single-tailed Student’s *t* test: *, P <0.05; **, P <0.01; ***, P <0.001; ****, P <0.0001. Time-dependent change in fluorescence intensity upon addition of H_2_S or H_2_O_2_. DBrHBI, PyDSBz-DBrHBI, MFHBI, and MBBA-MFHBI (5 µM), Baby Spinach (1 µM), HEPES (pH 7.4, 50 mM), KCl (100 mM), MgCl_2_ (10 mM), and Na_2_S (500 µM) or H_2_O_2_ (100 µM). Error bars represent standard deviation based on triplicates. Bar graph: Signal turn on (*F*/*F_0_*) 60 minutes after addition of analyte.

We then evaluated the redox-specificities of the aptamer pre-ligands PyDSBz-DBrHBI (Figures 5B) and MBBA-MFHBI (Figures 5E). The fluorescence intensities were recorded after incubating a mixture of the pre-ligand and Baby Spinach with a redox species for 1 hour, and *F/F_0_* were calculated, where *F* and *F_0_* are the fluorescence intensities for samples with and without the redox species, respectively. The *F/F_0_* value for the control samples, which included only the pre-ligand and Baby Spinach in buffer, were set to 1 for comparison. For a mixture of PyDSBz-DBrHBI and Baby Spinach was incubated with Na_2_S (500 µM) for 1 hour, *F/F_0_* was measured as 37.3. Of significant note, when the same mixture was incubated with cysteine and glutathione, which are nucleophilic thiols abundant in biological environments, *F/F_0_* reached just 6.6 and 3.9, respectively. Other RSS we tested (Cys, GSH, SO_3_^2–^, S_2_O_3_^2–^, SO_4_^2−^, and SCN^−^) generated *F/F_0_* values centered around ∼1, indicating no measurable pre-ligand activation. As for the samples containing MBBA-MFHBI, *F/F_0_* was 25.2 with H_2_O_2_ after 1 hour of incubation. In contrast, *F/F_0_* values were <4 for other reactive oxygen species (ROS), (•NO, ^1^O_2_, O_2_^−^•, •OH), suggesting that MBBA-MFHBI is selective for H_2_O_2_.

Limit of detection (LoD) values were determined for pre-ligands PyDSBz-DBrHBI and MBBA-MFHBI by evaluating the statistical difference between *F* and *F_0_* measured with Baby Spinach at different concentrations of H_2_S and H_2_O_2_, respectively (Figures 5C and 5F). Here, LoD is defined as the minimum redox agent concentration at which *F* is statistically higher than *F_0_* (*P*>0.05) according to one-tailed Student’s *t* test. We determined LoD for PyDSBz-DBrHBI as 0.2 µM of H_2_S and that for MBBA-MFHBI as 0.2 µM of H_2_O_2_. The LoD of PyDSBz-DBrHBI determined is close to other recently reported fluorescence-based H_2_S sensing systems.^57–59^ The sensitivity of MBBA-MFHBI is comparable to fluorescence-based H_2_O_2_ sensors that use small-molecule fluorophore activation.^60,61^

### Turning E. coli into cell-based sensors using pET-21-3xBaby Spinach and pre-ligands

Bacteria, notably *E. coli*, are convenient and robust model organisms widely employed for top-down construction of whole-cell biosensors.^62^ In accordance, we focused on building chemically competent *E. coli* BL21 Star (DE3) cells as redox sensors, and initially transformed them with a DNA plasmid containing tRNA-Spinach2 construct.^7^ Confocal imaging in the presence of MFHBI (preliminary positive control) exhibited weak intracellular brightness following 488 nm excitation. Therefore, we designed and cloned a gene encoding for tRNA fused with three continual Baby Spinach consensus regions, each connected through a 6-bp sealing stem, named pET-21-3xBaby Spinach (Figures 6A, S5, and S6). We reasoned that *E. coli* transcribing this pET-21-3xBaby Spinach construct would provide up to three ligand binding domains per transcript compared to tRNA-Spinach2 with a single domain, thus generate stronger fluorescence in the presence of the native ligands. Indeed, we observed bright signals in samples incubated with DBrHBI (440 nm excitation, Figure 6B top-left, Figure 6C DBrHBI) or MFHBI (488 nm excitation, Figure 6D top-left, Figure 6E MFHBI). This observation supported the notion that RNA transcripts with multiple ligand-binding FLAP domains could amplify the fluorescence output, which is also in agreement with a previous report demonstrating an increase in brightness of Spinach-tagged guide RNA in CRISPR-Display.^63^ Consequently, for the subsequent experiments, *E. coli* carrying pET-21-3xBaby Spinach plasmid were grown in LB at 37 °C. RNA transcription was induced by the addition of 1 mM isopropyl β-D-1-thiogalactopyranoside (IPTG) once OD (600) reached 0.4, and cells were grown for an additional 2 hours to further facilitate aptamer transcription. Harvested cells were incubated with M9 media containing either PyDSBz-DBrHBI (100 µM) or MBBA-MFHBI (100 µM) for 15 minutes. The media was replaced with fresh M9 media containing either H_2_O_2_ (100 µM) or Na_2_S (2 mM). This media replacement ensured the removal of pre-ligand molecules not internalized by the cells. Although we have no direct measurement of intracellular concentrations of pre-ligands, we postulate that pre-ligands can be internalized by *E*. *coli* through passive diffusion while some may be secreted out through efflux. The signal generation could result from either passive diffusion of pre-ligands into *E*. *coli* followed by intracellular redox reaction or extracellular redox reaction of effluxed pre-ligands and subsequent cellular uptake. The latter mechanism, however, would constitute a major decrease in signal generation due to substantial dilution of pre-ligands when effluxed to the newly replaced media. Aliquots (1 mL) of *E. coli.* culture were used at different time points for confocal imaging and a red membrane dye (Fm 4-64FX) was used as a co-stain to label the *E. coli* membrane. It is worth noting that the use of Fm 4-64FX allowed us to observe heterogeneity in the overall population for aptamer transcription levels, as the positive controls displayed different levels of brightness following excitation at either 440 nm or 488 nm (Figures 6B or 6D, top-left). Furthermore, it allowed for visualization of cell morphology from the time of pre-ligand addition until the end of redox incubation time. The observation that cells retained their morphology (rod shaped) indicated that *E*. *coli* tolerates the concentrations we used for pre-ligands (100 µM) and redox species (100 µM H_2_O_2_ or 2 mM Na_2_S). Samples without DBrHBI (Figure 6B, top-middle) or MFHBI (Figure 6D, top-middle) served as a measure of background fluorescence while providing the microscopy parameters for subsequent confocal images and data analyses. *E. coli.* carrying the plasmid pUC19 (negative control), lacking 3xBaby Spinach aptamer gene, exhibited no detectable fluorescence under 440 or 488 nm excitation in the presence of DBrHBI (Figure 6B, top-right) or MFHBI (Figure 6D, top-right), respectively.

**Figure 6.**
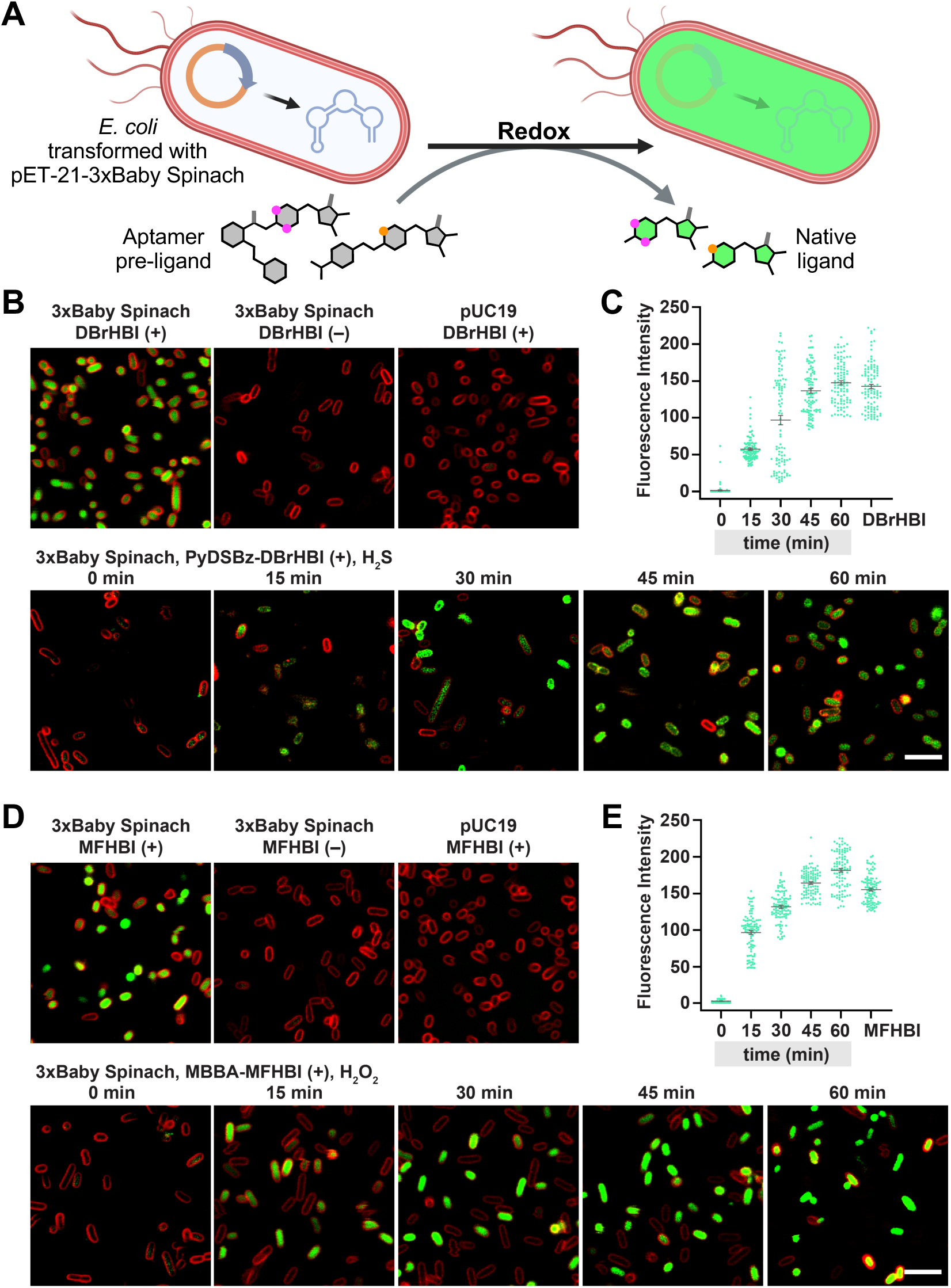
Turning *E*. *coli* into redox sensors. **(A)** When transformed with pET-21-3xBaby Spinach and incubated with a Baby Spinach pre-ligand, BL21 Star (DE3) *E. coli* can fluoresce in response to a redox cue. **(B**, **D)** Confocal microscopy images of live *E. coli* sensing **(B)** H_2_S or **(D)** H_2_O_2_. Top row panels represent positive and negative controls, including transfection with pUC19 plasmid, which contains no aptamer gene. Bottom row panels represent redox sensing at 0, 15, 30, 45, and 60 minutes. **(C, E)** Quantitative assessment of fluorescence outputs from cells over 60 minutes, along with the positive control measured at 60-minute time point using either **(C)** DBrHBI or (**E)** MFHBI. Cells were incubated in M9 media containing 100 µM of the HBI derivative. Redox species introduced to the imaging media: **(B, C)** Na_2_S (2 mM), **(D, E)** H_2_O_2_ (100 µM). For co-staining, cells were treated with a FM 4-64FX (red membrane dye, 1 µg/ml, 2 minutes). Channel/excitation (nm): DBrHBI/440, MFHBI/488, Fm4-64FX/568. Scale bar: 5 µm.

We obtained the fluorescence intensity vs time plots (Figure 6C and 6E) from the confocal microscopy data by quantifying the mean fluorescence of each cell imaged at time intervals 0–60 minutes following 440 or 488 nm excitation (see Supporting Information for detailed procedure). This analysis allowed us to gain a quantitative insight into the aptamer-dependent fluorescence enhancement in *E. coli*. For both reductive (H_2_S) and oxidative (H_2_O_2_) sensing, cells displayed statistically significant fluorescence enhancement over 60 minutes of incubation. The degree of heterogeneity in fluorescence intensity, especially for 15- and 30-minute samples, supported the visual observation of increase in brightness over time. Samples incubated with PyDSBz-DBrHBI (100 µM) and then exposed to exogenous Na_2_S (2 mM) showed a progressive increase in cell populations with detectable intracellular fluorescence over the course of imaging (Figures 6B bottom and 6C). At 60 minutes following H_2_S treatment, the mean fluorescence intensity value was determined to be 105-fold higher than that at 0 minute (147.6 vs 1.4). The single cell fluorescence outputs were detected within 15 minutes upon addition of Na_2_S, which then plateaued over 60 minutes (Figure 6C). We observed similar trends in *E. coli* incubated with MBBA-MFHBI (100 µM) and then exposed to H_2_O_2_ (100 µM) (Figures 6D bottom and 6E). The mean fluorescence intensity increased from 2.6 a.u. at 0 minute to 181.7 a.u. at 60 minutes following H_2_O_2_ addition. The single cell fluorescence intensities reached to a maximum (centered around 200 a.u.) within 30 minutes, while the overall mean fluorescence of the *E. coli* populations progressively increased over 60 minutes.

## CONCLUSION

Interest in fluorescent light-up aptamer (FLAP)-based target detection is substantial given the ever-evolving needs in analyte biosensing, diagnostics, and environmental surveillance. The traditional target recognition, whether directly by the FLAP itself or through its conditional folding induced by a secondary aptamer domain,^19^ entails the availability of a distinct aptamer consensus sequence for each new and distinct target. Even though these approaches find biological use, there has been no report on the design of a single aptamer that can be repurposed for more than one structurally and chemically unique targets, especially reactive inorganic molecules. This study describes how structure- and reactivity-guided design principles in synthetic organic chemistry can address these fundamental challenges. Supported by molecular docking (Figures 2 and S1), a focused library of benzylidene imidazolinones was built to screen for redox-responsive pre-ligands of the RNA aptamer Baby Spinach (Figure 3). Among those, PyDSBz-DBrHBI and MBBA-MFHBI showed favorable solubility, stability, and redox selectivity profiles. In addition, both pre-ligands displayed markedly inhibited fluorogenic binding to Baby Spinach (Figure 4), providing minimal background signal generation. Notably, the pre-ligand-Baby Spinach system exhibited significant levels of fluorescence enhancement in the presence of Baby Spinach and their respective inorganic target, H_2_S or H_2_O_2_ (Figure 5). The target sensitivities and rates of fluorescence generation are comparable to previously reported fluorescent probes, and improving on these parameters warrants future pre-ligand designs with potential modifications on the benzylidene-caging motifs or imidazolinone heterocycle. We demonstrated the implementation of aptamer-based redox sensing in *E. coli* through plasmid transfection followed by pre-ligand incubation, which conveniently transformed them into cell-based sensors (Figure 6). This FLAP-mediated activity-based sensing approach is not limited to H_2_S or H_2_O_2_ and can, in principle, be tailored to detect other reactive molecular targets on demand. Collectively, our work demonstrates the potential of synthetic organic chemistry for advancing the applications of aptamers beyond their current capabilities in analyte sensing and traditional roles in synthetic biology.

## ACKNOWLEDGMENTS

We thank Edward W. Castner, Advaitha Madireddy, George Giambasu, Alec Meacham, Robert Puhak, Kern Hast and Jonathan Hong for useful discussions. We acknowledge Magali Rhia L. Stone for guidance in microscopy sample preparation. We thank Xinyi (Cindy) Miao for assistance in fluorescence quantification of *E*. *coli*. We acknowledge the Office of Advanced Research Computing (OARC) at Rutgers for providing access to the Amarel cluster. This work was supported by the US National Institutes of Health / National Institute of Biomedical Imaging and Bioengineering, Trailblazer Award (EB029548), the American Cancer Society, Institutional Research Grant Early Investigator Award, and the Rutgers Cancer Institute of New Jersey NCI Cancer Center Support Grant (P30CA072720), and Busch Biomedical Grant (to E.C.I.); and Steven A. Cox Scholarship for Cancer Research (to T.A.).

## AUTHOR CONTRIBUTIONS

E.C.I. conceived and supervised the study. T.A. conducted molecular docking, plasmid design, molecular biology experiments, and microscopy. L.W., B.G., H.E., and H.G. conducted the pre-ligand syntheses, spectroscopic and spectrometric characterizations. All of the authors contributed to the interpretation of data. T.A., L.W., and E.C.I. wrote the manuscript with input from other authors.

## REFERENCES

(1) Roth, A.; Breaker, R. R. The Structural and Functional Diversity of Metabolite-Binding Riboswitches. Annu. Rev. Biochem. 2009, 78 (1), 305–334. 10.1146/annurev.biochem.78.070507.135656.

(2) Ellington, A. D.; Szostak, J. W. In Vitro Selection of RNA Molecules That Bind Specific Ligands. Nature 1990, 346 (6287), 818–822. 10.1038/346818a0.

(3) Tuerk, C.; Gold, L. Systematic Evolution of Ligands by Exponential Enrichment: RNA Ligands to Bacteriophage T4 DNA Polymerase. Science 1990, 249 (4968), 505–510. 10.1126/science.2200121.

(4) Neubacher, S.; Hennig, S. RNA Structure and Cellular Applications of Fluorescent Light-Up Aptamers. Angew. Chem. Int. Ed. 2019, 58 (5), 1266–1279. 10.1002/anie.201806482.

(5) McConnell, E. M.; Nguyen, J.; Li, Y. Aptamer-Based Biosensors for Environmental Monitoring. Front. Chem. 2020, 8, 434. 10.3389/fchem.2020.00434.

(6) Gooch, J.; Daniel, B.; Parkin, M.; Frascione, N. Developing Aptasensors for Forensic Analysis. TrAC Trends Anal. Chem. 2017, 94, 150–160. 10.1016/j.trac.2017.07.019.

(7) Wang, X. C.; Wilson, S. C.; Hammond, M. C. Next-Generation RNA-Based Fluorescent Biosensors Enable Anaerobic Detection of Cyclic Di-GMP. Nucleic Acids Res. 2016, 44 (17), e139–e139. 10.1093/nar/gkw580.

(8) Kato, T.; Shimada, I.; Kimura, R.; Hyuga, M. Light-up Fluorophore–DNA Aptamer Pair for Label-Free Turn-on Aptamer Sensors. Chem. Commun. 2016, 52 (21), 4041–4044. 10.1039/c5cc08816j.

(9) Khan, P.; Aufdembrink, L. M.; Engelhart, A. E. Isothermal SARS-CoV-2 Diagnostics: Tools for Enabling Distributed Pandemic Testing as a Means of Supporting Safe Reopenings. ACS Synth. Biol. 2020, 9 (11), 2861–2880. 10.1021/acssynbio.0c00359.

(10) Kolpashchikov, D. M.; Spelkov, A. A. Binary (Split) Light-up Aptameric Sensors. Angew. Chem. Int. Ed. 2021, 60 (10), 4988–4999. 10.1002/anie.201914919.

(11) Wang, L.; Hast, K.; Aggarwal, T.; Baci, M.; Hong, J.; Izgu, E. C. MicroRNA Detection in Biologically Relevant Media Using a Split Aptamer Platform. Bioorg. Med. Chem. 2022, 69, 116909. 10.1016/j.bmc.2022.116909.

(12) Wu, R.; Mudiyanselage, A. P. K. K. K.; Shafiei, F.; Zhao, B.; Bagheri, Y.; Yu, Q.; McAuliffe, K.; Ren, K.; You, M. Genetically Encoded Ratiometric RNA-Based Sensors for Quantitative Imaging of Small Molecules in Living Cells. Angew. Chem. Int. Ed. 2019, 58 (50), 18271–18275. 10.1002/anie.201911799.

(13) Ren, K.; Wu, R.; Mudiyanselage, A. P. K. K. K.; Yu, Q.; Zhao, B.; Xie, Y.; Bagheri, Y.; Tian, Q.; You, M. In Situ Genetically Cascaded Amplification for Imaging RNA Subcellular Locations. J. Am. Chem. Soc. 2020, 142 (6), 2968–2974. 10.1021/jacs.9b11748.

(14) Sunbul, M.; Lackner, J.; Martin, A.; Englert, D.; Hacene, B.; Grün, F.; Nienhaus, K.; Nienhaus, G. U.; Jäschke, A. Super-Resolution RNA Imaging Using a Rhodamine-Binding Aptamer with Fast Exchange Kinetics. Nat. Biotechnol. 2021, 39 (6), 686–690. 10.1038/s41587-020-00794-3.

(15) Sun, Z.; Wu, R.; Zhao, B.; Zeinert, R.; Chien, P.; You, M. Live-Cell Imaging of Guanosine Tetra- and Pentaphosphate (p)PpGpp with RNA-based Fluorescent Sensors**. Angew. Chem. Int. Ed. 2021, 60 (45), 24070–24074. 10.1002/anie.202111170.

(16) Englert, D.; Burger, E.-M.; Grün, F.; Verma, M. S.; Lackner, J.; Lampe, M.; Bühler, B.; Schokolowski, J.; Nienhaus, G. U.; Jäschke, A.; Sunbul, M. Fast-Exchanging Spirocyclic Rhodamine Probes for Aptamer-Based Super-Resolution RNA Imaging. Nat. Commun. 2023, 14 (1), 3879. 10.1038/s41467-023-39611-1.

(17) Dunn, M. R.; McCloskey, C. M.; Buckley, P.; Rhea, K.; Chaput, J. C. Generating Biologically Stable TNA Aptamers That Function with High Affinity and Thermal Stability. J. Am. Chem. Soc. 2020, 142 (17), 7721–7724. 10.1021/jacs.0c00641.

(18) Climent-Catala, A.; Ouldridge, T. E.; Stan, G.-B. V.; Bae, W. Building an RNA-Based Toggle Switch Using Inhibitory RNA Aptamers. ACS Synth. Biol. 2022, 11 (2), 562–569. 10.1021/acssynbio.1c00580.

(19) Su, Y.; Hammond, M. C. RNA-Based Fluorescent Biosensors for Live Cell Imaging of Small Molecules and RNAs. Curr. Opin. Biotechnol. 2020, 63, 157–166. 10.1016/j.copbio.2020.01.001.

(20) You, M.; Litke, J. L.; Jaffrey, S. R. Imaging Metabolite Dynamics in Living Cells Using a Spinach-Based Riboswitch. Proc. Natl. Acad. Sci. 2015, 112 (21), E2756–E2765. 10.1073/pnas.1504354112.

(21) Spitale, R. C.; Incarnato, D. Probing the Dynamic RNA Structurome and Its Functions. Nat. Rev. Genet. 2023, 24 (3), 178–196. 10.1038/s41576-022-00546-w.

(22) Ganser, L. R.; Kelly, M. L.; Herschlag, D.; Al-Hashimi, H. M. The Roles of Structural Dynamics in the Cellular Functions of RNAs. Nat. Rev. Mol. Cell Biol. 2019, 20 (8), 474–489. 10.1038/s41580-019-0136-0.

(23) Hou, Q.; Jaffrey, S. R. Synthetic Biology Tools to Promote the Folding and Function of RNA Aptamers in Mammalian Cells. RNA Biol. 2023, 20 (1), 198–206. 10.1080/15476286.2023.2206248.

(24) Förster, U.; Weigand, J. E.; Trojanowski, P.; Suess, B.; Wachtveitl, J. Conformational Dynamics of the Tetracycline-Binding Aptamer. Nucleic Acids Res. 2012, 40 (4), 1807–1817. 10.1093/nar/gkr835.

(25) Paige, J. S.; Wu, K. Y.; Jaffrey, S. R. RNA Mimics of Green Fluorescent Protein. Science 2011, 333 (6042), 642–646. 10.1126/science.1207339.

(26) Babendure, J. R.; Adams, S. R.; Tsien, R. Y. Aptamers Switch on Fluorescence of Triphenylmethane Dyes. J. Am. Chem. Soc. 2003, 125 (48), 14716–14717. 10.1021/ja037994o.

(27) Warner, K. D.; Chen, M. C.; Song, W.; Strack, R. L.; Thorn, A.; Jaffrey, S. R.; Ferré-D’Amaré, A. R. Structural Basis for Activity of Highly Efficient RNA Mimics of Green Fluorescent Protein. Nat. Struct. Mol. Biol. 2014, 21 (8), 658–663. 10.1038/nsmb.2865.

(28) Huang, H.; Suslov, N. B.; Li, N.-S.; Shelke, S. A.; Evans, M. E.; Koldobskaya, Y.; Rice, P. A.; Piccirilli, J. A. A G-Quadruplex–Containing RNA Activates Fluorescence in a GFP-like Fluorophore. Nat. Chem. Biol. 2014, 10 (8), 686–691. 10.1038/nchembio.1561.

(29) Strack, R. L.; Disney, M. D.; Jaffrey, S. R. A Superfolding Spinach2 Reveals the Dynamic Nature of Trinucleotide Repeat RNA. Nat. methods 2013, 10 (12), 1219–1224. 10.1038/nmeth.2701.

(30) Olas, B. Hydrogen Sulfide as a “Double-Faced” Compound: One with Pro- and Antioxidant Effect. Adv. Clin. Chem. 2016, 78, 187–196. 10.1016/bs.acc.2016.07.005.

(31) Kimura, H. Hydrogen Sulfide Induces Cyclic AMP and Modulates the NMDA Receptor. Biochem. Biophys. Res. Commun. 2000, 267 (1), 129–133. 10.1006/bbrc.1999.1915.

(32) Han, Y.; Qin, J.; Chang, X.; Yang, Z.; Bu, D.; Du, J. Modulating Effect of Hydrogen Sulfide on Gamma-Aminobutyric Acid B Receptor in Recurrent Febrile Seizures in Rats. Neurosci. Res. 2005, 53 (2), 216–219. 10.1016/j.neures.2005.07.002.

(33) Sun, W.-H.; Liu, F.; Chen, Y.; Zhu, Y.-C. Hydrogen Sulfide Decreases the Levels of ROS by Inhibiting Mitochondrial Complex IV and Increasing SOD Activities in Cardiomyocytes under Ischemia/Reperfusion. Biochem. Biophys. Res. Commun. 2012, 421 (2), 164–169. 10.1016/j.bbrc.2012.03.121.

(34) Kimura, Y.; Goto, Y.-I.; Kimura, H. Hydrogen Sulfide Increases Glutathione Production and Suppresses Oxidative Stress in Mitochondria. Antioxid. Redox Signal. 2010, 12 (1), 1–13. 10.1089/ars.2008.2282.

(35) Lefer, D. J. A New Gaseous Signaling Molecule Emerges: Cardioprotective Role of Hydrogen Sulfide. Proc. Natl. Acad. Sci. 2007, 104 (46), 17907–17908. 10.1073/pnas.0709010104.

(36) Polhemus, D. J.; Lefer, D. J. Emergence of Hydrogen Sulfide as an Endogenous Gaseous Signaling Molecule in Cardiovascular Disease. Circ. Res. 2014, 114 (4), 730–737. 10.1161/circresaha.114.300505.

(37) Mathai, J. C.; Missner, A.; Kügler, P.; Saparov, S. M.; Zeidel, M. L.; Lee, J. K.; Pohl, P. No Facilitator Required for Membrane Transport of Hydrogen Sulfide. Proc. Natl. Acad. Sci. 2009, 106 (39), 16633–16638. 10.1073/pnas.0902952106.

(38) D’Autréaux, B.; Toledano, M. B. ROS as Signalling Molecules: Mechanisms That Generate Specificity in ROS Homeostasis. Nat. Rev. Mol. Cell Biol. 2007, 8 (10), 813–824. 10.1038/nrm2256.

(39) Sies, H.; Jones, D. P. Reactive Oxygen Species (ROS) as Pleiotropic Physiological Signalling Agents. Nat. Rev. Mol. Cell Biol. 2020, 21 (7), 363–383. 10.1038/s41580-020-0230-3.

(40) Sies, H. Role of Metabolic H2O2 Generation REDOX SIGNALING AND OXIDATIVE STRESS*. J. Biol. Chem. 2014, 289 (13), 8735–8741. 10.1074/jbc.r113.544635.

(41) Lennicke, C.; Rahn, J.; Lichtenfels, R.; Wessjohann, L. A.; Seliger, B. Hydrogen Peroxide – Production, Fate and Role in Redox Signaling of Tumor Cells. Cell Commun. Signal. : CCS 2015, 13 (1), 39. 10.1186/s12964-015-0118-6.

(42) Cordeiro, J. V.; Jacinto, A. The Role of Transcription-Independent Damage Signals in the Initiation of Epithelial Wound Healing. Nat. Rev. Mol. Cell Biol. 2013, 14 (4), 249–262. 10.1038/nrm3541.

(43) Cheung, E. C.; Vousden, K. H. The Role of ROS in Tumour Development and Progression. Nat. Rev. Cancer 2022, 22 (5), 280–297. 10.1038/s41568-021-00435-0.

(44) Bienert, G. P.; Møller, A. L. B.; Kristiansen, K. A.; Schulz, A.; Møller, I. M.; Schjoerring, J. K.; Jahn, T. P. Specific Aquaporins Facilitate the Diffusion of Hydrogen Peroxide across Membranes*. J. Biol. Chem. 2007, 282 (2), 1183–1192. 10.1074/jbc.m603761200.

(45) Bienert, G. P.; Desguin, B.; Chaumont, F.; Hols, P. Channel-Mediated Lactic Acid Transport: A Novel Function for Aquaglyceroporins in Bacteria. Biochem. J. 2013, 454 (3), 559–570. 10.1042/bj20130388.

(46) Trott, O.; Olson, A. J. AutoDock Vina: Improving the Speed and Accuracy of Docking with a New Scoring Function, Efficient Optimization, and Multithreading. J. Comput. Chem. 2010, 31 (2), 455–461. 10.1002/jcc.21334.

(47) Eberhardt, J.; Santos-Martins, D.; Tillack, A. F.; Forli, S. AutoDock Vina 1.2.0: New Docking Methods, Expanded Force Field, and Python Bindings. J. Chem. Inf. Model. 2021, 61 (8), 3891–3898. 10.1021/acs.jcim.1c00203.

(48) Su, M.; Yang, Q.; Du, Y.; Feng, G.; Liu, Z.; Li, Y.; Wang, R. Comparative Assessment of Scoring Functions: The CASF-2016 Update. J. Chem. Inf. Model. 2019, 59 (2), 895–913. 10.1021/acs.jcim.8b00545.

(49) Wang, Z.; Li, Y.; Zhang, Q.; Jing, C.; Jiang, Y.; Yang, T.; Han, T.; Xiong, F. A Highly Selective and Easily Acquisitive Near-Infrared Fluorescent Probe for Detection and Imaging of Hydrogen Sulfide in Cells. Spectrochim. Acta Part A: Mol. Biomol. Spectrosc. 2023, 293, 122428. 10.1016/j.saa.2023.122428.

(50) Lin, V. S.; Chen, W.; Xian, M.; Chang, C. J. Chemical Probes for Molecular Imaging and Detection of Hydrogen Sulfide and Reactive Sulfur Species in Biological Systems. Chem. Soc. Rev. 2014, 44 (14), 4596–4618. 10.1039/c4cs00298a.

(51) Peng, B.; Chen, W.; Liu, C.; Rosser, E. W.; Pacheco, A.; Zhao, Y.; Aguilar, H. C.; Xian, M. Fluorescent Probes Based on Nucleophilic Substitution–Cyclization for Hydrogen Sulfide Detection and Bioimaging. Chem. A Eur. J. 2014, 20 (4), 1010–1016. 10.1002/chem.201303757.

(52) Lippert, A. R.; Bittner, G. C. V. de; Chang, C. J. Boronate Oxidation as a Bioorthogonal Reaction Approach for Studying the Chemistry of Hydrogen Peroxide in Living Systems. Acc. Chem. Res. 2011, 44 (9), 793–804. 10.1021/ar200126t.

(53) Cuevasanta, E.; Lange, M.; Bonanata, J.; Coitiño, E. L.; Ferrer-Sueta, G.; Filipovic, M. R.; Alvarez, B. Reaction of Hydrogen Sulfide with Disulfide and Sulfenic Acid to Form the Strongly Nucleophilic Persulfide. J. Biol. Chem. 2015, 290 (45), 26866–26880. 10.1074/jbc.m115.672816.

(54) Yadav, A. K.; Zhao, Z.; Weng, Y.; Gardner, S. H.; Brady, C. J.; Peguero, O. D. P.; Chan, J. Hydrolysis-Resistant Ester-Based Linkers for Development of Activity-Based NIR Bioluminescence Probes. J Am Chem Soc 2023. 10.1021/jacs.2c12984.

(55) Papapetropoulos, A.; Pyriochou, A.; Altaany, Z.; Yang, G.; Marazioti, A.; Zhou, Z.; Jeschke, M. G.; Branski, L. K.; Herndon, D. N.; Wang, R.; Szabó, C. Hydrogen Sulfide Is an Endogenous Stimulator of Angiogenesis. Proc National Acad Sci 2009, 106 (51), 21972–21977. 10.1073/pnas.0908047106.

(56) Wang, R. Physiological Implications of Hydrogen Sulfide: A Whiff Exploration That Blossomed. Physiol Rev 2012, 92 (2), 791–896. 10.1152/physrev.00017.2011.

(57) Pose, M.; Dillon, K. M.; Denicola, A.; Alvarez, B.; Matson, J. B.; Möller, M. N.; Cuevasanta, E. Fluorescent Detection of Hydrogen Sulfide (H2S) through the Formation of Pyrene Excimers Enhances H2S Quantification in Biochemical Systems. J. Biol. Chem. 2022, 298 (10), 102402. 10.1016/j.jbc.2022.102402.

(58) Peng, S.; Zhong, T.; Guo, T.; Shu, D.; Meng, D.; Liu, H.; Guo, D. A Novel Fluorescent Probe for Selective Detection of Hydrogen Sulfide in Living Cells. N. J. Chem. 2018, 42 (7), 5185–5192. 10.1039/c7nj04577h.

(59) Chen, H.; Wu, X.; Yang, S.; Tian, H.; Liu, Y.; Sun, B. A Visible Colorimetric Fluorescent Probe for Hydrogen Sulfide Detection in Wine. J. Anal. Methods Chem. 2019, 2019, 2173671. 10.1155/2019/2173671.

(60) Bolland, H. R.; Hammond, E. M.; Sedgwick, A. C. A Fluorescent Probe Strategy for the Detection and Discrimination of Hydrogen Peroxide and Peroxynitrite in Cells. Chem. Commun. 2022, 58 (76), 10699–10702. 10.1039/d2cc03406a.

(61) Zuo, Y.; Jiao, Y.; Ma, C.; Duan, C. A Novel Fluorescent Probe for Hydrogen Peroxide and Its Application in Bio-Imaging. Molecules 2021, 26 (11), 3352. 10.3390/molecules26113352.

(62) Khalil, A. S.; Collins, J. J. Synthetic Biology: Applications Come of Age. Nat. Rev. Genet. 2010, 11 (5), 367–379. 10.1038/nrg2775.

(63) Shechner, D. M.; Hacisuleyman, E.; Younger, S. T.; Rinn, J. L. Multiplexable, Locus-Specific Targeting of Long RNAs with CRISPR-Display. Nat. Methods 2015, 12 (7), 664–670. 10.1038/nmeth.3433.

