## Supplementary Information for "A Small-Molecule Approach to Bypass In Vitro Selection of New Aptamers: Designer Pre-Ligands Turn Baby Spinach into Sensors for Reactive Inorganic Targets"

| Contents | Page |
|---|---|
| **Chemicals** | **S3** |
| **General Synthetic Methods and Instrumental Analyses** | **S4** |
| **Preparation and Characterization of Small Molecules** | **S5–S39** |
| AEC-MFHBI | S5 |
| AEC-DFHBI | S6 |
| DNP-MFHBI | S7 |
| PyDSBz-MFHBI | S8 |
| PyDSBz-DBrHBI | S9 |
| MBBAPE-MFHBI | S10 |
| MBBAPE-DFHBI | S11 |
| MBBA-MFHBI | S12 |
| MC-MFHBI | S13 |
| $^{1}$H, $^{13}$C, $^{19}$F NMR Spectra | S14–S39 |
| **Docking Simulation Method** | **S40** |
| Figure S1 | S41 |
| **RSS selectivity of AEC-MFHBI and MC-MFHBI** | **S42** |
| Figure S2 | |
| **Determination of pKa of DBrHBI and MFHBI** | **S43** |
| Figure S3 and S4 | |
| **Fluorescence Measurement Protocols** | **S44** |
| **Protocols for Cell Studies** | **S45** |
| **Fluorescence imaging quantification using ImageJ** | **S46** |
| **Sequence and Proposed 2-Dimensional Structure of 3xBaby Spinach** | **S47** |
| Figure S5 | |
| **Plasmid Map of pET-21-3xBaby Spinach** | **S48** |
| Figure S6 | |
| **References** | **S49** |

### Chemicals

**Reagents:** Hydrogen peroxide ($H_2O_2$), sodium hydroxide (NaOH), triphosgene, sodium bicarbonate ($NaHCO_3$), sodium sulfate ($Na_2SO_4$), 1-octanol, sucrose, sodium nitrite ($NaNO_2$), potassium superoxide ($KO_2$), sodium ascorbate, methyl amine, benzyl bromide, Dihydroxyethyl disulfide, diisopropylethylamine, *N*-acetylglycine, acetic anhydride, triethylamine, 1-azidoethanol, methylchloroformate, pyridine, dimethylformamide (DMF), 4-Bromomethylphenylboronic acid pinacol ester, 4-Bromomethylphenylboronic acid, Ampicillin (Amp), luria broth (LB), isopropyl-β-D-thiogalactopyranoside (IPTG), sodium chloride (NaCl), hydrochloric acid (HCl), glycerol, potassium chloride (KCl), potassium carbonate ($K_2CO_3$), ethylenediaminetetraacetic acid (EDTA), magnesium chloride ($MgCl_2$), calcium chloride ($CaCl_2$), Sodium sulfide nonahydrate ($Na_2S{\cdot}9H_2O$), Ampicillin (Amp) were purchased from Millipore-Sigma. Deuterated solvents, containing 0.05% (v/v) TMS, were purchased from either Cambridge Isotope Laboratories or Millipore Sigma.

**Buffers, Solvents, Media:** Phosphate-buffered saline (PBS), Luria broth (LB), HEPES, tris(hydroxymethyl)aminomethane (Tris), dichloromethane ($CH_2Cl_2$), ethyl acetate (EtOAc), benzene, toluene, tetrahydrofuran (THF), and chloroform ($CHCl_3$) were purchased from Millipore Sigma. Acetonitrile ($CH_3CN$) and methanol (MeOH, HPLC grade) were purchased from Fisher Scientific. Water was deionized and filtered to a resistivity of 18.2 ΩM·cm with a Milli-Q® Plus water purification system (Millipore, Massachusetts). Buffers were prepared freshly in Milli-Q® water and their pH were adjusted using HCl or NaOH using a Thermo Scientific Orion Star pH meter. Hanks' Balanced Salt Solution (HBSS), Petri dish, culture tubes were purchased from VWR International.

**Biologics:** All the recombinant DNA and molecular biology work was carried out in a biosafety cabinet; these materials and protocols were approved by Rutgers Institutional biosafety committee. BL21-star-DE3 competent *Escherichia coli* cells were purchased from New England Biolabs. Baby Spinach (5'-GGUGAAGGACGGGUCCAGUAGUUCGCUACUGUUGAGUAGAGU GUGAGCUCC-3') was obtained either commercially (Sigma or IDT) or from cell-free transcription using a T7 RNA transcription kit (New England Biolabs). The DNA plasmid pET-21-3xBabySpinach encoding for tRNA-3xBabySpinach construct was custom designed and was purchased from Twist Bioscience.

### General Synthetic Methods and Instrumental Analyses

All reactions were performed under a dry nitrogen atmosphere unless otherwise stated. All glassware was oven-dried before use. Purification of the synthesized compounds was performed using a Büchi Reveleris® flash chromatography system equipped with a FlashPure EcoFlex Diol (50 µm spherical) column or a silica (50 µm irregular) column. Nuclear magnetic resonance (NMR) spectroscopic analyses were carried out using a Bruker Avance Neo 500 MHz spectrometer. NMR data is provided for new compounds. $^{1}$H NMR spectra were acquired at 500 MHz, $^{13}$C NMR spectra were acquired at 126 MHz, and $^{19}$F NMR spectra were acquired at 470 MHz. Chemical shifts ($\delta$) for $^{1}$H NMR spectra were referenced to $(C\boldsymbol{H}_3)_4Si$ at $\delta$ = 0.00 ppm, to $C\boldsymbol{H}Cl_3$ at $\delta$ = 7.26 ppm, or to $C\boldsymbol{H}D_2S(O)CD_3$ at $\delta$ = 2.50 ppm. $^{13}$C NMR spectra were referenced to $\boldsymbol{C}DCl_3$ at $\delta$ = 77.23 ppm or to $(\boldsymbol{C}D_3)_2SO$ at $\delta$ = 39.52 ppm.

The following abbreviations are used to describe $^{1}$H NMR resonances: s (singlet), d (doublet), t (triplet), m (multiplet), dd (doublet of doublets), br (broad), and app (apparent). Coupling constants (*J*) are reported in Hz. Liquid chromatography followed by high-resolution mass spectrometry (LC-HRMS) analysis from electrospray ionization (ESI) was carried out on a Waters Acquity-Xevo G2-XS QTof instrument. Low-resolution mass spectrometry (LRMS) analysis was performed using a Finnigan LCQ™ DUO mass spectrometer. Both fluorescence and UV-Vis absorbance measurements were conducted using a Spectra Max ID3 instrument (Molecular Devices). Confocal fluorescence microscopy was performed on a Leica TCS SP8 tauSTED microscope with a 63”, 1.40 NA oil Immersion objective and Images were processed using LasX Lightning deconvolution. All statistical analyses were performed using OriginLab software or GraphPad Prism software version 9.

### Preparation and Characterization of Small Molecules

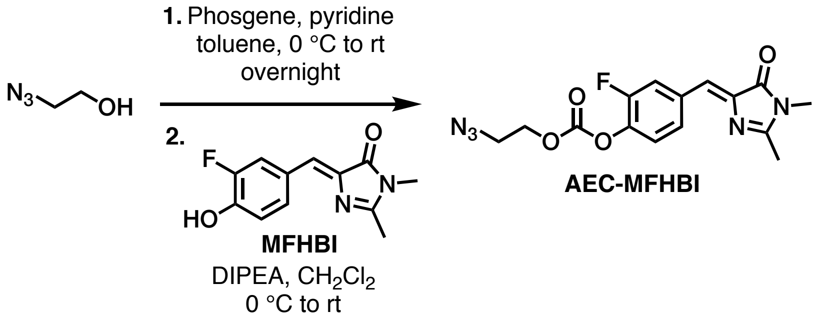


**Synthesis of AEC-MFHBI:** To a dry 10-mL round-bottom flask containing 2-azidoethanol (43.5 mg, 0.50 mmol, 1.0 equiv) dissolved in 3.5 mL of toluene was added phosgene (1.43 mL, 2.00 mmol, 4.0 equiv, wt. 15% in toluene) and mixture was allowed to stir for 2 hours at 0 °C. The reaction solution was purged with nitrogen and concentrated under reduced pressure. The chloroformate was dissolved in 2 mL of $CH_2Cl_2$ and cooled to 0°C under a nitrogen atmosphere. A mixture of MFHBI (117 mg, 0.50 mmol, 1.0 equiv) and DIPEA (0.26 mL, 1.5 mmol, 3.0 equiv) in $CH_2Cl_2$ (3 mL) were added dropwise to the reaction vessel. The reaction was allowed to stir at room temperature for 20 hours. The reaction was concentrated to a solid under reduced pressure and then, purified by column chromatography using a silica column (Hex/EtOAc from 0 to 60%), providing AEC-MFHBI (128 mg, 74%) as a light-yellow solid.

**$^1$H NMR** (500 MHz, $CDCl_3$): δ 8.24 (dd, *J* = 11.6, 1.9 Hz, 1H), 7.73 (dt, *J* = 8.4, 1.5 Hz, 1H), 7.26 (d, *J* = 8.2 Hz, 1H), 6.99 (s, 1H), 4.43 (t, *J* = 5.2 Hz, 2H), 3.61 (t, *J* = 5.2 Hz, 2H), 3.19 (s, 3H), 2.38 (s, 3H).

**$^{13}$C NMR** (126 MHz, $CDCl_3$): δ 170.5, 163.9, 153.8 (d, *J* = 249.8 Hz), 152.2, 139.9, 139.3 (d, *J* = 12.9 Hz), 134.1 (d, *J* = 7.7 Hz), 128.7 (d, *J* = 3.4 Hz), 124.3 (d, *J* = 2.7 Hz), 123.2, 119.8 (d, *J* = 20.0 Hz), 67.5, 49.6, 26.7, 15.8.

**$^{19}$F NMR** (471 MHz, $CDCl_3$): δ -128.1 (m).

**HRMS** (ESI) m/z: Calculated for $C_{15}H_{15}FN_5O_4^+$, $[M + H]^+$ requires 348.1103; found 348.1109.

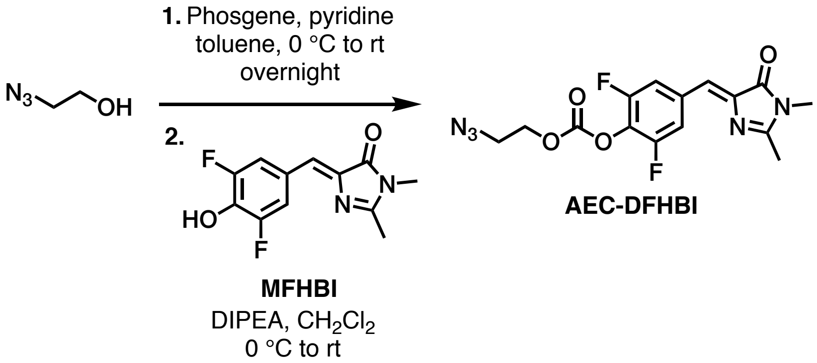


**Synthesis of *AEC-DFHBI***: To a dry 10-mL round-bottom flask containing 2-azidoethanol (22 mg, 0.25 mmol, 1.0 equiv) dissolved in 2.0 mL of toluene was added phosgene (0.72 mL, 1.00 mmol, 4.0 equiv, wt. 15% in toluene) and mixture was allowed to stir for 2 hours at 0°C. The reaction solution was purged with nitrogen and concentrated under reduced pressure. The chloroformate was dissolved in 1 mL of $CH_2Cl_2$ and cooled to 0°C under a nitrogen atmosphere. A mixture of DFHBI (63 mg, 0.25 mmol, 1.0 equiv) and DIPEA (0.13 mL, 0.75 mmol, 3.0 equiv) in $CH_2Cl_2$ (2 mL) were added dropwise to the reaction vessel. The reaction was allowed to stir at RT for 20 hours. The reaction was concentrated to a solid under reduced pressure and then, purified by column chromatography using a silica column (Hex/EtOAc from 0 to 60%), providing AEC-DFHBI as a light-yellow solid (59 mg, 65%).

**$^1$H NMR** (500 MHz, $CDCl_3$): δ 7.83 (d, *J* = 8.8 Hz, 2H), 6.88 (s, 1H), 4.44 (t, *J* = 5.2 Hz, 2H), 3.62 (t, *J* = 5.2 Hz, 2H), 3.18 (s, 3H), 2.38 (s, 3H).

**$^{13}$C NMR** (126 MHz, $CDCl_3$): δ 170.4, 164.7, 155.9, 153.9, 151.7, 140.8, 133.7, 128.5, 128.2, 123.1, 115.5, 66.1, 49.6, 26.8, 15.9.

**$^{19}$F NMR** (471 MHz, $CDCl_3$): δ -126.1 (s).

**HRMS** (ESI) m/z: Calculated for $C_{15}H_{14}F_2N_5O_4^+$, $[M + H]^+$ requires 366.1008; found 366.1012.

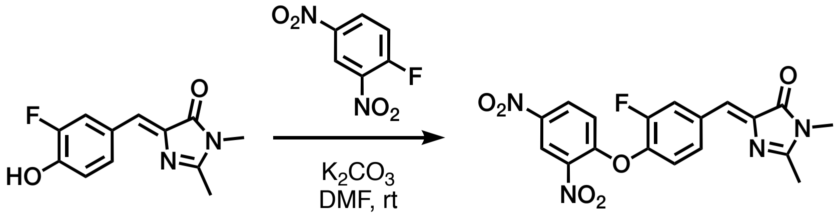


**MFHBI** **DNP-MFHBI**

**Synthesis of DNP-MFHBI:** To a dry 10-mL vial containing MFHBI (100.0 mg, 0.427 mmol, 1.0 equiv) dissolved in 4.0 mL of $CH_2Cl_2$ was added $K_2CO_3$ (59.0 mg, 0.427 mmol, 1.0 equiv) and 1-fluoro-2,4-dinitrobenzene (119.2 mg, 0.641 mmol, 1.5 equiv). The vial was capped and the mixture was stirred at room temperature overnight. The reaction mixture was then filtered and the solid residue was washed with $CH_2Cl_2$. Residual solvent was removed from the residue under reduced pressure, providing DNP-MFHBI as a bright-yellow solid (165.2 mg, 97%).

**$^1$H NMR** (500 MHz, $CDCl_3$): δ 8.89 (d, J = 2.7 Hz, 1H), 8.36-8.32 (m, 2H), 7.82f (d, J = 8.4 Hz, 1H), 7.27 (t, J = 8.2 Hz, 1H), 7.03 (dd, J = 9.2, 1.1 Hz, 1H), 7.01(s, 1H), 3.21 (s, 3H), 2.41 (s, 3H)

**$^{13}$C NMR** (126 MHz, $CDCl_3$): δ 170.48, 164.55, 155.22, 153.68 (d, *J* = 251.3 Hz), 142.17, 141.28, 140.30, 139.13, 134.72 (d, *J* = 7.5 Hz), 129.59 (d, *J* = 3.4 Hz), 129.07, 123.72 (d, *J* = 2.6 Hz), 123.01, 122.43, 120.66 (d, *J* = 19.3 Hz), 117.76, 26.85, 15.93.

**$^{19}$F NMR** (471 MHz, $CDCl_3$): δ -127.3 (m).

**HRMS** (ESI) m/z: Calculated for $C_{18}H_{14}FN_4O_6^+$, $[M + H]^+$ requires 401.0892; found 401.0913.

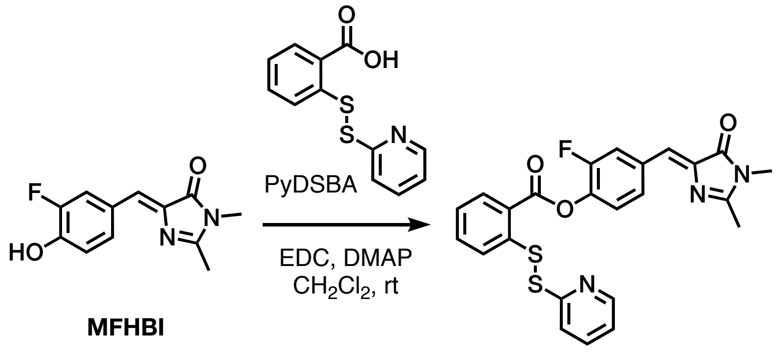


**Synthesis of PyDSBz-MFHBI:** To a dried 25-mL round bottom containing PyDSBA (100.0 mg, 0.38 mmol, 1.0 equiv) dissolved in 10 mL anhydrous $CH_2Cl_2$ was added EDC (72.8 mg, 0.38 mmol, 1.0 equiv), and mixture was stirred at room temperature for 15 min. Then to the reaction solution, MFHBI (88.9 mg, 0.38 mmol, 1.0 equiv) and DMAP (4.6 mg, 0.04 mmol, 0.1 equiv) were added. The reaction mixture was stirred at room temperature for 24 hours. $CH_2Cl_2$ was removed by rotavapor and the solid was redissolved in 50% acetonitrile in water (v/v), purified by a C18 column (water/acetonitrile with 0.1% TFA, from 5% to 95%), affording PyDSBz-MFHBI (82.0 mg, 45%) as a light yellow solid.

**$^1$H NMR** (500 MHz, $CDCl_3$): δ 8.33 (d, *J* = 8.5 Hz, 1H), 8.28 (d, *J* = 11.5 Hz, 1H), 7.86-7.77 (m, 3H), 7.50 (m, 1H), 7.35 (m, 1H), 7.29 (t, *J* = 7.4 Hz, 1H), 7.06 (s, 1H), 3.21 (s, 3H), 2.41 (s, 1H)

**$^{13}$C NMR** (126 MHz, $CDCl_3$): δ 170.68, 169.91, 163.84, 163.73, 154.17 (d, *J* = 249.7 Hz), 142.02, 141.19, 139.62, 139.44, 139.34, 134.24, 134.05, 132.59 (d, *J* = 8.3 Hz), 128.83(d, *J* = 2.9 Hz), 126.36, 125.90 (d, *J* = 7.8 Hz), 126.07 (d, *J* = 36.8 Hz), 124.99, 124.11, 119.89 (d, *J* = 19.9 Hz), 26.82, 15.86.

**$^{19}$F NMR** (471 MHz, $CDCl_3$): δ -127.0 (m).

**HRMS** (ESI) m/z: Calculated for $C_{15}H_{14}F_2N_5O_4^+$, $[M + H]^+$ requires 480.0847; found 480.0873.

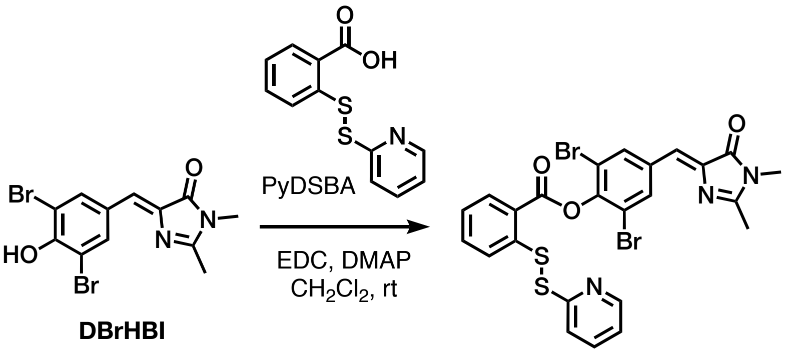


**PyDSBz-DBrHBI**

**Synthesis of PyDSBz-DBrHBI:** Text to be written by Liming. To a dried 25-mL round bottom containing PyDSBA (168.8 mg, 0.64 mmol, 1.2 equiv) dissolved in 10 mL anhydrous $CH_2Cl_2$ was added EDC (122.9 mg, 0.64 mmol, 1.2 equiv), and mixture was stirred at room temperature for 15 minutes. Then to the reaction solution, DBrHBI (200.0 mg, 0.53 mmol, 1.0 equiv) and DMAP (6.4 mg, 0.05 mmol, 0.1 equiv) were added. The reaction mixture was stirred at room temperature for 24 hours. Half of the $CH_2Cl_2$ was removed by rotavapor and remaining mixture was injected to a neutral aluminum oxide column (Hex/EtoAc, from 10% to 55%), affording PyDSBz-DBrHBI (20.2 mg, 5%) as a white solid.

**$^1$H NMR** (500 MHz, $CDCl_3$): δ 8.48 (app d, *J* = 4.8 Hz, 1H), 8.43 (s, 2H), 8.41 (dd, *J* = 8.0, 1.2 Hz, 1H), 7.99 (d, *J* = 8.0 Hz, 1H), 7.63–7.55 (m, 3H), 7.37 (t, *J* = 7.4 Hz, 1H), 7.10 (m, 1H), 6.91 (s, 1H), 3.19 (s, 3H), 2.41 (s, 3H).

**$^{13}$C NMR** (126 MHz, $CDCl_3$): δ 170.5, 164.8, 162.7, 159.2, 149.8, 146.9, 142.4, 140.7, 137.6, 135.6, 135.3, 134.5, 132.6, 126.6, 126.2, 125.5, 122.7, 121.3, 119.9, 118.2, 26.9, 16.0.

**HRMS** (ESI) m/z: Calculated for $C_{15}H_{14}F_2N_5O_4^+$, $[M + H]^+$ requires 619.9131; found 619.9120.

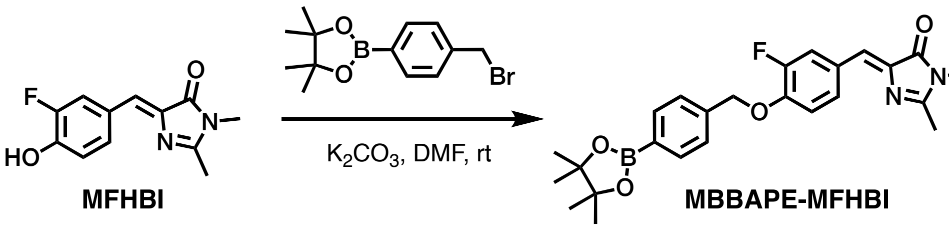


**Synthesis of MBBAPE-MFHBI:** To a dry 25-mL round-bottom flask MFHBI (200 mg, 0.85 mmol, 1.0 equiv), 4-Bromomethylphenylboronic acid pinacol ester (330 mg, 1.10 mmol, 1.3 equiv), $K_2CO_3$ (410 mg, 2.98 mmol, 3.5 equiv) were added in 17 mL of DMF under a nitrogen environment. The reaction was allowed to stir at room temperature and monitored by TLC until completion (~23 hours). The mixture was diluted with ethyl acetate and the organic layer was washed with water 3 times. The collected aqueous layers were washed with ethyl acetate 3 times. Lastly, the collected organic layers were washed with saturated aqueous NaCl. The organic layer was collected and dried with sodium sulfate. The organic solvent was removed under reduced pressure and the crude solid was washed with cold methanol, providing MBBAPE-MFHBI (240 mg, 63%) as a light-yellow solid.

**$^1$H NMR** (500 MHz, $CDCl_3$): δ 8.47-8.44 (d, *J* = 12.3 Hz, 1H), 7.83-7.81 (d, *J* = 9.0 Hz, 2H), 7.78-7.76 (d, *J* = 8.5 Hz, 2H), 7.44-7.42 (d, *J* = 9.0 Hz, 2H), 7.11 (s), 6.97-6.94 (t, *J* = 9.0 Hz, 1H), 3.17 (s, 3H), 2.30 (s, 3H), 1.33 (s, 12H).

**$^{13}$C NMR** (126 MHz, $CDCl_3$): δ 167.6, 159.0, 153.2, 151.2, 149.0, 139.2, 138.8, 136.2, 135.3, 129.6, 127.5, 126.6, 119.3, 114.4, 84.1, 71.1, 26.8, 25.1, 15.6.

**$^{19}$F NMR** (471 MHz, $CDCl_3$): δ -132.9 (m).

**HRMS** (ESI) m/z: Calculated for $C_{25}H_{29}BFN_2O_4^+$, $[M + H]^+$ requires 451.2199; found 451.2183.

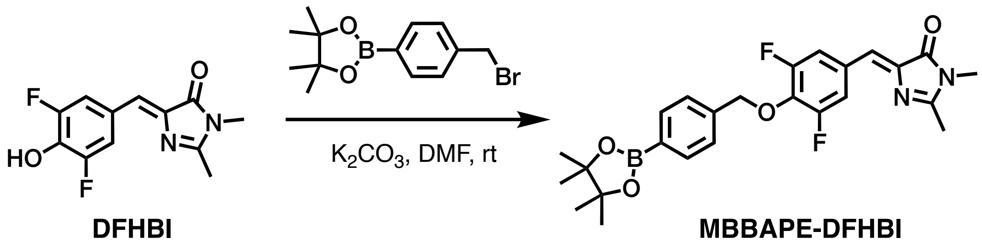


**Synthesis of MBBAPE-DFHBI:** To a dry 10-mL round-bottom flask DFHBI (200 mg, 1.00 mmol, 1.0 equiv), 4-bromomethylphenylboronic acid pinacol ester (300 mg, 0.96 mmol, 1.2 equiv), triethylamine (0.27 mL, 2.00 mmol, 2.5 equiv) were added in 4 mL of $CH_2Cl_2$ under a nitrogen environment. The reaction was allowed to stir at room temperature and monitored by TLC until completion (18 hours). The reaction was diluted in $CH_2Cl_2$, filtered, and concentrated to a solid under reduced pressure and purified by column chromatography using a silica column ($CH_2Cl_2$/MeOH from 0 to 4%), affording MBBAPE-DFHBI (81 mg, 21%) as a yellow solid.

**$^1$H NMR** (500 MHz, $CDCl_3$): $\delta$ 7.80-7.79 (d, $J$ = 7.99 Hz, 2H), 7.72-7.70 (d, $J$ = 9.11 Hz, 2H), 7.44-7.43 (d, $J$ = 7.86 Hz, 2H), 6.87 (s, 1H), 5.26 (s, 2H), 3.18 (s, 3H), 2.38 (s, 3H), 1.26 (s, 12H).

**$^{13}$C NMR** (126 MHz, $(CD_3)_2SO$): $\delta$ 169.5, 165.6, 155.8, 153.8, 139.7, 139.4, 135.3, 134.3, 129.8, 127.5, 121.6, 115.5, 83.7, 75.3, 26.2, 24.6, 15.4.

**$^{19}$F NMR** (471 MHz, $(CD_3)_2SO$): $\delta$ -127.4 (d, $J$ = 9.1 Hz).

**LRMS** (ESI) m/z: Calculated for $C_{25}H_{28}BF_2N_2O_4^+$, $[M + H]^+$ requires 469.21; found 469.22.

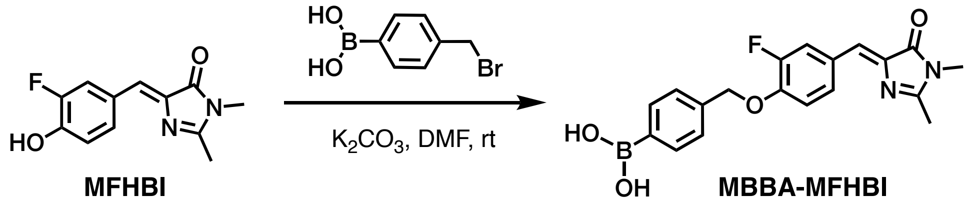


**Synthesis of MBBA-MFHBI:** To a dry 25-mL round-bottom flask MFHBI (50 mg, 0.21 mmol, 1.0 equiv), 4-bromomethylphenylboronic acid (60 mg, 0.28 mmol, 1.3 equiv), $K_2CO_3$ (100 mg, 0.74 mmol, 3.5 equiv) were added in 4.2 mL of DMF under a nitrogen environment. The reaction was allowed to stir at room temperature and monitored by TLC until completion (48 h). The mixture was diluted with ethyl acetate and the organic layer was washed with water 3 times. The collected aqueous layers were washed with ethyl acetate 3 times. Lastly, the collected organic layers were washed with brine. The organic layer was collected and dried with sodium sulfate. The organic solvent was removed under reduced pressure and the crude solid was washed with cold methanol, providing MBBA-MFHBI (32 mg, 40%) as a light-yellow solid.

**$^1$H NMR** (500 MHz, $(CD_3)_2SO$): δ 8.30-8.27 (d, *J* = 12.5 Hz, 1H), 8.07 (s, 2H), 7.86-7.84 (d, *J* = 9.1 Hz, 1H), 7.82-7.81 (d, *J* = 9.1 Hz, 2H), 7.43-7.42 (d, *J* = 9.4 Hz, 2H), 7.32-7.28 (t, *J* = 9.1 Hz, 1H), 6.93 (s, 1H), 5.25 (s, 2H), 3.08 (s, 3H), 2.35 (s, 3H).

**$^{13}$C NMR** (126 MHz, $(CD_3)_2SO$): δ 170.2, 164.6, 152.8, 150.9, 148.2, 138.6, 134.8, 129.9, 128.0, 127.3, 123.9, 118.8, 115.5, 70.7, 26.7, 15.9.

**$^{19}$F NMR** (471 MHz, $(CD_3)_2SO$): δ -134.1 (m).

**LRMS** (ESI) m/z: Calculated for $C_{19}H_{19}BFN_2O_4^+$, $[M + H]^+$ requires 369.14; found 369.15.

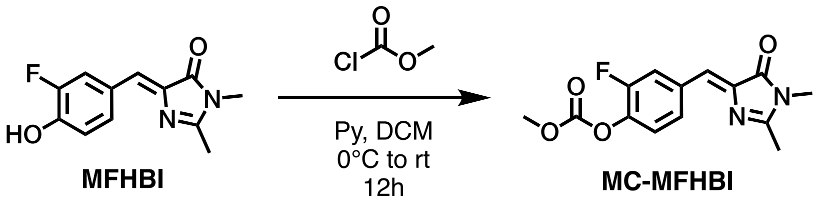


**Synthesis of MC-MFHBI:** To a dry 25-mL round-bottom flask methyl chloroformate (0.08 mL, 1.06 mmol, 2 equiv) and $CH_2Cl_2$ (10 mL) were added, and the solution was cooled to 0°C under a nitrogen environment. A mixture of pyridine (0.13 mL, 1.59 mmol, 3 equiv) and MFHBI (115 mg, 0.53 mmol, 1 equiv) in 2 mL of DCM was then added slowly. The reaction mixture was stirred at room temperature and monitored by TLC until completion (~12 hours). The reaction mixture was concentrated to an orange solid under reduced pressure and then purified by column chromatography using a silica column (Hex/EtoAc from 0 to 50%), providing MC-MFHBI as a light-yellow solid (86 mg, 56%).

**$^1$H NMR** (500 MHz, $CDCl_3$): δ 8.24-8.22 (d, *J* = 11.6 Hz, 1H), 7.73-7.72 (d, *J* = 10.7 Hz, 1H), 7.26-7.23 (t, *J* = 8.3 Hz, 1H), 6.99 (s, 1H), 3.94 (s, 3H), 3.19 (s, 3H), 2.38 (s, 3H).

**$^{13}$C NMR** (126 MHz, $CDCl_3$): δ 170.7, 163.9, 155.0, 153.2, 139.9, 139.6, 134.1, 128.7, 124.7, 123.4, 120.0, 56.2, 26.9, 15.9.

**$^{19}$F NMR** (471 MHz, $CDCl_3$): δ -128.2 (m).

**HRMS** (ESI) m/z: Calculated for $C_{14}H_{14}FN_2O_4^+$, $[M + H]^+$ requires 293.0932; found 293.0947.

**AEC-DFHBI** ($^1H$ NMR: 500 MHz, $CDCl_3$)

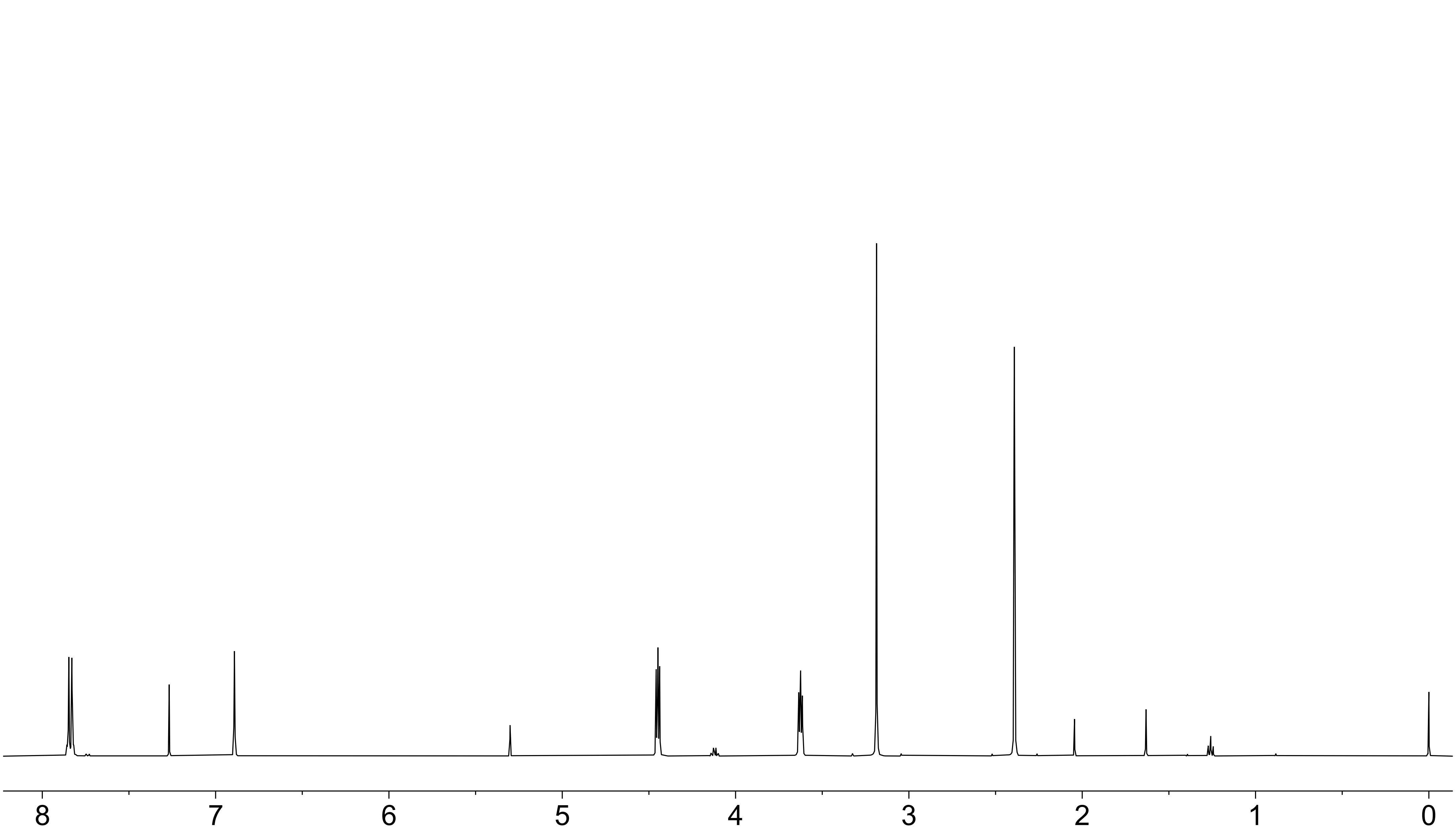

**AEC-DFHBI** ($^{13}$C NMR: 126 MHz, $CDCl_3$)

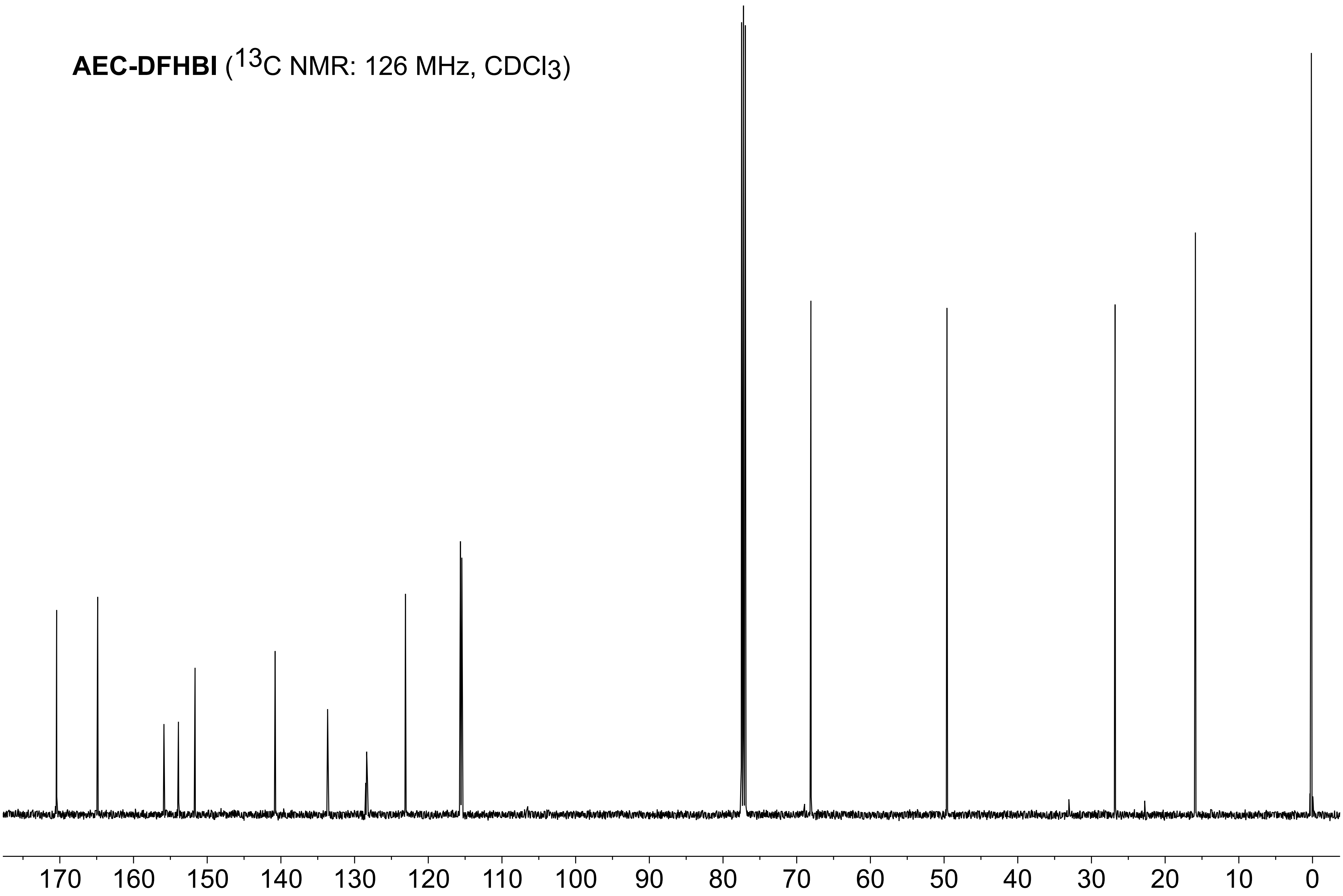

**AEC-DFHBI** ($^{19}F$ NMR: 470 MHz, $CDCl_3$)

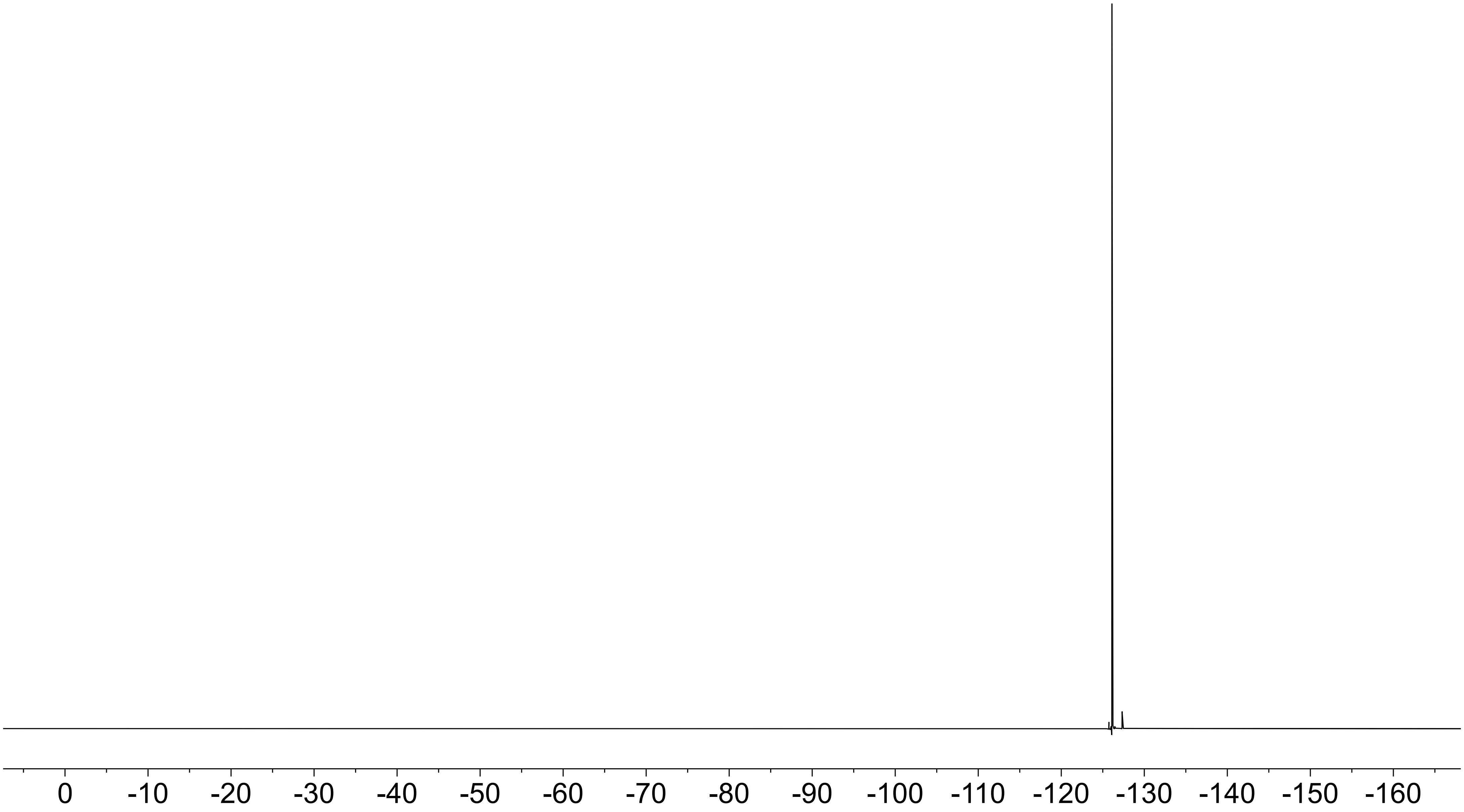

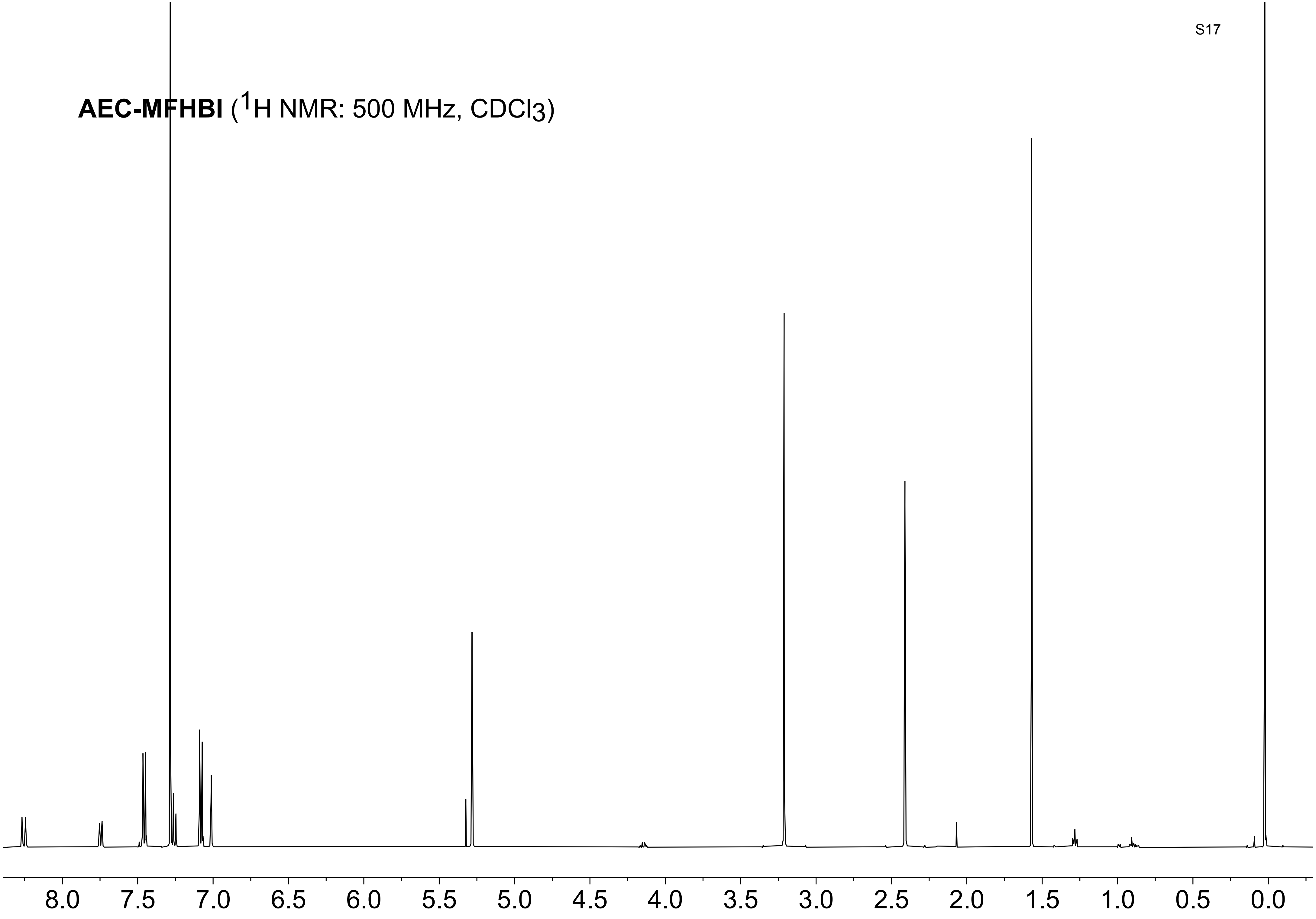
AEC-MFHBI (1H NMR: 500 MHz, CDCl3)
8.0
7.5
7.0
6.5
6.0
5.5
5.0
4.5
4.0
3.5
3.0
2.5
2.0
1.5
1.0
0.5
0.0

**AEC-MFHBI** ($^{13}$C NMR: 126 MHz, $CDCl_3$)

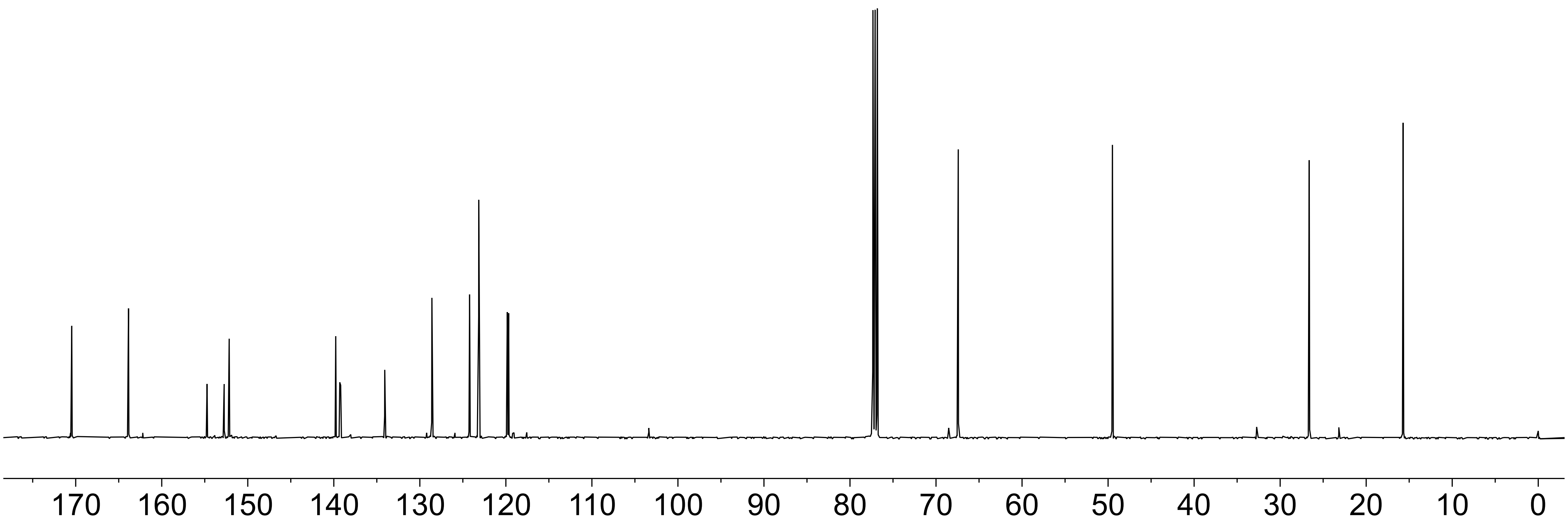

**AEC-MFHBI** ($^{19}$F NMR: 470 MHz, $CDCl_3$)

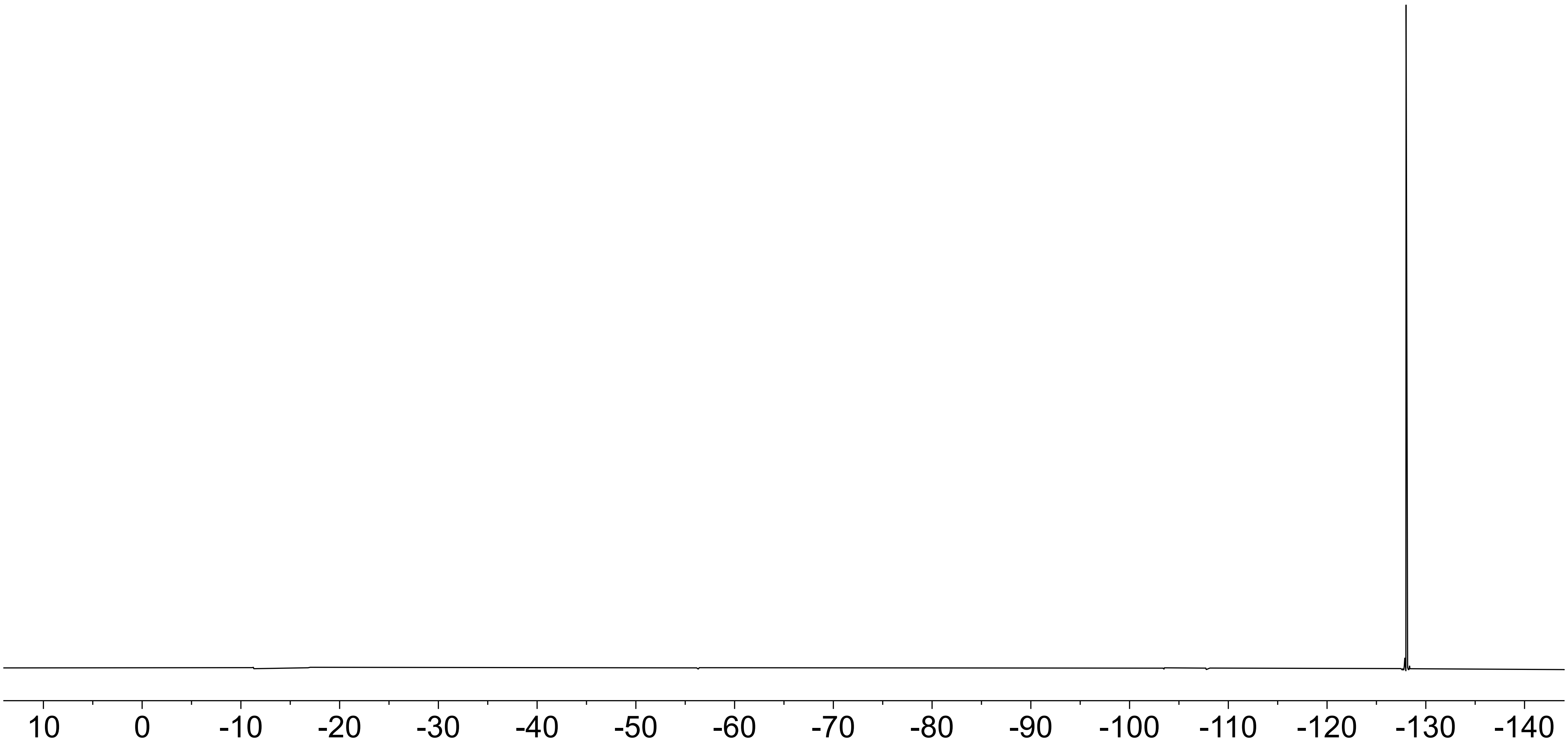

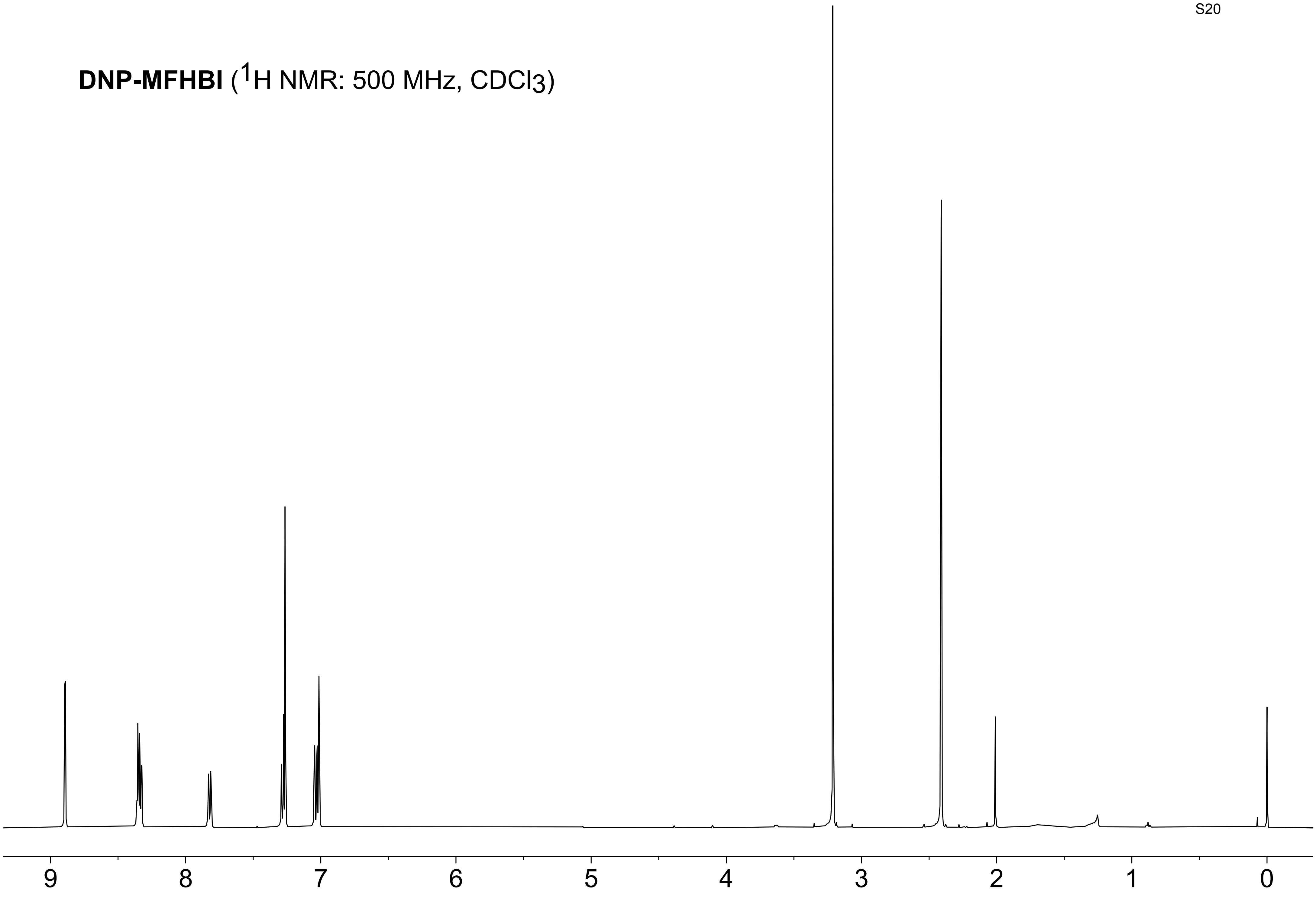
**DNP-MFHBI** ($^1$H NMR: 500 MHz, $CDCl_3$)
9
8
7
6
5
4
3
2
1
0

**DNP-MFHBI** ($^{13}C$ NMR: 126 MHz, $CDCl_3$)

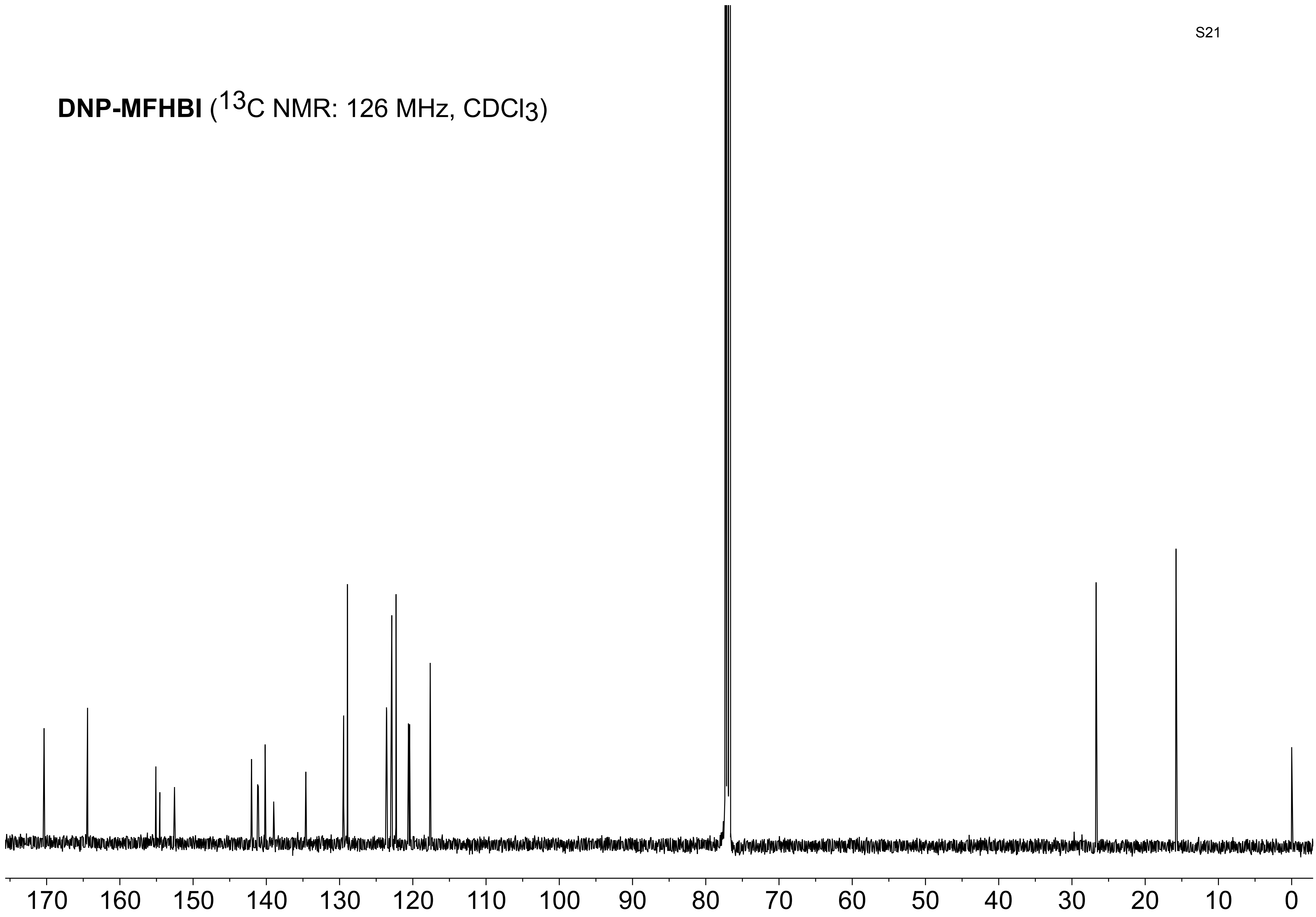

**DNP-MFHBI** ($^{19}F$ NMR: 470 MHz, $CDCl_3$)

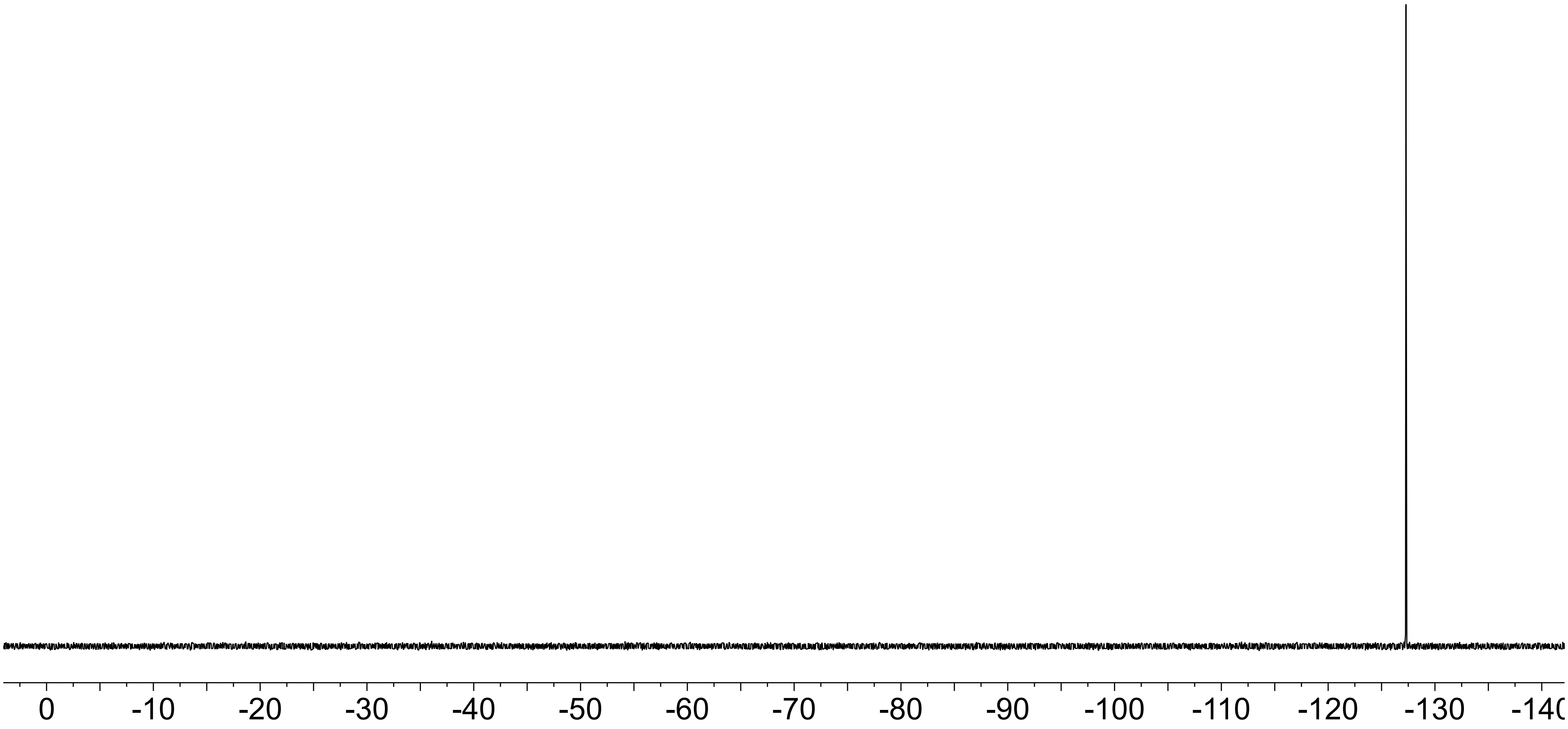

**PyDSBz-MFHBI** ($^1$H NMR: 500 MHz, $CDCl_3$)

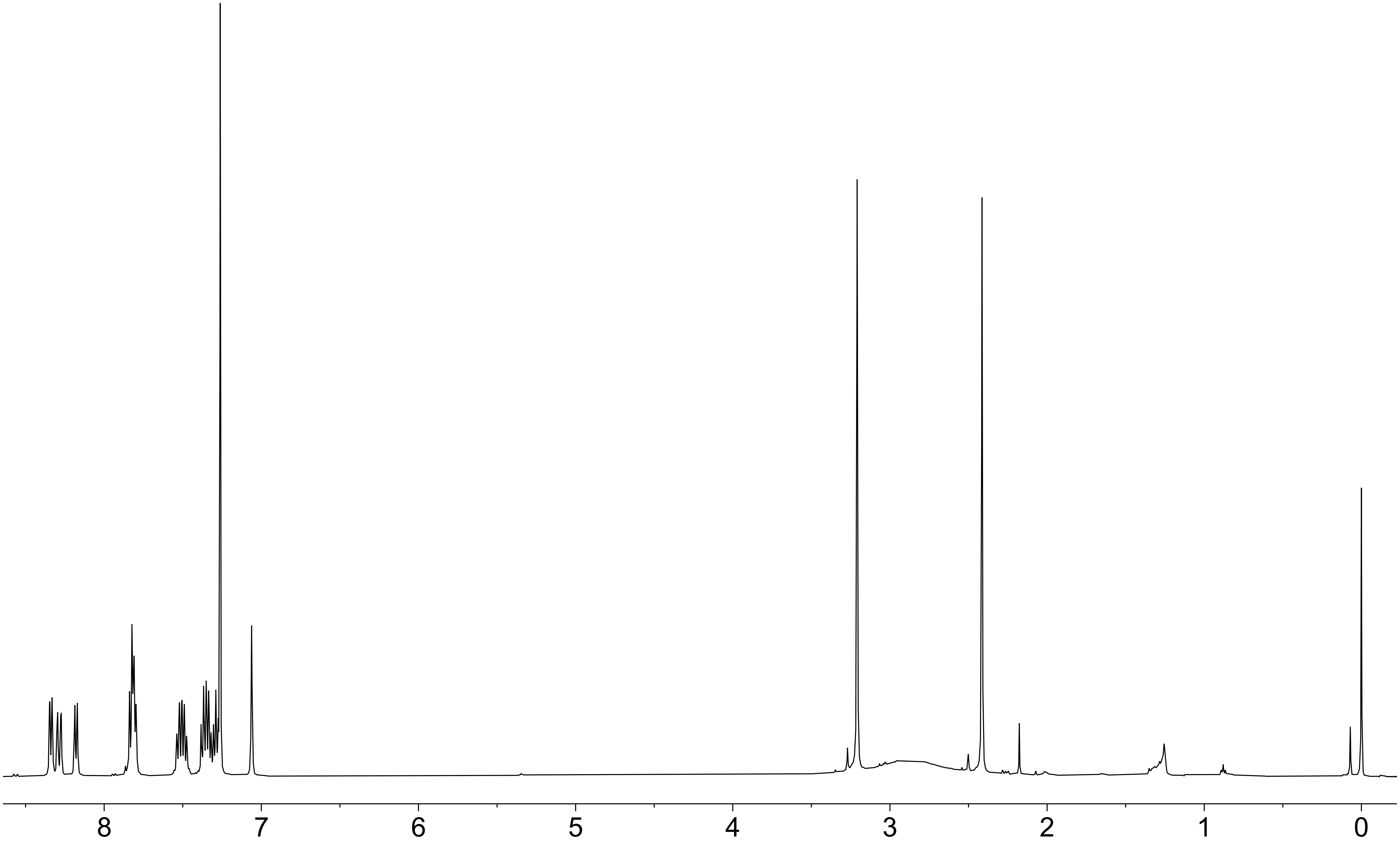

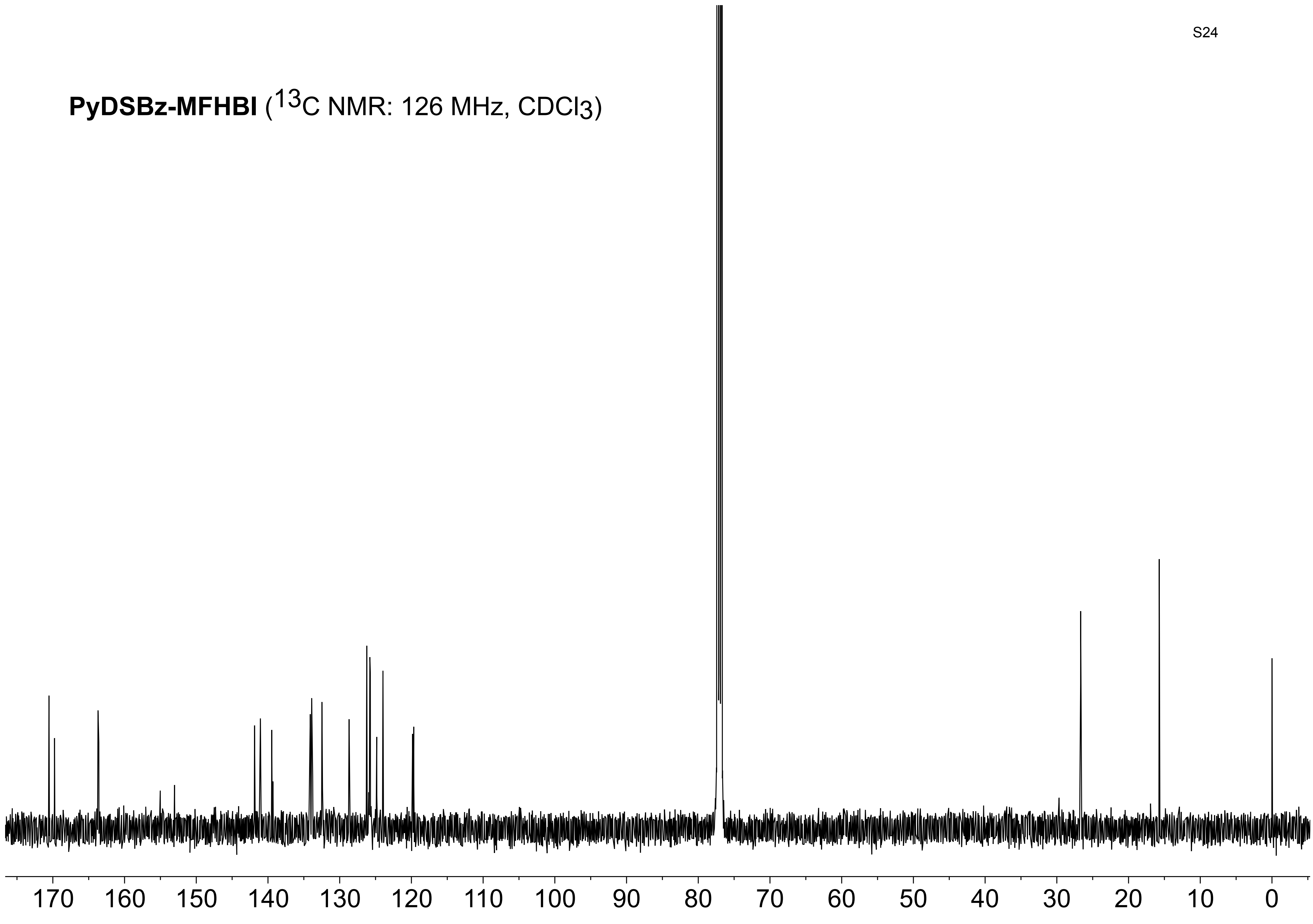
PyDSBz-MFHBI (13C NMR: 126 MHz, CDCl3)
170 160 150 140 130 120 110 100 90 80 70 60 50 40 30 20 10 0

**PyDSBz-MFHBI** ($^{19}F$ NMR: 470 MHz, $CDCl_3$)

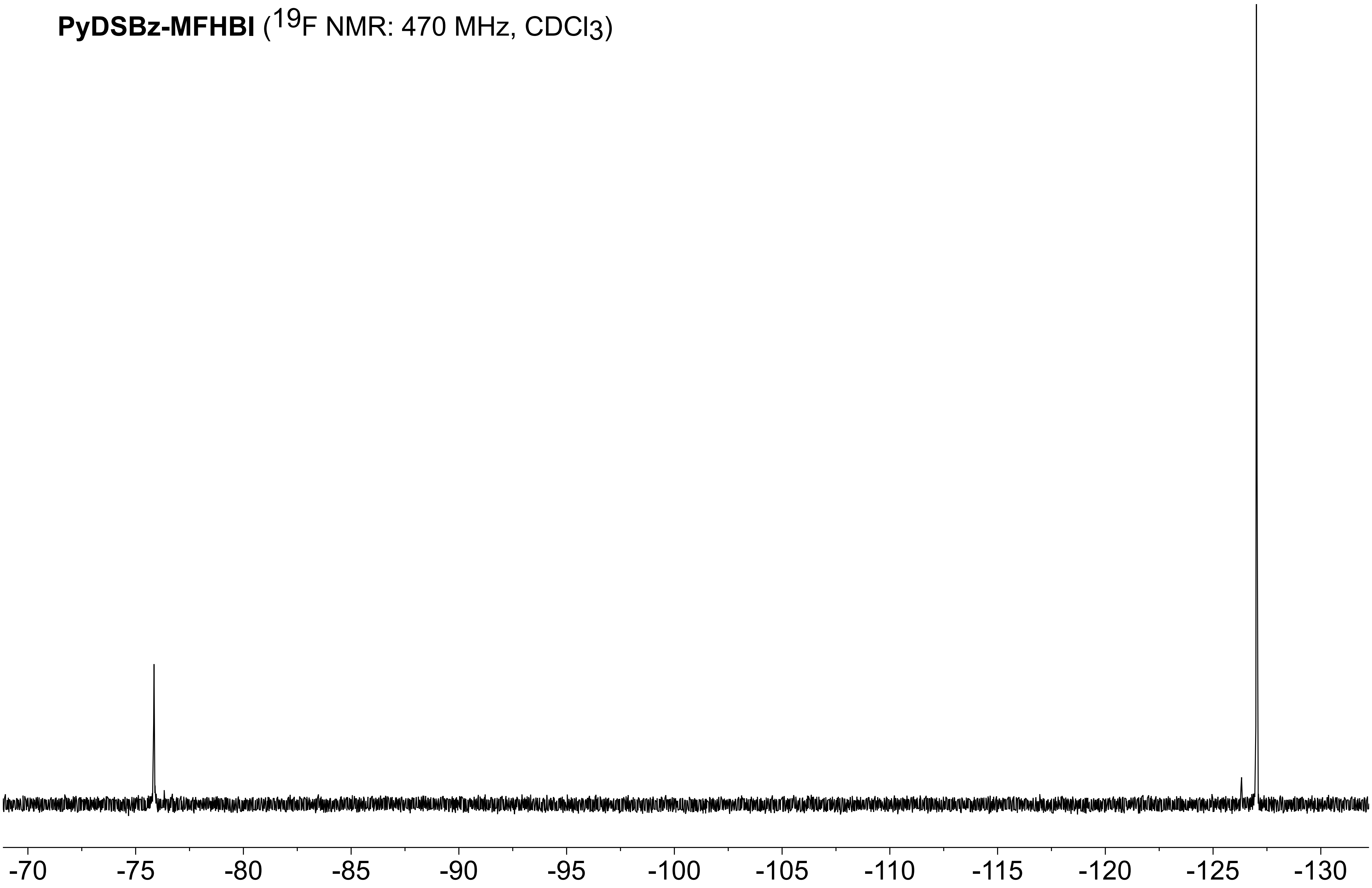

**PyDSBz-DBrHBI** ($^1$H NMR: 500 MHz, $CDCl_3$)

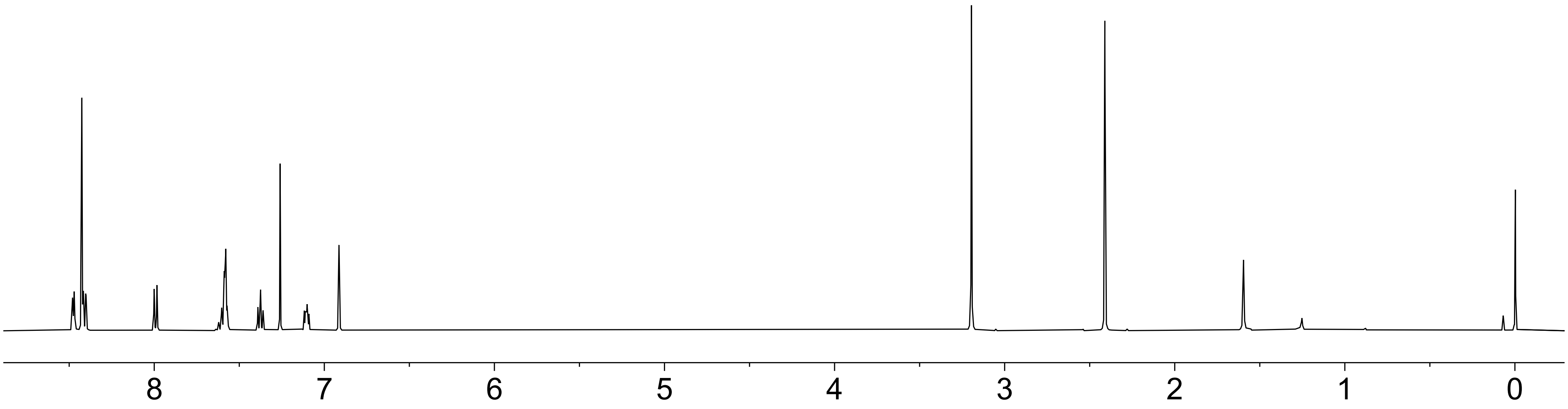

**PyDSBz-DBrHBI** ($^{13}C$ NMR: 126 MHz, $CDCl_3$)

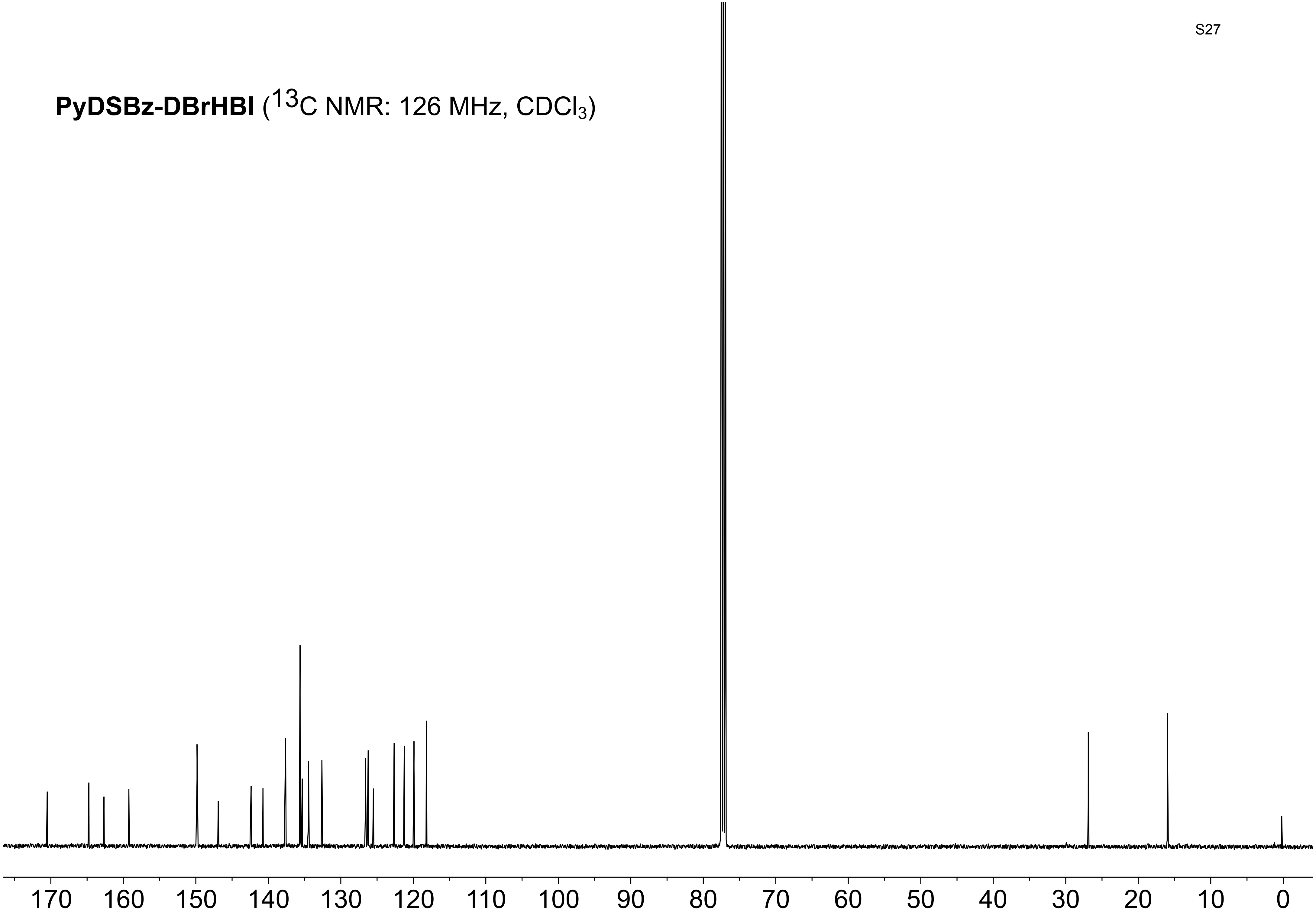

**MBBA-MFHBI** ($^1$H NMR: 500 MHz, d6-DMSO)

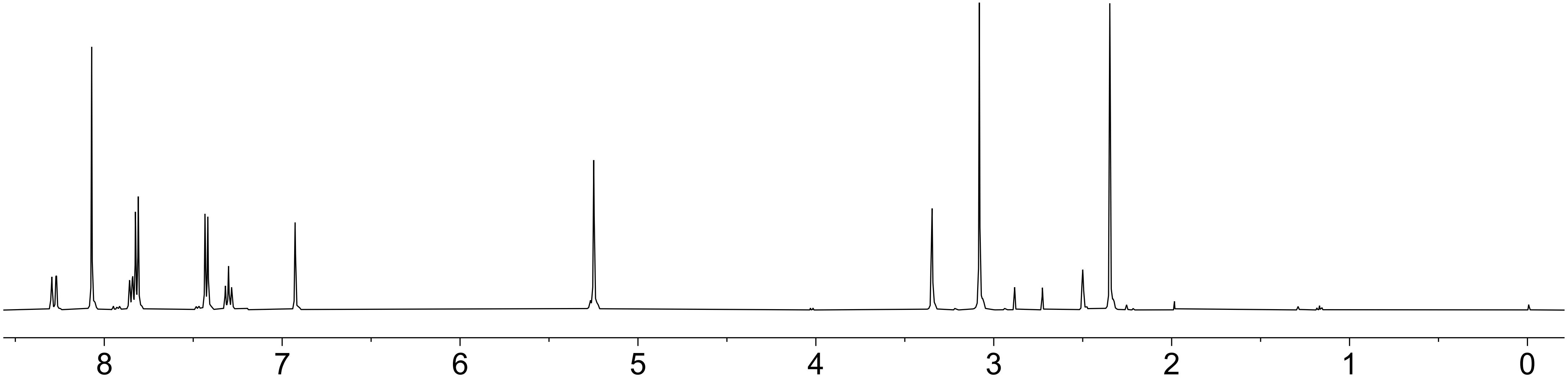

**MBBAPE-MFHBI** ($^{13}C$ NMR: 126 MHz, $CDCl_3$)

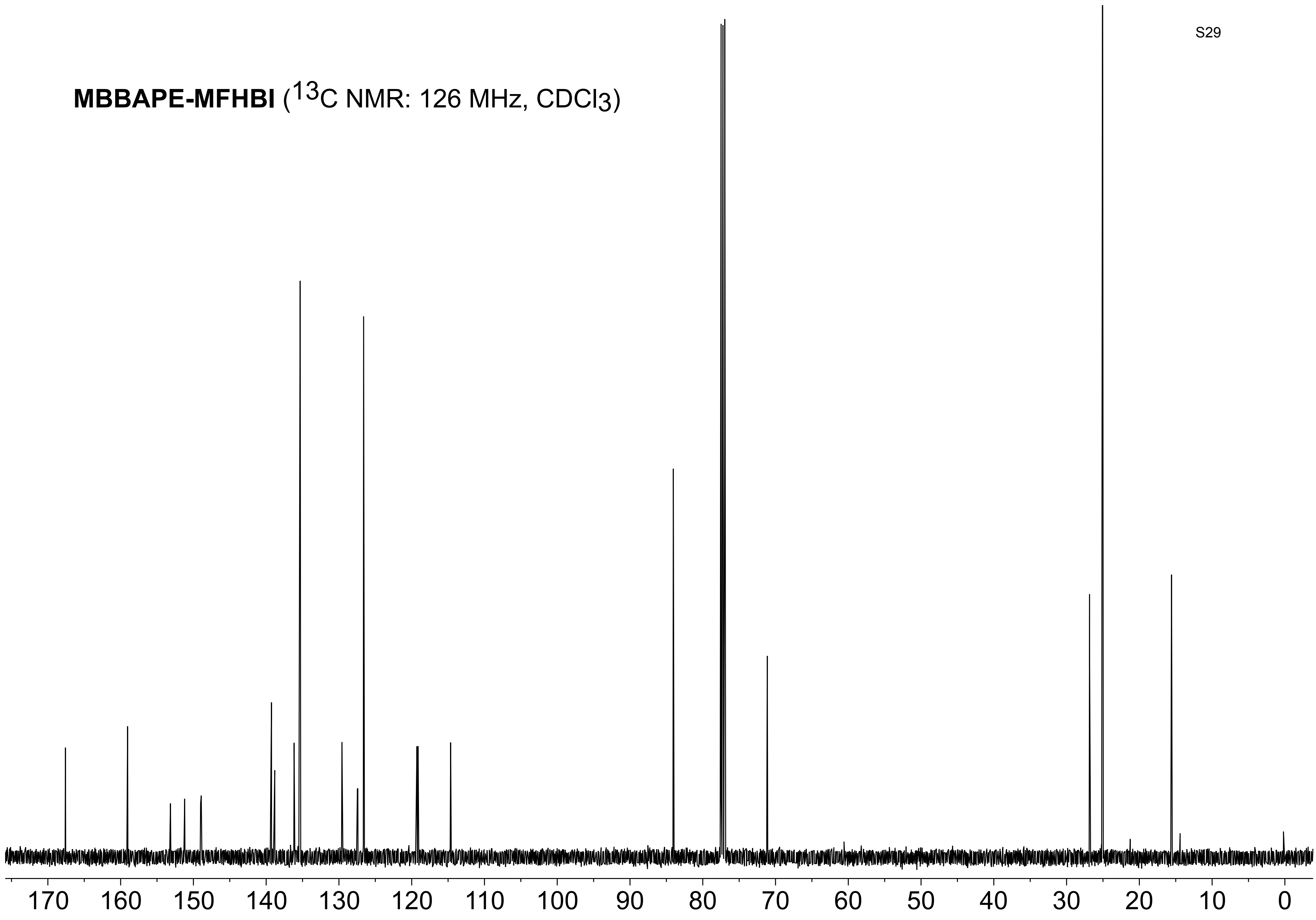

**MBBAPE-MFHBI** ($^{19}F$ NMR: 470 MHz, $CDCl_3$)

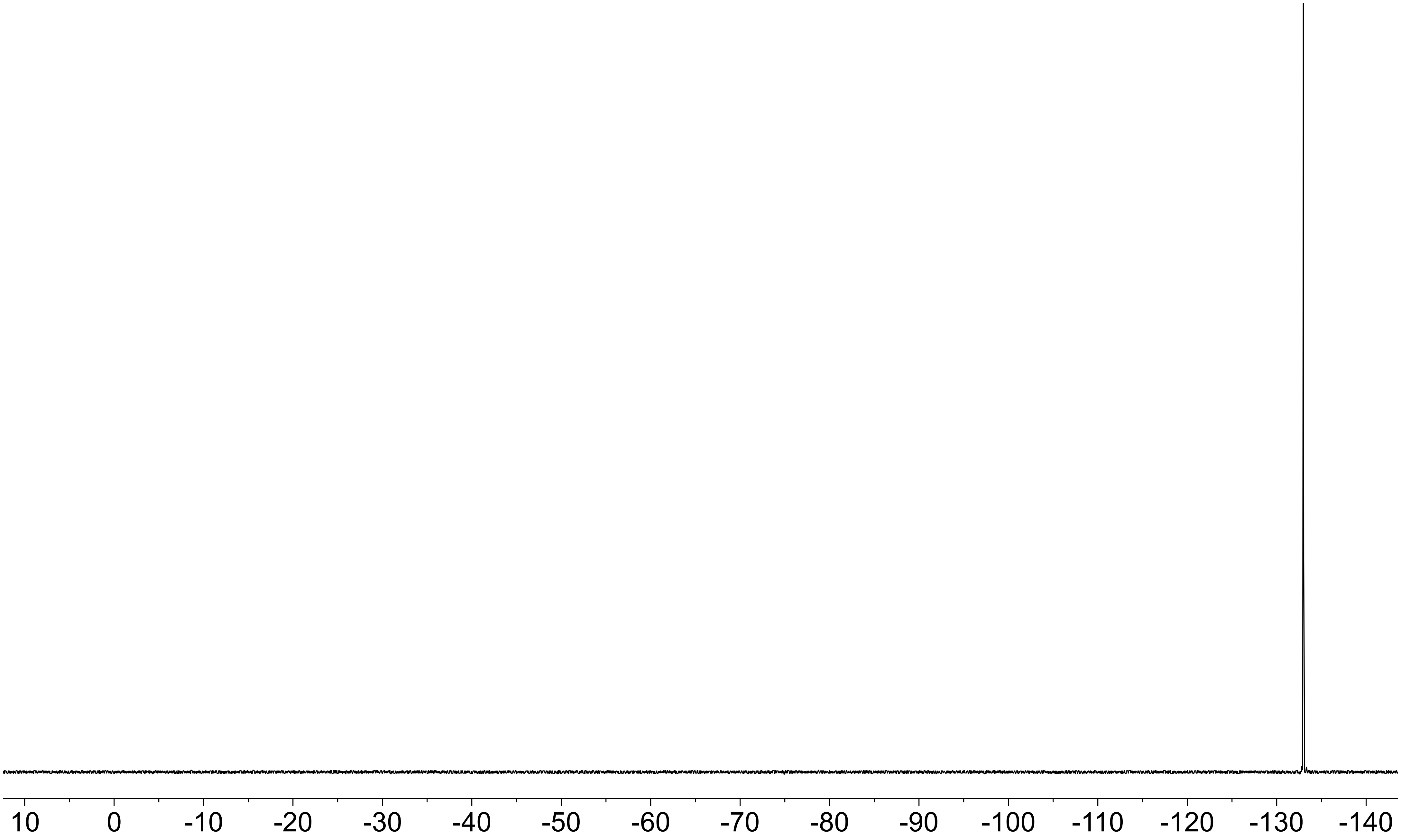

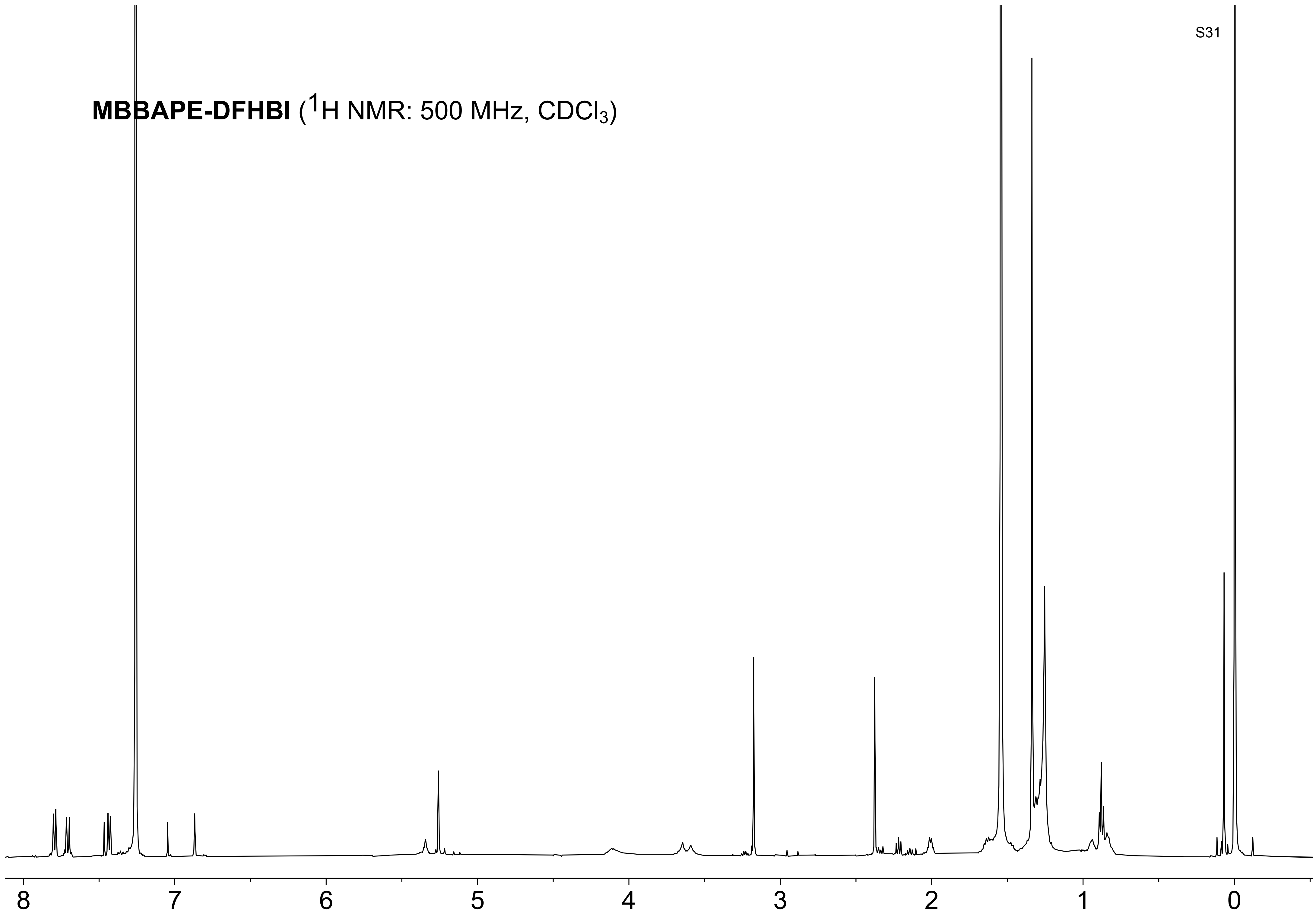
MBBAPE-DFHBI ($^1$H NMR: 500 MHz, $CDCl_3$)
8
7
6
5
4
3
2
1
0

**MBBAPE-DFHBI** ($^{13}C$ NMR: 126 MHz, d6-DMSO)

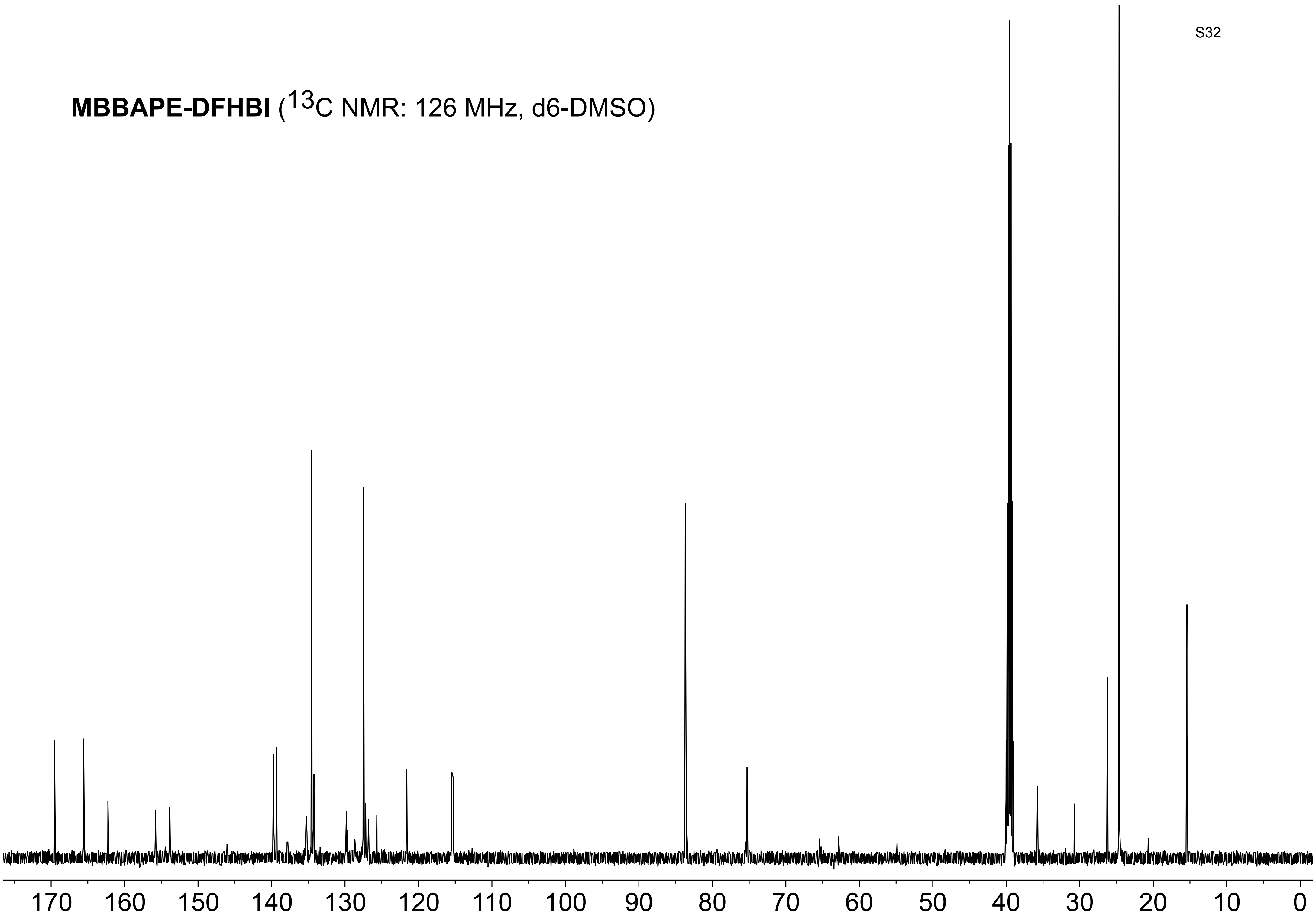

**MBBAPE-DFHBI** ($^{19}$F NMR: 470 MHz, d6-DMSO)

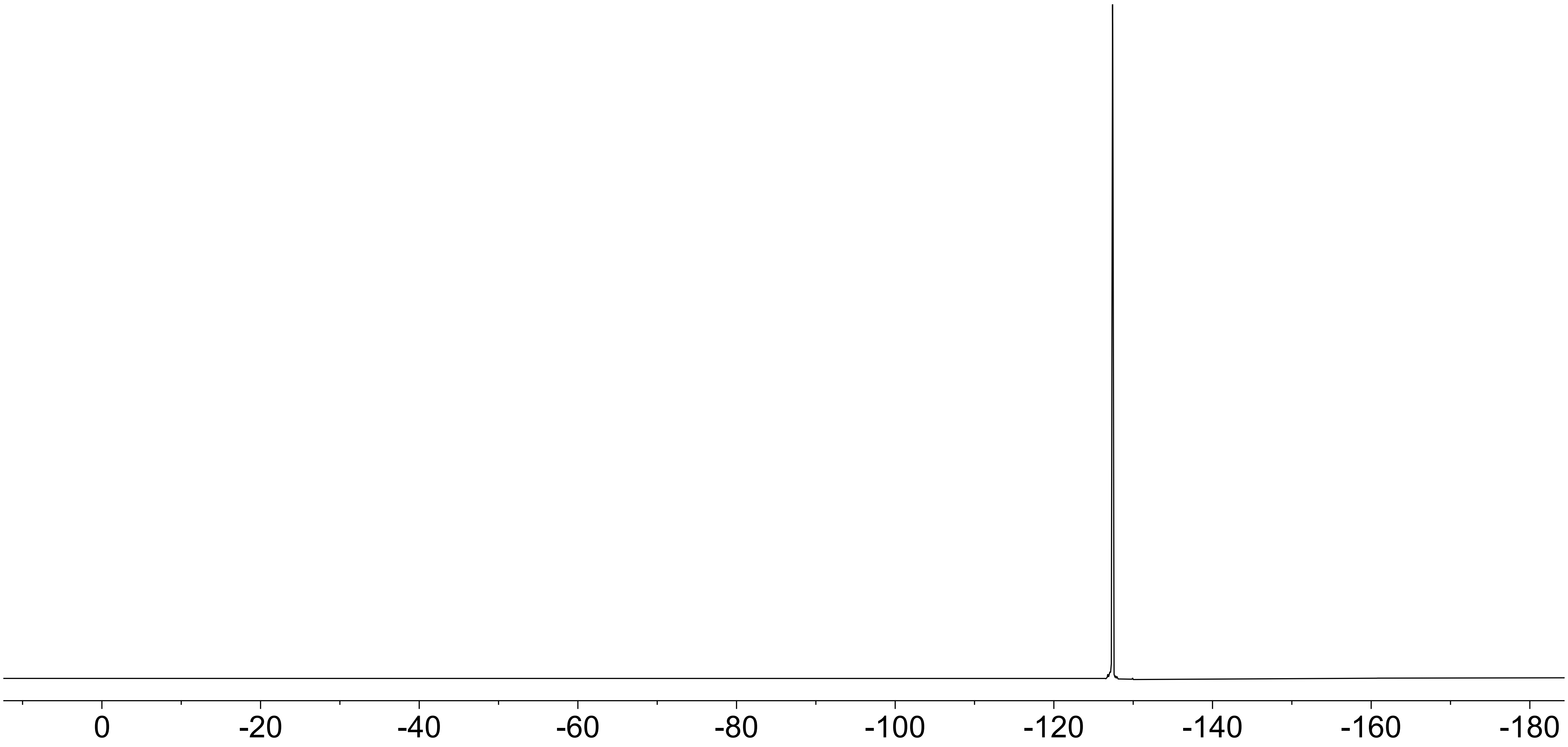

**MBBAPE-MFHBI** ($^1H$ NMR: 500 MHz, $CDCl_3$)

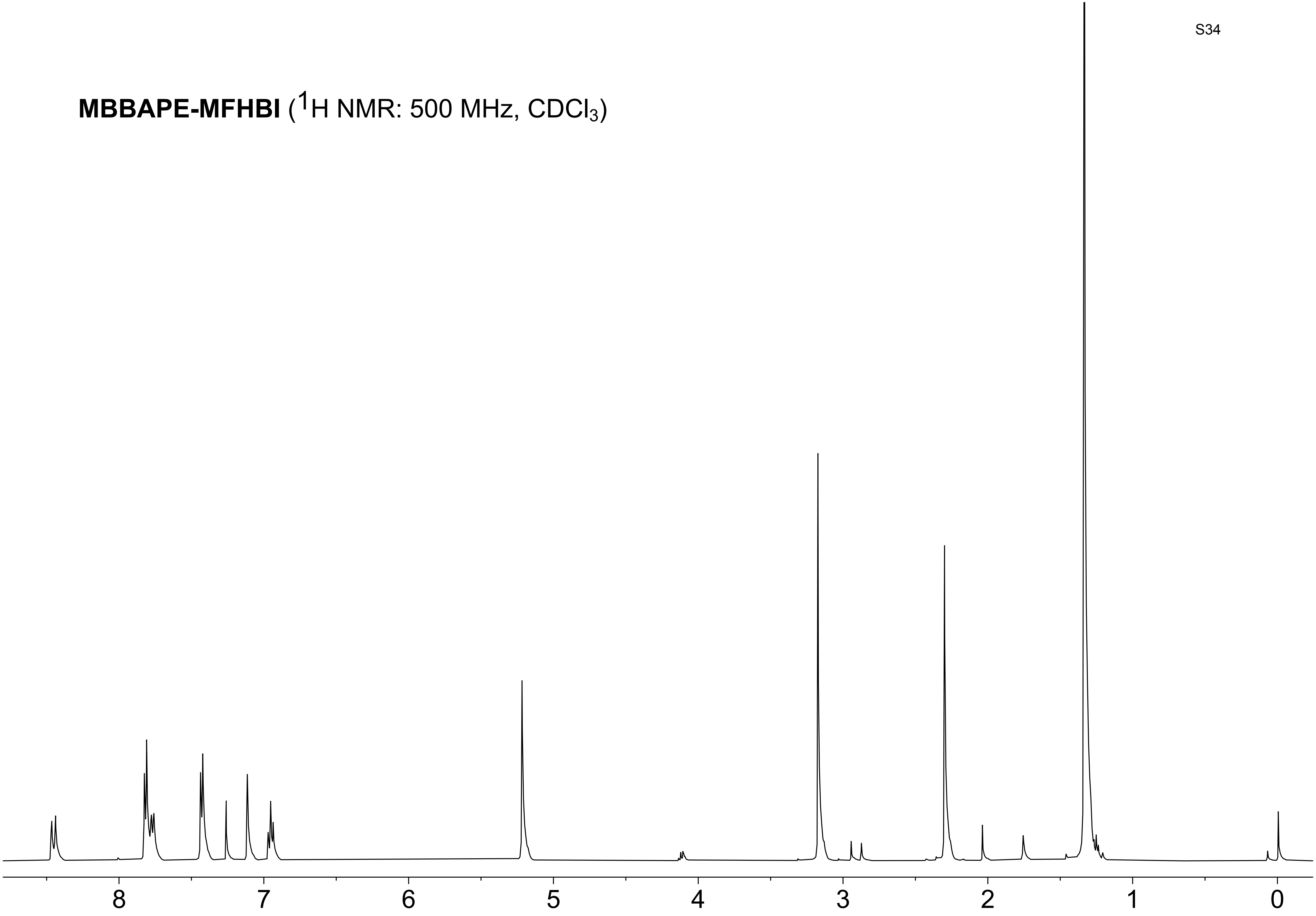

**MBBA-MFHBI** ($^{13}$C NMR: 126 MHz, d6-DMSO)

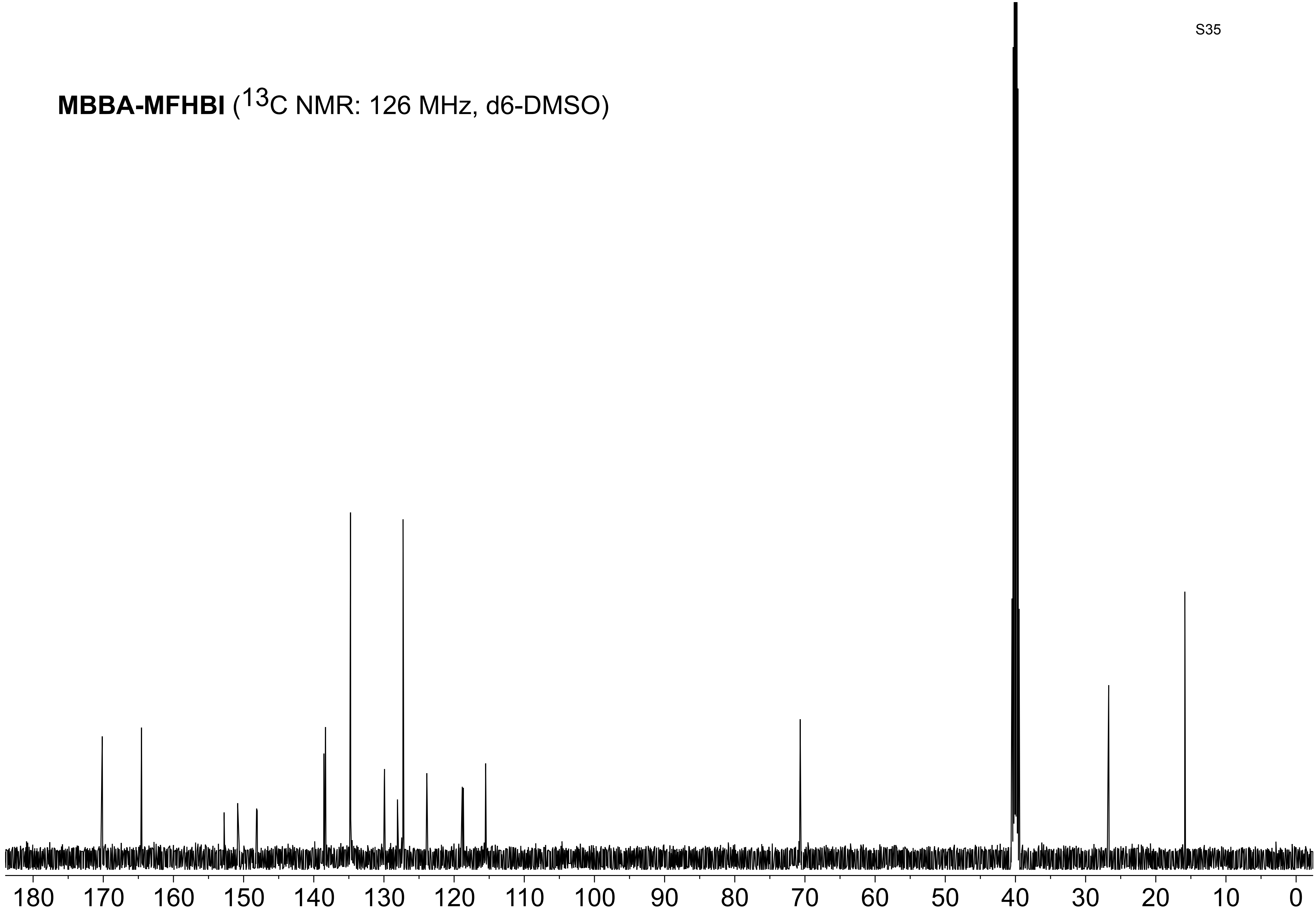

**MBBA-MFHBI** ($^{19}F$ NMR: 470 MHz, d6-DMSO)

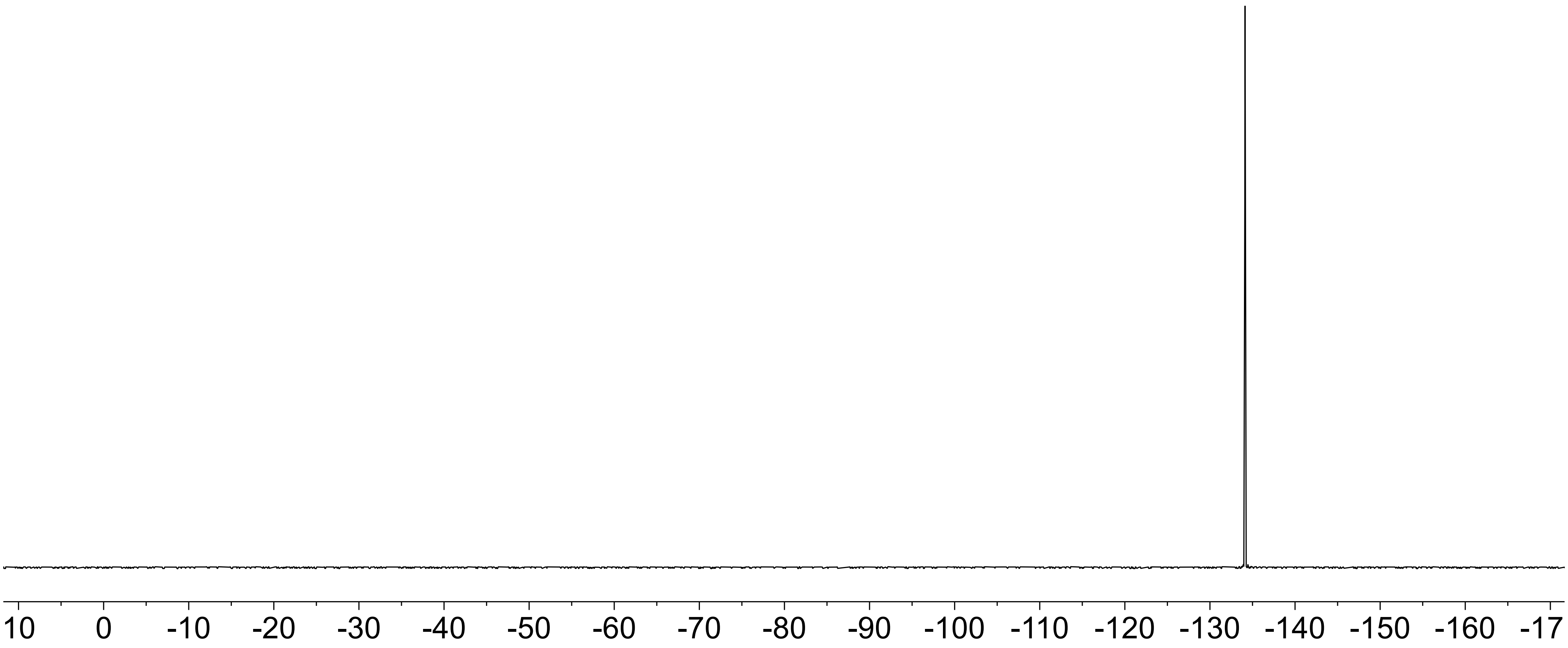

**MC-MFHBI** ($^1$H NMR: 500 MHz, $CDCl_3$)

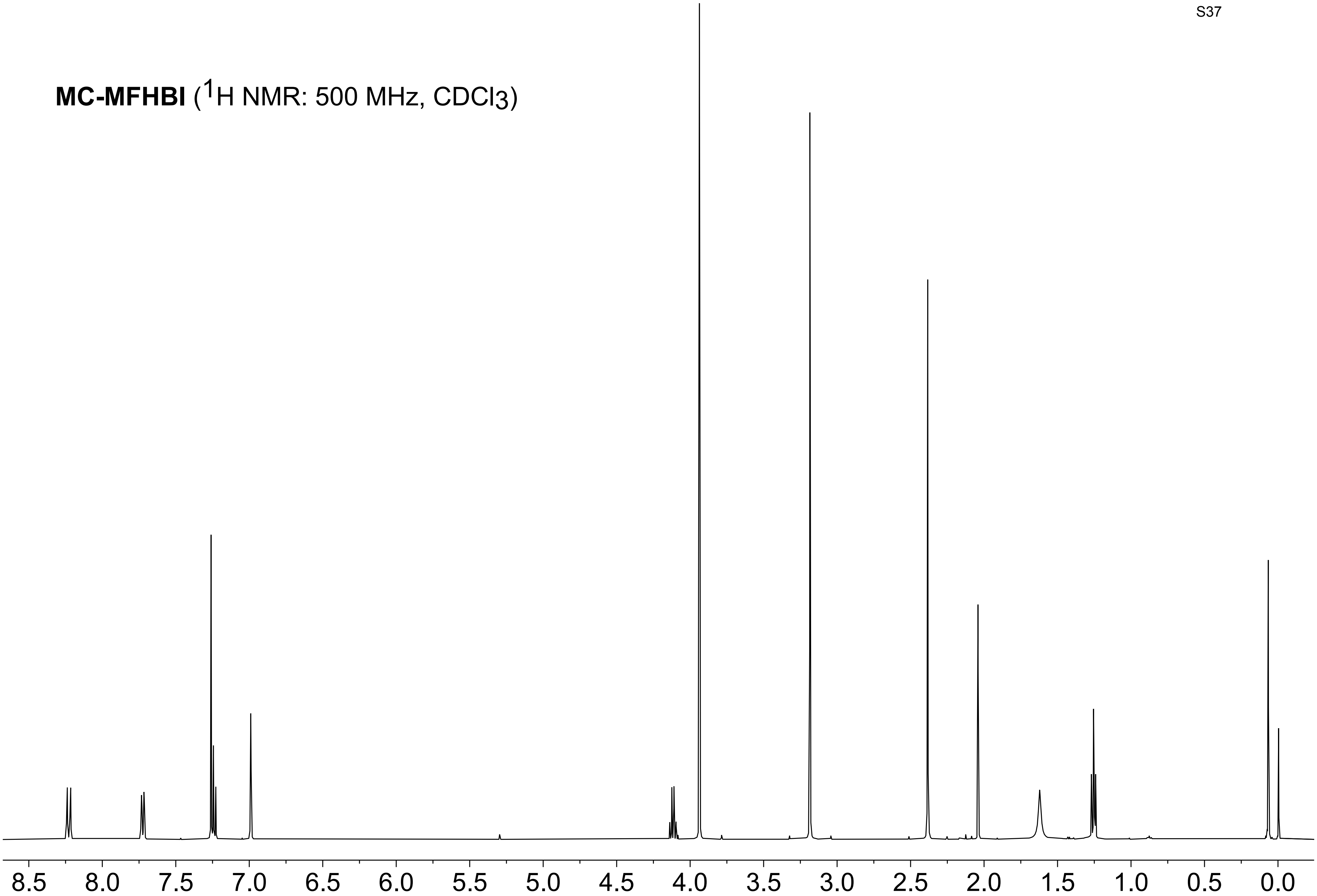

**MC-MFHBI** ($^{13}C$ NMR: 126 MHz, $CDCl_3$)

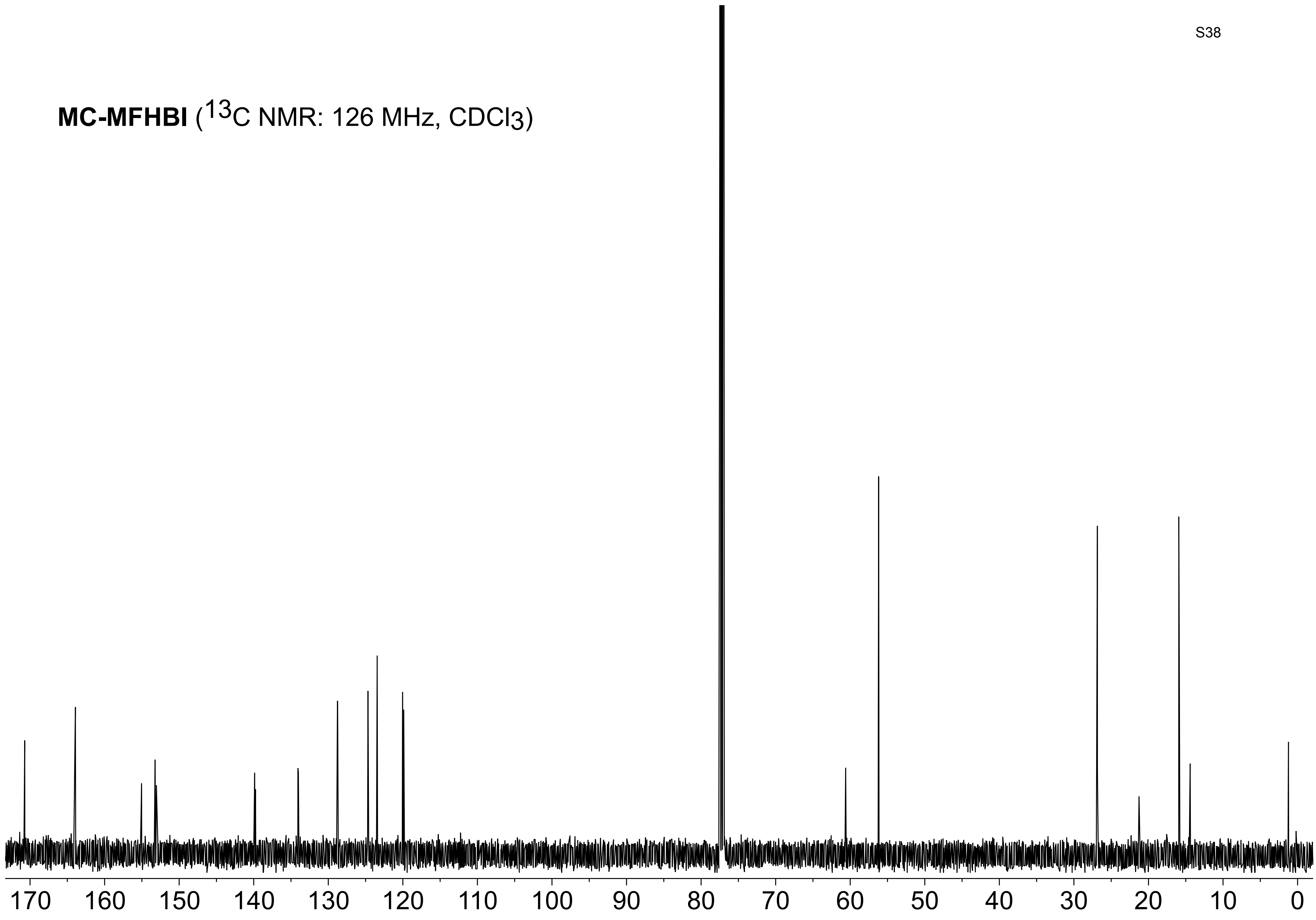

**MC-MFHBI** ($^{19}$F NMR: 470 MHz, $CDCl_3$)

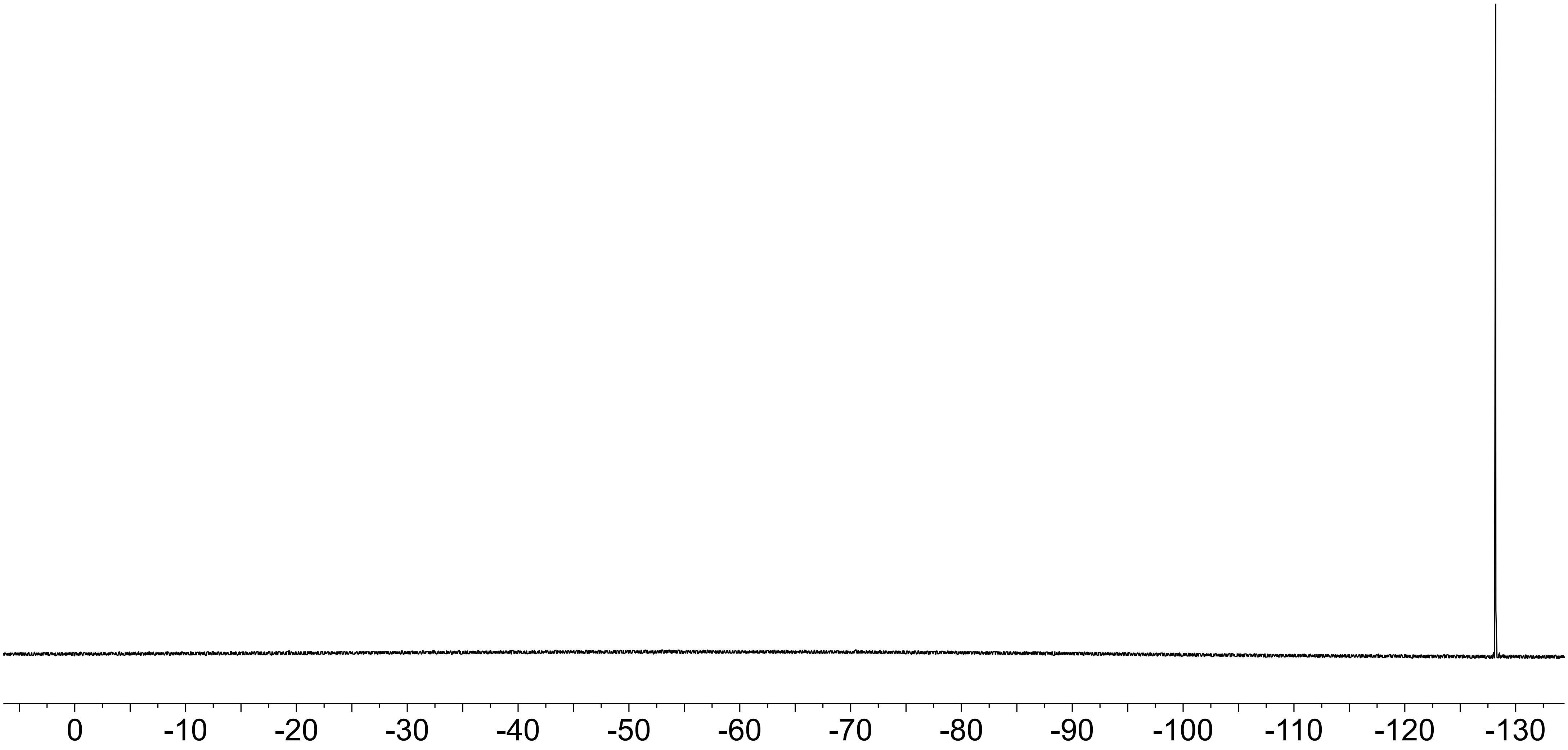

### Docking Simulation Method

Initial structures of the ligands to predict potential binding targets were generated and optimized in Gaussian16[1] using the DFT/B3LYP/6-31G basis set. Optimized input structure files of ligand and aptamer for AutoDock Vina[2,3] were coverted in the final pdbqt file format using AutoDockTools (version 1.5.7)[4] with only polar hydrogens included in the input files. All possible torsions in the ligand structure were included to maximize potential conformational changes in the binding site. Search box sizes were chosen to be large enough to encompass the binding sites of 4TS2 but otherwise minimal to optimize search time. Box size was set as 12x12x12 $Å^3$ and centered around the binding site (Coordinates: center_x = -11.072, center_y = -0.006, center_z = -15.492). Docking exhaustiveness was set to 8 to ensure a significant sampling of poses were generated but only the highest scored pose was chosen for analysis. The free binding energies were generated by in-built linear regression model in AutoDock Vina.

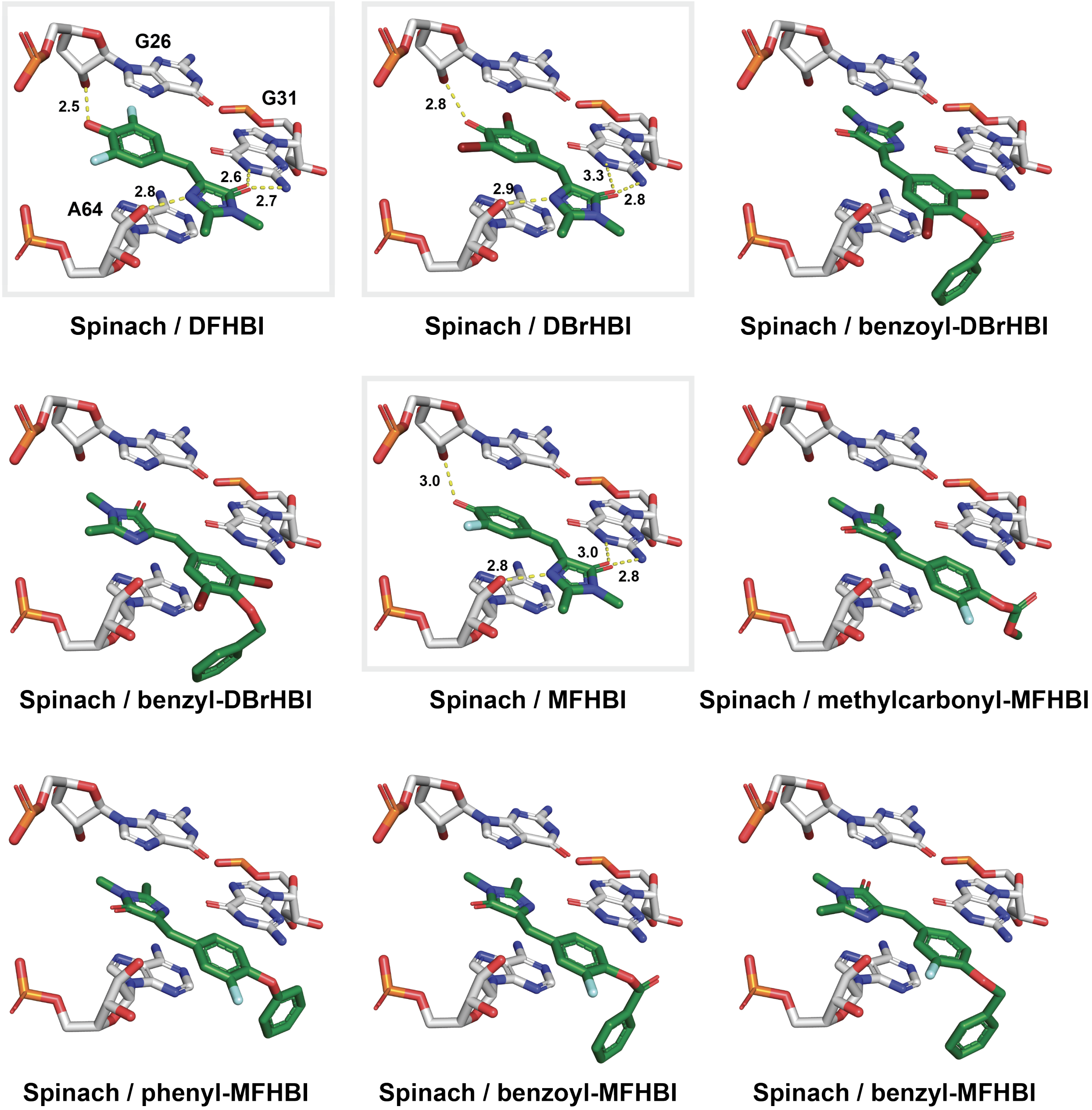


**Figure S1. Docked poses: binding mode of 4-*O*-aryl-caged small molecules derived of DBrHBI and MFHBI.** The crystal structure of Spinach-DFHBI (PDB:4TS2) near the binding site shows the key aptamer-ligand hydrogen-bonding interactions. Molecular docking of the 4-*O*-aryl-caged DBrHBI and MFHBI were performed using AutoDock Vina.

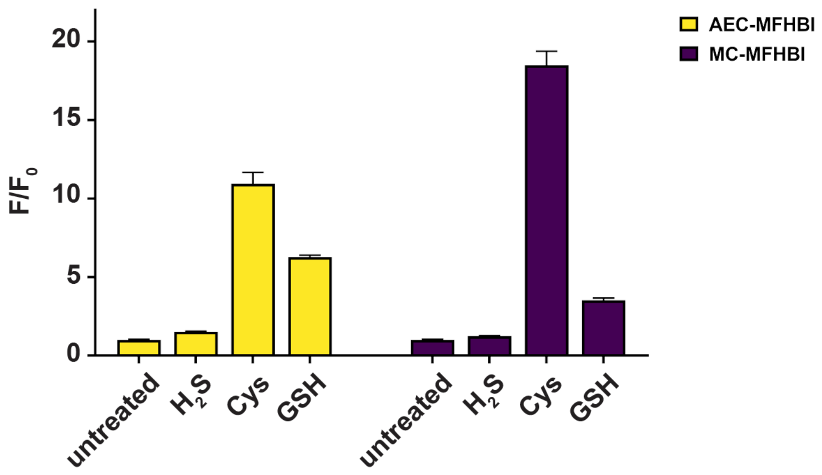


**Figure S2. RSS selectivity of AEC-MFHBI and MC-MFHBI.** Plotted are signal-to-background ratio (*$F/F_0$*) of AEC-MFHBI and MC-MFHBI upon incubation with 2 mM $H_2S$, Cys, or GSH for 2 hours.

### Determination of $pK_a$ of DBrHBI and MFHBI

The $\lambda_{max}$ emission of DBrHBI dissolved in a solution at pH 8.35 was determined to be 480 nm. The absorbance of DBrHBI (10 µM) at 480 nm was measured in 40 mM Britton-Robinson buffers (boric acid/phosphoric acid/acetic acid, 1:1:1 molar ratio) with pH values of 1.90, 4.69, 5.15, 6.03, 7.68, and 8.35. Measured values were used to plot a sigmoidal regression model (Figure S1 and S2). The half-maximal pH value, representing $pK_a$, was determined as 5.22 ± 0.15.

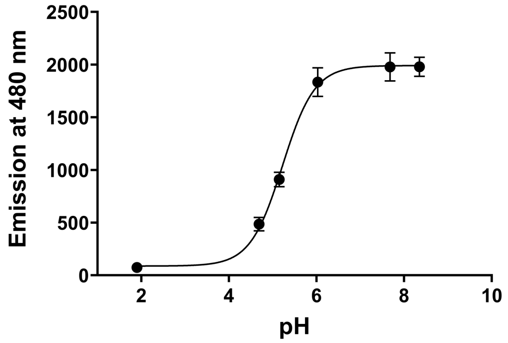


**Figure S3.** Emission (480 nm) of DBrHBI vs. pH. For each absorbance measurement, a 10 µM solution of DBrHBI was prepared in Britton-Robinson buffers (pH 1.90–8.35). $R^2$ = 0.989. Error bars represent standard deviation, *n* = 3.

A similar protocol was followed to determine the $pK_a$ of the MFHBI. The maximum absorbance of MFHBI dissolved in a solution at pH 10.44, which should primarily contain the aryloxide anion form of MFHBI, was determined to be 422 nm. The absorbance of MFHBI (10 µM) at 422 nm was measured in 40 mM Britton-Robinson buffers (boric acid/phosphoric acid/acetic acid, 1:1:1 molar ratio) with pH values of 5.17, 6.03, 6.93, 7.68, 8.79, and 9.90. Measured values were used to plot a sigmoidal regression model (Figure S1). The half-maximal pH value, representing $pK_a$, was determined as 6.78 ± 0.15, which supports the previous report of 6.9.[5]

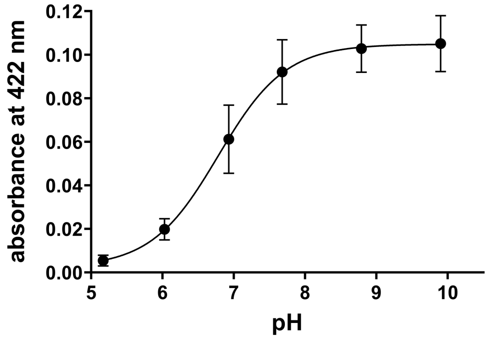


**Figure S4.** Absorbance (422 nm) of MFHBI vs. pH. For each absorbance measurement, a 10 µM solution of MFHBI was prepared in Britton-Robinson buffers (pH 5.17–9.90). $R^2$ = 0.948. Error bars represent standard deviation, *n* = 3.

### Fluorescence Measurement Protocols

**$H_2S$ detection:** 100 μM stock solution of Baby Spinach was first diluted into 2 μM using the buffer. PyDSBz-DBrHBI stock solution was prepared at 8 mM in DMSO, and diluted into 11.1 μM by buffer. 50 μL of the RNA, 45 μL of PyDSBz-DBrHBI, and 5 μL of the corresponding reactive sulfur species at 10 mM will be mixed in a capped PCR tube to prevent $H_2S$ leakage. The solutions were transferred to 96-well plates at different time points (0, 10, 20, 30, 40, 50, 60 min) for fluorescence measurement (ex/em 446/499 nm). Final samples contained 1 μM BabySpinach, 5 μM PyDSBz-DBrHBI, and 500 μM RSS (Figure 4A).

**$H_2O_2$ detection:** 4 μL of MFHBI or MBBA-MFHBI solution (125 μM) in DMSO and 1 μL of the stock solution of the RNA (100 μM) were added to a 96-well plate containing Baby Spinach buffer at pH 7.4 (50 mM HEPES, 100 mM KCl, and 5 mM $MgCl_2$), for a total volume of 100 μL. At this final volume, the concentrations of MFHBI or MFHBI derivative and RNA were 5 μM and 1 μM, respectively. Fluorescence intensity (ex/em 470/510 nm) of the samples were measured on a microplate reader (Molecular Devices SpectraMax iD3). For $H_2O_2$ reactivity assays, final concentrations were set to be 100 μM. Measurements were taken at 1, 2, 4, 6, 8, 10, 12, 15, 20, 25, 30, 40, 45, 50, and 60 minutes time interval. RFU vs time data were plotted using GraphPad Prism software (Figure 4D).

**Limit of Detection:** Samples were prepared using the steps mentioned above, except the corresponding redox agents were added at serial dilutions within a range of 0–1 μM. Fluorescence intensities were measured after 1 hour of incubation. Samples at each concentration were repeated for three replicates, and one-tailed Student’s t-test was used to evaluate whether the fluorescence intensities at each redox agent concentration is statistically higher than that at 0 μM (Figures 5C and 5F).

### Protocols for Cell Studies

**Design and cloning of 3xBaby Spinach in *E. coli*:** Custom-made plasmid designed to express a single motif with three continual baby spinach binding sites was purchased from Twist Bioscience. The resulting pET-21(+)-3xBaby Spinach plasmid contained a 3xBaby Spinach gene sequence, ampicillin resistance gene and an IPTG-inducible T7 promoter. This plasmid was first sequence confirmed and then transformed into BL21-star-(DE3) chemically competent cells using the heat shock method and were selected on the LB-agar plate with Amp (100 μg/mL).

**Preparation of *E. coli* for the live cell imaging:** 5 mL LB containing Amp (100 μg/mL) was inoculated with a single isolated colony and was grown overnight at 37 °C in incubator shaker. The overnight culture was used to inoculate (1:100 dilution) 100 mL LB containing Amp (100 μg/mL) and grown at 37 °C in incubator shaker until the optical density (OD 600) of the culture reached 0.4. RNA expression was induced by the addition of IPTG to a final concentration of 1 mM and culture was further shaken at 37 °C for 2 hours. Cells were harvested by centrifugation for 5 minutes at 3000x g and washed with 100 mL Hanks' Balance Salt Solution (HBSS), centrifuged, and the liquid decanted. Cell pellet was resuspended in M9 media solution containing either 100 μM MFHBI, DBrHBI (for positive controls) or 100 μM MBBA-MFHBI, PyDSBz-DBrHBI (experiment) and incubated for 15 minutes at 37 °C to allow for the uptake of probes by the cells. Culture was centrifuged and media was decanted. Bacteria pellet was resuspended into the media containing either $H_2O_2$ (100 μM) or $Na_2S$ (2mM). 1 ml culture was taken at every 15 minutes interval for up to 60 minutes to observe the kinetics of the fluorescence generation in the cells. Bacteria were centrifuged, the media decanted, then resuspended in 500 μL of 5 μg/mL FM4-64FX in HBSS and incubated at room temperature for 2 min. A final washing step with HBSS was carried out and the resulting pellet was resuspended in 20 μL HBSS and 3 μL was spread on a microscopy coverslip and put on the glass slide. Edges of the coverslip on glass slide was sealed using clear nail polish. Live cell imaging was conducted on a Leica SP8 tauSTED Confocal Microscope equipped with a Tokai Hit stage-top incubator.

### Fluorescence imaging quantification using ImageJ

**Step 1:** Covert RGB format figure to an 8-bit format figure.

1. Download and open ImageJ Fiji software.
2. Click “File” > “Open”.
3. Click on the desired composite image.
4. Click “Image”, select “Type” > “8-bit”. This image should now be converted to a gray colored image.

**Step 2:** Creating the threshold and outlining the boundaries of the cells.

1. Click “Image” > “Adjust” > “Threshold”. A new window should pop up, adjust the minimum and maximum ranges based on the data selected on the image, adjust the values such that the maximum number of cells are selected, click “Apply”.
2. Click “Process” > “Binary” > “Fill Holes”. This should fill out the cell outlines, which is a crucial step; otherwise the final outlines will be incorrect.
3. Click “Process” > “Binary” > “Watershed”. This breaks each cell into separate objects.
4. Click “Analyze” > “Analyze Particles”. A new window should pop up.
5. Put the values in the “Size” option as “100-Infinity”, Under the “Show” option, select “Outlines” from the drop-down menu. Also select “Add to Manager” and click “OK”.
6. Save “Outline drawing figure” and “ROI” on the computer.

**Step 3:** Transfer the outlines of the composite figure onto a single channel image.

1. Click “File” > “Open”.
2. Select the same figure again.
3. Click “Image” option, select “Color” > “Split Channels”. This should split the composite image into separate single channel images. Select the desired channel (green for our experiment purposes) and close the rest of the images.
4. Go to “ROI Manager” window, unselect and select the “Show all” box. This should allow all the outlines to be transferred to the single channel image.
5. Click on the “Measure” icon within the “ROI Manager” window. A new window with all statistical values should pop up. Copy the data and transfer it to an Excel file.
6. Repeat this analysis for rest of the data.

**Step 4:** Sorting and plotting the data for statistical analysis.

1. Sort the mean fluorescence values from largest to smallest.
2. Select the top performing cells (in our case, we selected 50 cells with highest fluorescence output) from each replicate of a defined time point (0, 15, 30, 45, or 60 min).
3. Copy the data to a GraphPad Prism sheet and plot a “scatter dot plot” with mean as a middle line and standard deviations of the mean as error bars.

**Sequence and Proposed 2-Dimensional Structure of 3xBaby Spinach**

GCCCGGAUAGCUCAGUCGGUAGAGCAGCGGCCGGAUGUAACUAAGGACGGGUCCGGAC
GCAAGGACGGGUCCGACCGAAAGGACGGGUCCAAUGGUGGAAACACCAUUGUUGAGUA
GAGUGUGAGUCGGUCGUUGAGUAGAGUGUGAGGCGUCCGUUGAGUAGAGUGUGAGAG
UUACAUCCGGCCGCGGGUCCAGGGUUCAAGUCCCUGUUCGGGCGCCA

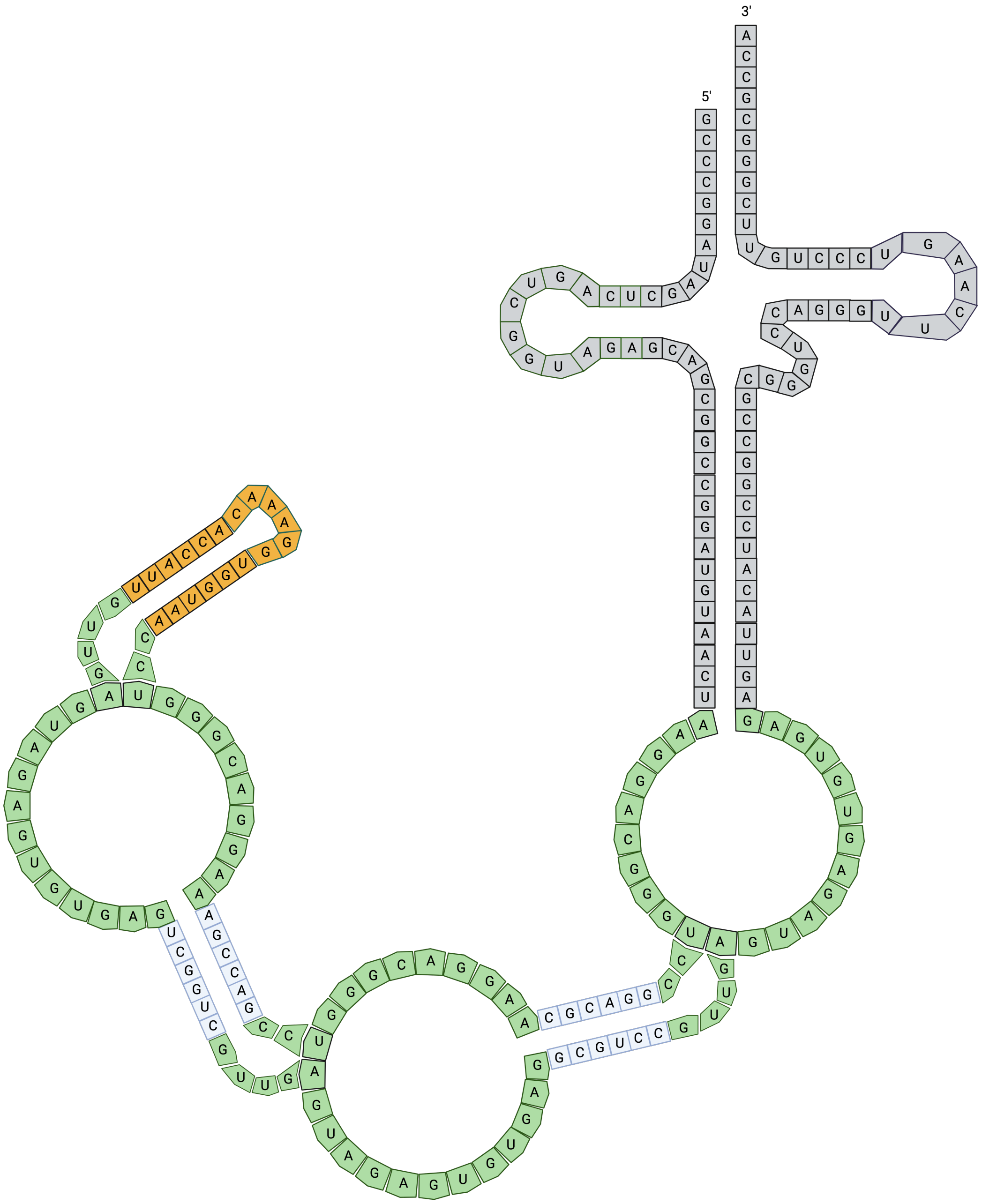


**Figure S5.** 2-Dimensional rendering of the proposed folded 3xBaby Spinach construct. Created using Biorender.

**Plasmid Map of pET-21-3xBaby Spinach**

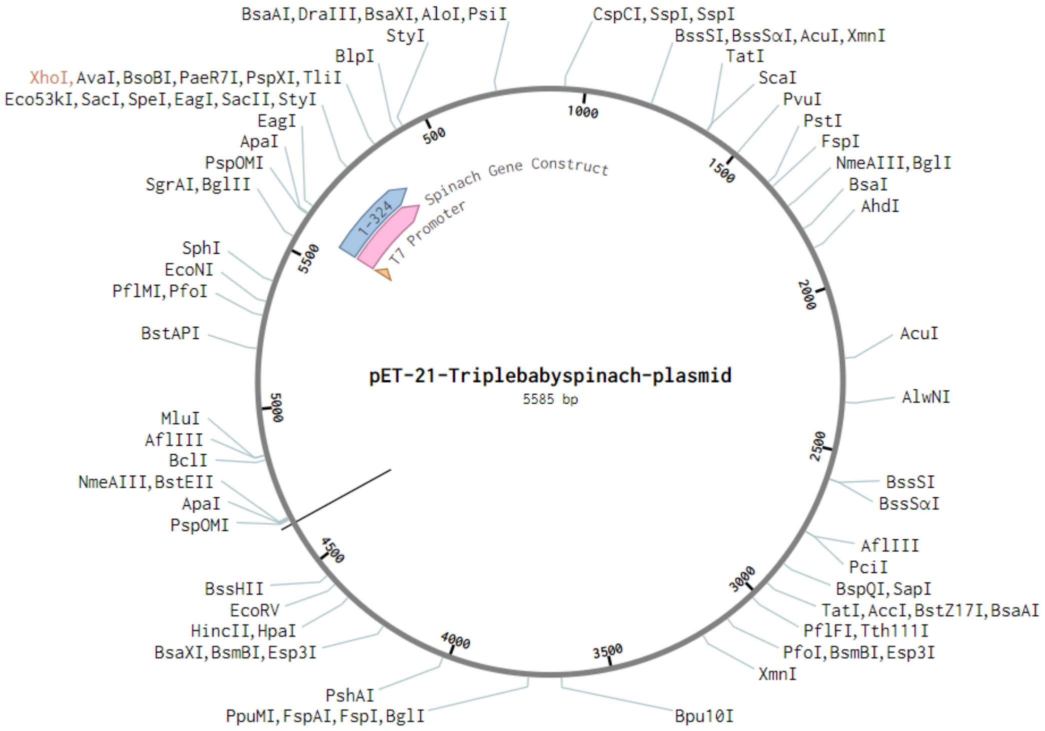


**Figure S6.** Plasmid map of 3xBaby Spinach in a pET-21 vector backbone, co-expressing ampicillin resistance gene and 3xBaby Spinach aptamer.

### References


1. Gaussian 16, Revision B.01, Frisch, M. J.; Trucks, G. W.; Schlegel, H. B.; Scuseria, G. E.; Robb, M. A.; Cheeseman, J. R.; Scalmani, G.; Barone, V.; Petersson, G. A.; Nakatsuji, H.; Li, X.; Caricato, M.; Marenich, A. V.; Bloino, J.; Janesko, B. G.; Gomperts, R.; Mennucci, B.; Hratchian, H. P.; Ortiz, J. V.; Izmaylov, A. F.; Sonnenberg, J. L.; Williams-Young, D.; Ding, F.; Lipparini, F.; Egidi, F.; Goings, J.; Peng, B.; Petrone, A.; Henderson, T.; Ranasinghe, D.; Zakrzewski, V. G.; Gao, J.; Rega, N.; Zheng, G.; Liang, W.; Hada, M.; Ehara, M.; Toyota, K.; Fukuda, R.; Hasegawa, J.; Ishida, M.; Nakajima, T.; Honda, Y.; Kitao, O.; Nakai, H.; Vreven, T.; Throssell, K.; Montgomery, J. A., Jr.; Peralta, J. E.; Ogliaro, F.; Bearpark, M. J.; Heyd, J. J.; Brothers, E. N.; Kudin, K. N.; Staroverov, V. N.; Keith, T. A.; Kobayashi, R.; Normand, J.; Raghavachari, K.; Rendell, A. P.; Burant, J. C.; Iyengar, S. S.; Tomasi, J.; Cossi, M.; Millam, J. M.; Klene, M.; Adamo, C.; Cammi, R.; Ochterski, J. W.; Martin, R. L.; Morokuma, K.; Farkas, O.; Foresman, J. B.; Fox, D. J. Gaussian, Inc., Wallingford CT, 2016.

2. Trott, O.; Olson, A. J. AutoDock Vina: Improving the Speed and Accuracy of Docking with a New Scoring Function, Efficient Optimization, and Multithreading. *J. Comput. Chem.* **2010**, *31*, 455–461.

3. Eberhardt, J.; Santos-Martins, D.; Tillack, A. F.; Forli, S. AutoDock Vina 1.2.0: New Docking Methods, Expanded Force Field, and Python Bindings. *J. Chem. Inf. Model.* **2021**, *61*, 3891–3898.

4. Morris, G. M.; Huey, R.; Lindstrom, W.; Sanner, M. F.; Belew, R. K.; Goodsell, D. S.; Olson, A. J. AutoDock4 and AutoDockTools4: Automated docking with selective receptor flexibility. *J. Comput. Chem.* **2009**, *30*, 2785–2791.

5. Song, W.; Strack, R. L.; Svensen, N.; Jaffrey, S. R. Plug-and-Play Fluorophores Extend the Spectral Properties of Spinach. *J. Am. Chem. Soc.* **2014**, *136* (4), 1198–1201.
